# SAFARI: Pangenome Alignment of Ancient DNA Using Purine/Pyrimidine Encodings

**DOI:** 10.1101/2024.08.12.607489

**Authors:** Joshua Rubin, Jan van Waaij, Louis Kraft, Jouni Sirén, Peter Wad Sackett, Gabriel Renaud

**Affiliations:** Department of Health Technology, Section for Bioinformatics, Technical University of Denmark, Kongens Lyngby, Denmark; UC Santa Cruz Genomics Institute, Santa Cruz, USA

## Abstract

Aligning DNA sequences retrieved from fossils or other paleontological artifacts, referred to as ancient DNA, is particularly challenging due to the short sequence length and chemical damage which creates a specific pattern of substitution (C→T and G→A) in addition to the heightened divergence between the sample and the reference genome thus exacerbating reference bias. This bias can be mitigated by aligning to pangenome graphs to incorporate documented organismic variation, but this approach still suffers from substitution patterns due to chemical damage. We introduce a novel methodology introducing the RYmer index, a variant of the commonly-used minimizer index which represents purines (A,G) and pyrimidines (C,T) as R and Y respectively. This creates an indexing scheme robust to the aforementioned chemical damage. We implemented `SAFARI`, an ancient DNA damage-aware version of the pangenome aligner `vg giraffe` which uses RYmers to rescue alignments containing deaminated seeds. We show that our approach produces more correct alignments from ancient DNA sequences than current approaches while maintaining a tolerable rate of spurious alignments. In addition, we demonstrate that our algorithm improves the estimate of the rate of ancient DNA damage, especially for highly damaged samples. Crucially, we show that this improved alignment can directly translate into better insights gained from the data by showcasing its integration with a number of extant pangenome tools.

## 1 Introduction

The advent of next-generation sequencing (NGS) technologies has enabled high-throughput sequencing of DNA samples in a wide variety of contexts. Typically the samples are from living tissue, leading to easily obtainable high-coverage genomes. However, this is not the case for ancient DNA (aDNA) samples. Ancient DNA has several idiosyncrasies which make it particularly difficult to analyze [34, 40]. First, much of the DNA will typically have been degraded (i.e. fragmented) due to post-mortem enzymatic processes. Moreover, there will typically be contamination from microbial DNA (modern or ancient) which often represents the bulk of the DNA extracted from ancient remains as well as contamination from present-day humans who have handled the sample [17]. This entails that the data will tend to be very low-coverage.

The most characteristic hallmark of ancient DNA, however, results from the hydrolytic cleavage of amino groups from certain nucleobases resulting from long-term exposure to moisture and is known as deamination [34, 7, 46]. The deamination process involves the modification of cytosine (C) nucleobases into uracil (U), which the sequencing machine will interpret as thymine (T) [19]. This therefore results in a pattern of C to T substitutions. When using a double-stranded library preparation is used, these substitutions can occur on the opposite strand thus resulting in G to A substitutions due to base complementarity. When using single-strand library protocols [14, 21], only the C to T substitutions will be observed. Hence, when adenine or thymine is identified in the sequencing data, it represents an ambiguity in the original nucleotide composition, as the base could have originally been either adenine or guanine (A/G), or thymine or cytosine (T/C), respectively. These substitutions complicate mapping to the reference genome. In particular, deamination events in an aDNA sequence may have the unfortunate consequence of forgoing the entire alignment for that fragment, a significant issue we will discuss in more detail.

Up to the present time there have been no widely-used genomic alignment tools specifically tailored for ancient DNA. Instead, the aDNA community has identified various user-defined parameters for standard short-read alignment tools which are typically more permissive and work better for aDNA samples. A number of benchmarking studies have been performed to determine such optimal parameters for aligning ancient data to modern reference genomes. For example, Martiniano [31] *et. al*. find that the more relaxed parameters `-n 0.02 -l 1024 -o 2` work best for `BWA aln` [27], while Xu *et. al*. find that `BWA-MEM` [25] works best for aDNA with parameters `-k 19 -r 2.5` [61], and Poullet *et. al*. find that `--end-to-end` mode is optimal for `Bowtie2` [23] in an aDNA context [38].

Modifying aligner parameters can help the alignment of aDNA samples to some extent but do not mitigate the problem that the non-reference base tends to be underrepresented in aDNA studies [18]. Günther *et al*. showed that this preferential alignment to the reference otherwise called reference bias can impact downstream analyses. Pangenome graphs (also known as genome graphs or variation graphs) are relatively new data structures which have been shown to help to improve the alignment of aDNA [31]. These are graph-theoretic extensions to the standard linear reference genomes that are able to capture more known variation by storing more knowledge about genomic diversity in an organism than is possible with a single genome [13]. More specifically a pangenome graph consists of a set of nodes containing DNA sequences connected by edges, as well as a set of haplotypes (stored separately) each of whose sequence can be reconstructed by walking the graph along a specified route. Crucially, the incorporation of more variation helps mitigate reference bias, and this mitigation has been shown to improve the alignment of both modern and ancient DNA reads [16, 31]. It also has the ability to take the circularity of genomes (such as in mitochondria or chloroplasts) into account by topologically connecting starting and ending nodes. Pangenome graphs are increasingly used for more accurate and informative genomic inference, and methods tailored to pangenome graphs are therefore rapidly being developed and fine-tuned [3, 12]. Notably, the first draft of a human pangenome reference has been released by the Human Pangenome Reference Consortium just last year as of the time of writing [28].

The subcommand `vg map` aligns read kmers against all kmers in the reference graph, including those that do not exist in any constituent haplotypes, which can lead to exponential growth in novel kmers and significant storage and complexity issues. A faster alternative, `vg giraffe`, was developed to map only to kmers traversed by the graph’s embedded walks thus reducing false positives which are then stored using the GBWT or GBZ index [53, 54, 52]. Unlike `vg map`, `vg giraffe` indexes the reference pangenome using minimizers. The minimizer index data structure was first invented to reduce storage space for biological sequence data [43]. Since then, minimizers have proven invaluable as a means of providing promising seeds for potential extension during alignment. Many alignment tools work under the “seed-and-extend” paradigm, in which portions of the input read (seeds) are found to match some hits (i.e. occurrences of these seed sequences) in the reference genome (via a hash table query of the minimizer index). In the extension phase, these hits are clustered and bidirectionally extended, and the most promising of these extensions are retained to produce alignments [2]. The basic idea of the minimizer index is as follows: slide a window across the reference genome. At each step, consider the set of kmers within that window (as well as their reverse complements on the opposite strand). From this set, select the kmer with the smallest hash value. The kmer becomes a key in the minimizer index, and its value stores the set of genomic coordinates where copies of the minimizer in question are to be found as well as a ‘payload’ of data relating to these coordinates [49]. The row of the hash table in which the value is placed is dictated by the hash function being used. The resulting hash table is called the (*w, k*)-minimizer index [43, 47]. The choice of *k* (i.e. the kmer size) is a tradeoff between sensitivity and specificity, with a larger *k* leading to greater specificity and lower sensitivity, and likewise for *w*, which determines the window size and therefore the density of minimizers in the index [43].

In addition to producing graph-specific GAM alignment files, `vg giraffe` can also be used to produce linear BAM files from a graph alignment, despite mapping to the full reference graph. This can be done via the operation of surjection, which lifts pangenome coordinates onto specific walks within the pangenome to produce an `HTSlib`-compatible output [48, 6]. During the procedure of surjection from graph alignments to a particular haplotype, regions of the alignment traversing nodes which are supported by that haplotype are left unchanged, while alignment to nodes unsupported by that haplotype are carried over to the nearest supported node, with appropriate edits added to affected node alignments [15]. Since at the time of writing the majority of aDNA researchers do not work with pangenomes, this feature allows the benefits of a pangenome aligner to be more widely useful across the aDNA community, even if they are accustomed to using downstream tools that expect linear BAM files. Indeed, mapping to a pangenome followed by surjecting to a linear reference can lead to improved alignments compared to directly mapping to the linear reference. (Please see Supplementary Section D.6 for more details).

A number of well-established full-genome alignment tools index the reference using minimizers. Among linear genome aligners, `minimap2` [26] uses minimizer sketches to quickly identify candidate alignment regions, thereby reducing the computational time required for sequence alignment, and `kraken2` [60] creates a database of minimizers from reference genomes to rapidly identify the taxonomic origin of sequence reads from a metagenomic sample. In the context of pangenome alignment, `SPUMONI2` uses minimizers to reduce the set of reads as well as the reference graph before computing matching statistics for multi-class classification [1], and `vg giraffe` [54] makes use of minimizers from the graph’s embedded haplotypes to align to all haplotypes simultaneously.

For typical alignment tools employing minimizers, while the entire alignment need not be an exact match to the reference genome, some portion of it (the seed) will need to match exactly, due to the use of hash tables for constant-time lookup of seeds in the index file. This heuristic is problematic in the case of ancient DNA (aDNA) where potential seeds may well contain deaminated bases and therefore these reads cannot be aligned. Since aDNA samples are very often low-coverage in addition to being chemically damaged and highly fragmented, loss of information is incurred when potentially non-spurious alignments are discarded due to deamination events. A more aDNA-tailored indexing scheme designed to be robust to deamination events due to aDNA damage is therefore clearly indicated. It is also worth noting that this issue also generalizes to tools which may not use minimizers or other preselected subsets of indexed kmers, but which still require exact seed matches for lookup (e.g. `BWA aln` [27]).

We introduce a modified version of the `vg giraffe` subcommand called `SAFARI` (Sensitive Alignments From A RYmer Index), available at https://github.com/grenaud/SAFARI which implements a novel damage-aware algorithm to align aDNA sequences to pangenomes. We leverage the fact that aDNA damage will result in transitions from purine to purine or pyrimidine to pyrimidine. We use kmer encodings in purine/pyrimidine space called RYmers which are impervious to aDNA damage. `SAFARI` can recover more alignments than `vg giraffe` in the presence of aDNA damage, and the increased number of non-spurious alignments is proportional to the rate of damage in the sample. In addition to the standard input for `vg giraffe` (namely the input FASTQ file(s), the reference graph, and the relevant index files), `SAFARI` takes as input an additional ancillary index file called the RYmer index (file extension: *.ry) which supplements the standard minimizer index, allowing `vg giraffe` to benefit from our novel RYmer encoding procedure. As the program computes the probability of a false RYmer match, `SAFARI` requires the user to provide estimated per-position damage rates. These estimates can be misspecified to a certain degree without undue adverse effect on the program. `SAFARI` also incorporates significant modifications to the `vg giraffe` alignment logic (pre-extension) as detailed in Section 2.

We will demonstrate the improved performance on aDNA samples both by directly comparing alignments produced between the two tools on simulated datasets at varying levels of damage, as well as by demonstrating that the alignments lead to improved downstream performance of pangenome tools which make use of `SAFARI` as compared to the standard `vg giraffe`.

## 2 Methods

### 2.1 RYmer Index

Here we introduce the first aDNA-tailored variant of the minimizer index, which we shall call a RYmer index. We define a RYmer to be a sequence obtained from a kmer by replacing each occurrence of G and A with R (the IUPAC encoding of a purine), and each occurrence of C and T with Y (the IUPAC encoding for a pyrimidine) [10]. For example, GATC becomes RRYY or CCCC and TTTT both become YYYY. This entails that deaminated bases will generate the same RYmer regardless of the rate of damage. Note that this encoding is a surjective and it entails that the RYmer index will be no larger than, and usually smaller than, the minimizer index. This is because two distinct minimizers may have the same RYmer encoding.

The notion of RYmers is not entirely new to this paper. Purine/pyrimidine encodings have been used in the context of DNA substitution models for phylogenetic analyses, for instance, [37, 59]. However, as far as we are aware, RYmers have never been used in the context of genomic alignment or indexing. RYmers make it possible to recover seeds that do not match exactly in nucleotide space due to deamination events, thereby rescuing these alignments for downstream processing. However, the cost is loss of information. Supposing that a RYmer has size *k*, then its (Shannon) information content (a measure of how much one can learn from the sequence) is exactly half that of a kmer of the same size, assuming all bases are equiprobable. More specifically, there are 4^*k*^ possible kmers. The Shannon information content of a kmer is log_2_(4^*k*^) = 2*k* bits. By similar reasoning, the Shannon information content of a RYmer of length *k* is log_2_(2^*k*^) = *k* bits. The ratio of the Shannon information content of kmers to RYmers is then 2*k/k* = 2. In other words, a RYmer will need to be precisely twice as long as a kmer to convey the same amount of information (which is readily seen by the fact that it conveys the information about whether a certain base is a pyrimidine or a purine, but not whether it is C or T, or G or A, respectively).

### 2.2 RYmer Index Hash Table Structure in SAFARI

Our RYmer index hash table structure in `SAFARI` is similar in structure to the standard `vg giraffe` minimizer index, with two crucial differences. In the `vg giraffe` framework, as with many other programs, the minimizer index constitutes a hash table where each key is 64 bits in length. Of these, 62 bits are dedicated to encoding the minimizer sequence. The value comprises a hit (if the minimizer instance is unique in the reference, in which case it is 192 bits) or a pointer to a vector of hits (if not, in which case it is 64 bits). 64 bits per hit are devoted to storing node identifier, orientation, and sequence offset on the node. The remaining 128 bits constitute the payload which stores arbitrary information related to the graph position [54, 49]. A hash function (in the specific case of `vg giraffe` this is an integer hash function described by Thomas Wang [57]) is used to map keys to rows in the hash table where the values are stored.

In the minimizer index the standard encoding of minimizers in `vg giraffe` is accomplished using a straightforward binary representation of the DNA nucleobases:

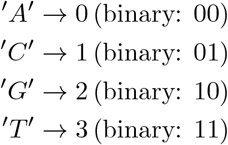

Strings of multiple nucleobases are concatenated together in their binary representation, forming a unique 62-bit key for every minimizer (up to a length of 31 bases).

The RYmer encoding of a sequence, in contrast, requires only one bit per nucleobase:

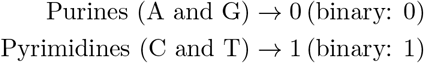

This is the first distinction between the two indexes, namely that the key stores the RYmer encoding of the minimizer.

The second distinction is the value of each hit. In the minimizer index, this is the pair of position and payload mentioned above. However, in the RYmer index we appropriate the payload to instead store the bitstring corresponding to the minimizer that gave rise to the RYmer hit in question. This is crucial for the `SAFARI` algorithm, firstly because it is in general impossible to recover a kmer sequence from its RYmer encoding due to the information loss, but also because it facilitates rapid, constant-time retrieval (as opposed to searching over all keys in the minimizer index, for example, which would be linear in the number of minimizers — a number which is typically very large). Figure 2 illustrates the hash table structures for each index.

**Figure 1:**
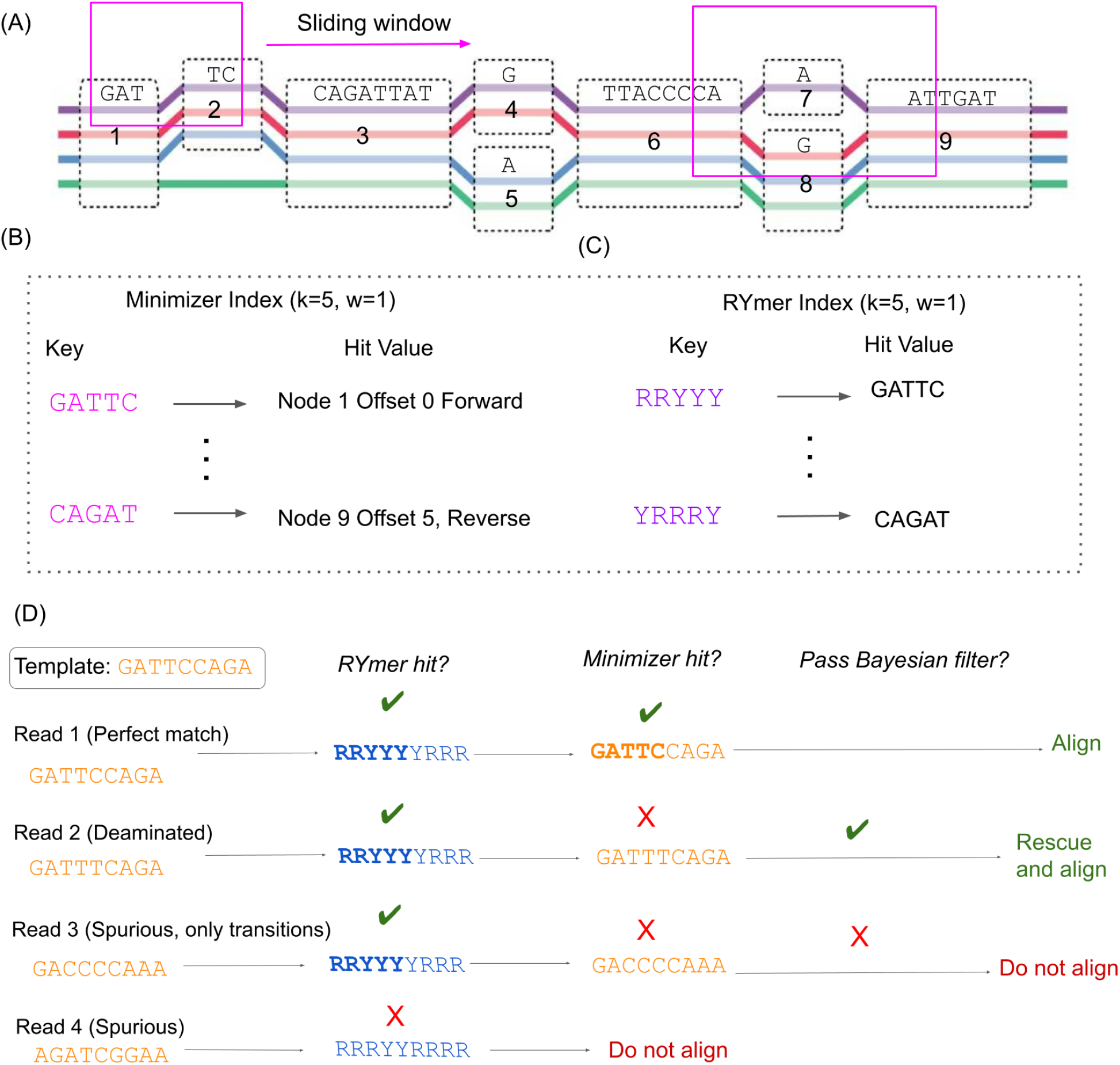
**(A)** A sliding window traverses the pangenome reference graph. The purple, red, blue, and green paths along the graph are the embedded haplotypes. Within each window, the hash value of all kmers of the graph’s embedded haplotypes, as well as their reverse complements, are considered. **(B)** The integer encoding of the kmer with the minimum hash value for a given window is placed as a key in the minimizer index, with the 64-bit value of this key (the payload) being the locations of all copies of this kmer in the graph, as well as associated metadata. The RYmer index is constructed in an analogous way to the minimizer index, but the sequence of the key is transformed into RYmer space, and the value stores the corresponding minimizer key that gave rise to this RYmer. Four situations might arise from a given template sequence containing a seed match to an indexed RYmer. **(i)** The match is perfect, in which case we will observe the seed match in RYmer space, and also kmer space. The read is immediately aligned. **(ii)** A base in the seed is deaminated (*C* → *T*), so we observe a match in RYmer space but not kmer space. This read is passed through our Bayesian filter where the posterior probability of being a true positive is computed. Supposing this posterior is above our predefined threshold, the read will be rescued. **(iii)** A spurious RYmer space match in which all mismatches in kmer space are transitions. Since the match in kmer space is not exact we apply our Bayesian filter, which does not let the read through. **(iv)** A spurious read which does not match in either index. It is not even considered for alignment.

**Figure 2:**
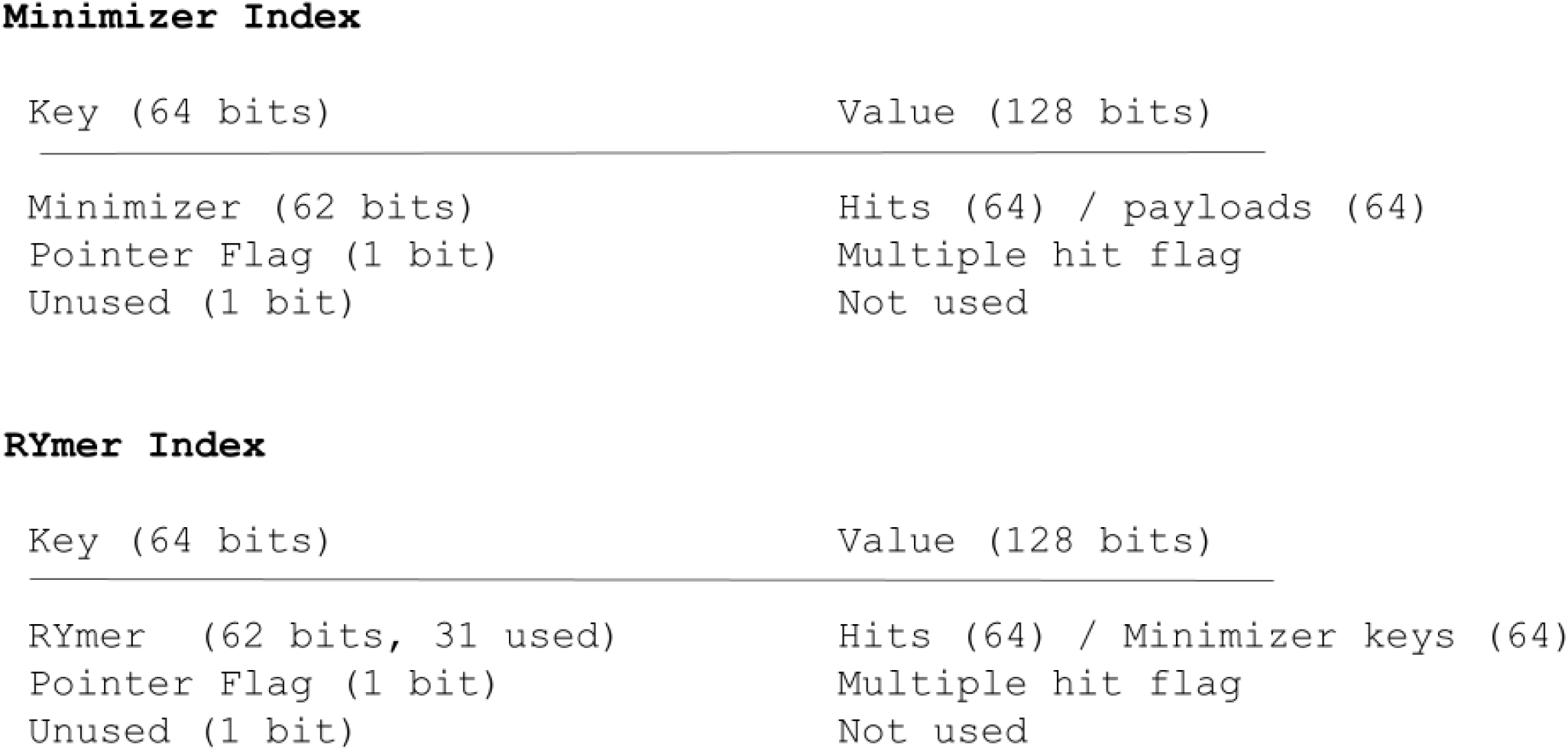
In the case of the RYmer index, the portion of the hash table value devoted to the payload is instead used to store the key of the corresponding minimizer that gave rise to that RYmer (or a pointer to all keys of minimizers which gave rise to that RYmer, if not unique), allowing for efficient, constant-time lookup during runtime.

It is noteworthy that due to the halving of the information content of a RYmer string, it is possible to encode RYmers of double the length of the largest permissible kmer (e.g., a RYmer of 62 bases under the `giraffe` encoding scheme).

### 2.3 Haplotype Indexing Implementation and Details

To implement the aforementioned RYmer index we introduce a new `vg` subcommand, `vg rymer`, which functions analogously to the standard `vg minimizer` subcommand of the `vg` toolkit in indexing the graph’s embedded haplotypes. Users define a desired *w* parameter (i.e. number of consecutive kmers per window on a single strand) and kmer size *k*. The subcommand then indexes the reference haplotypes using these parameters and serializes the resulting RYmer hash table to disk. The subcommand code is found here: https://github.com/grenaud/SAFARI/blob/master/src/subcommand/rymermain.cpp.

In `vg giraffe`, minimizers per window are selected for indexing with the help of a circular buffer data structure of fixed size 2^*i*^ (where *i* is the smallest number for which 2^*i*^ ≥ *w* and *w* is the number of consecutive kmers per window on a single strand) [49]. During indexing the circular buffer advances, which represents the sliding of the window across one of the pangenome graph’s embedded haplotypes. This is done in both the forward and the reverse complement orientations. Every time the window slides (i.e. the circular buffer advances), one kmer from the previous window is dropped. The forward and reverse complement of each candidate kmer is hashed, and the candidate with the smallest hash value is included (kmers containing ambiguous nucleobases are never indexed). In this way all minimizers are selected in turn and placed in the minimizer (*.min) index. The procedure is repeated for every haplotype in the graph [50, 51].

The logic for RYmers is similar but not analogous. We require that the RYmer indexing parameters are identical to those used in the corresponding minimizer index. Under these parameters we use the same sliding window approach to selecting RYmers for indexing in the RYmer (*.ry) index file. However, we wish to ensure that for every kmer in our minimizer index, the corresponding RYmer exists in the RYmer index as well. Therefore we do not index RYmers by finding those with minimum hash values in the transformed sequence space, as this RYmer set can be different from the minimizer set. We instead take the minimizer set and index their corresponding RYmers (for pseudocode on how to go from a minimizer key to a RYmer key please see Supplementary Section H.1). The second difference is that we overwrite the first unsigned integer in the kmer value payload (which comprises two unsigned 64-bit integers in total) to store instead the key of the corresponding kmer to the RYmer in question from the minimizer index for later lookup. This is possible because we never use the RYmer value payload for any other purpose, as explained in Section 2.4.1.

### 2.4 Inference Model Details

### 2.4.1 SAFARI Bayesian Filter Model

We consider here the problem of quantifying the degree of trust which should be given to an alignment in RYmer space. This quantification is based on the comparison of the seed (in the read sequence) and the hit (in the index) in the original kmer space (given that we know an exact match has occurred in RYmer space).

In the case we have a match between seed and hit in kmer space, we always allow the gapless haplotype-consistent extension to proceed, just as in standard `vg giraffe`. In this case, no deamination event has occurred in the seed. Therefore the only interesting case is when we have a seed whose sequence matches a hit in RYmer space, but the corresponding kmer sequences do not match. In our simplified model there are only two possible explanations for this: i) alignment is correct but a deamination event occurred in the seed, or ii) the alignment is spurious. For the latter case, there are other explanations such as sequencing errors or private mutations in the sample with respect to the reference, which we do not model as we seek to rescue alignments that would have been skipped due to deamination.

Let *M* denote the event that the sole explanation for the RYmer mismatch is due to deamination. In other words, the kmer would have matched had it not been for deamination events. For a read *r*, our aim is to compute the posterior probability *P* (*M* | *r*) given by:

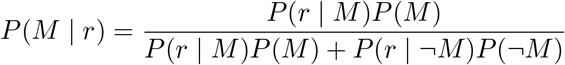

The probability of observing the read *r* given that the alignment was spurious *P* (*r*| *¬M*) depends on the nature of any potential contaminant genomes in the given sample. We mean by spurious that the RYmer match was due to incidental agreement rather than identity followed by damage events. To obtain a reasonable likelihood function, we therefore ran some *in silico* simulations of bacterial reads mapped to the human mitogenome to obtain a reasonable model. The exact details can be found in Supplementary Section E.2. The results roughly suggested a truncated power-law distribution for number of mismatches in kmer space given an exact match in RYmer space. Mismatches in the two encodings can occur anywhere in the kmer with equal probability, leading us to a binomial PMF for the spurious model likelihood function:

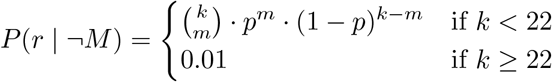

Where the probability of mismatch *p* = *ak*^*b*^, *a* = 1.0014, *b* = − 0.6628 (the best-fit parameter values (via the method of least squares) from the *in silico* simulations), and *m* is the observed number of mismatches. Note that after *k* = 22 the mismatch proportion in the seeds tends to taper off, making it difficult to model precisely. To mitigate false positives, we therefore impose a constant probability for this kmer size. It is important to note that this spurious RYmer space match model is only a cursory approximation, and we expect the mismatch distribution to be heavily dependent on the nature of the contaminant genomes. More details about the bacterial simulations and spurious mapping model can be found in Supplementary Section E.2.

It remains to find a model for *P* (*r*|*M*). For simplicity, we assume that each observed nucleobase *b* is independent:

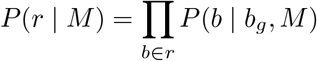

where *P* (*b*|*M*) will depend on the probability of deamination for the given position along the fragment according to the damage profile for the input sample, which must be provided by the user via the `--deam-3p` and `--deam-5p` flags, as dictated by a commonly-used model from `ROHan` [42]. (We note in passing that misspecification of these initial estimates can adversely effect performance, but generally not unduly, as shown in Supplementary Section D.5). We also emphasize that the assumption of independence is not strictly true in general, but modeling such higher-order dependencies would involve several factors (e.g. measures of linkage disequilibrium, dinucletotide frequencies, etc.) which would be cumbersome and largely unnecessary for present purposes.

Let the base of the reference graph be denoted *b*_*g*_. Then we have:

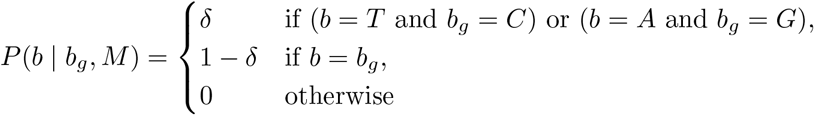

where *δ*depends on the type of substitution and position on the aDNA fragment. The implementation for the above is taken from Renaud *et. al*. [42]. The one difference is that we do not account for sequencing errors as our goal is not to make `vg giraffe` more sensitive to sequencing errors but rather impervious to deamination.

The user-defined posterior probability of non-spurious alignment threshold (the flag “-j” in `SAFARI`) constitutes a threshold on *P* (*M*|*r*) to determine whether we allow the RYmer in question to proceed to the extension phase. The prior on an alignment being non-spurious as opposed to spurious (conditioned on a RYmer space match) is also provided by the user via the “-V” flag, with default value 0.5 (i.e. by default both models are equiprobable). Importantly, we note that a RYmer passing this filter does not necessarily mean it will produce an alignment — if too many mismatches are encountered during extension the alignment will fail nonetheless, for example, providing another avenue of filtering for spurious RYmer matches. Further comments about this threshold and other user-defined parameters in `SAFARI` can be found in Supplementary Section E.1.

In the case of the RYmer passing our Bayesian filter, the next step is to convert it into a corresponding minimizer. As previously mentioned, we use the RYmer index payload (which is never used to locate GBWT graph coordinates unlike its minimizer counterpart) to instead store the key in the minimizer index of the minimizer(s) from which the RYmer in question was derived. From this key the corresponding minimizer object is obtained using a preexisting function. By doing this, for a given RYmer at runtime we can locate the corresponding minimizer(s) key in constant *O*(1) time. If the converted minimizer is a duplicate of one already found in the standard kmer space, the duplicate is removed.

### 2.5 Alignment Code Modifications

#### 2.5.1 Bayesian Filter Implementation

To apply the Bayesian spurious alignment filter described in Section 2.4.1, we first consider both the set of kmers and RYmers for the input data as discussed above. The set of RYmers are then passed through a custom lambda function which implements the filtering logic, and those that do not pass the filter are removed from downstream processing. The RYmers that do pass the filter are converted back to their minimizer equivalents by accessing the RYmer hash table value, which stores the corresponding minimizer key. These rescued minimizers are then aggregated with the standard minimizer set. The resulting collection of augmented minimizers is then passed to the seed-finding function which proceeds just as usual in `vg giraffe`. Importantly, since `SAFARI` effectively only rescues seeds that would have been discarded due to deamination, we note that downstream processing remains unaffected by the modification. For instance, the clustering and extension steps proceed exactly as usual. The resulting alignment (GAM) file from `SAFARI` will correctly report the edits needed to go from reported mappings to the reference sequence. Since everything downstream of seed finding (e.g. clustering and extension) is still done in the standard kmer space, mapping qualities from `SAFARI` should remain trustworthy, although it is true that in theory they will be very slightly misspecified due to the lack of a model for falsely rescued seeds. Please see Supplementary Section H.2 for high-level pseudocode of the RYmer Bayesian filter.

## 3 Results

We used four benchmarking strategies to demonstrate the efficacy of `SAFARI` for rescuing deaminated seeds in ancient DNA samples. (i) We compare `SAFARI` to `vg giraffe` with regard to alignment metrics on the surjected linear (i.e. one genome) BAM files, and show that the degree of improvement of these alignments is commensurate with the level of damage. (ii) We then compare damage rate matrix estimates obtained from these surjected BAM files, and show that damage rate estimates improve with `SAFARI` due largely to survivor bias mitigation. (iii) After this we compare `SAFARI` to `vg giraffe` as the internal aligner used in the program `HaploCart` [44], a subcommand of the `vgan` suite of pangenome tools which calls human mtDNA haplogroups, and show that the rescue of alignments with deaminated seeds helps to improve `HaploCart`’s performance on aDNA samples. (iv) Finally, we do the same for `euka` [56], another subcommand of `vgan` which characterizes the eukaryotic composition of an environmental aDNA sample. We show that using `SAFARI` as the internal aligner improves the performance of `euka` as measured by number of detected fragments per taxon present in the sample.

### 3.1 Surjected Linear Alignments

We compared `vg giraffe` and `SAFARI` alignments from *in silico* simulations to a reference graph comprised solely of the Revised Cambridge Reference Sequence (rCRS) which serves as the human mitochondrial reference (NCBI accession: NC 012920.1, H2a2a1 haplogroup in Phylotree build 17) [55, 4]. This is to demonstrate how our RYmer index can be exploited to improve aDNA inference via the rescue of deaminated seeds even when integrating with a pipeline that does not make use of pangenome graphs throughout. We experimented with a wide number of distinct minimizer and RYmer indexing parameters, namely the minimizer length *k* and number of consecutive kmers per window per strand *w* (enumerated in Supplementary Table 6). We conducted the experiment using two separate empirically-derived fragment length distributions, the first from Chagyrskaya Cave in Siberia [30], and the second from Vindija Cave in Croatia [39]. These two fragment length distributions are visualized in Supplementary Figure 1. The simulations try to emulate empirical mitochondrial aDNA fragments contaminated with both bacteria and NuMTs (i.e. nuclear pseudogenes of mitochondrial origin) [5]. See Supplementary Section F.2 for more details. In the case of `SAFARI`, the input damage matrices are those estimated for the given sample by aligning using `BWA aln` (with parameters “-n 0.01 -o 2 -l 16500”) and calculated using the damage profiler `bam2prof` [41]. For comparison to standard (i.e. non-graphical) aligners, we also included in the experiment a number of other linear short read alignment tools, namely `SHRiMP` [45], `BBMap` [8], `BWA aln` (both default and with aDNA-specific parameter settings) [27, 32], `BWA-MEM` [25], and `Bowtie2` [23] with the more sensitive `--end-to-end --sensitive` flags as recommended for aDNA samples in [38]. `SHRiMP` is quite a sensitive alignment tool for aDNA samples but is unfortunately not maintained since 2014 [9]. Full details about sample generation and exact commands for this experiment are provided in the Supplementary Section F.2.1. We stratify over four distinct damage patterns when generating samples, namely zero damage, single-stranded damage, double-stranded medium-level damage, and double-stranded high-level damage. Examples of each are visualized in Supplementary Figures 2 and 2, which was generated using a script from the `bam2prof` Github repository (https://github.com/grenaud/bam2prof/blob/master/src/plotbam2prof.py).

Since reads can either be aligned or unaligned, and since the alignment is generally either clearly correct or clearly incorrect, it is sensible to treat this as a binary classification task. Reads from the mitochondrion are considered positives. A mitochondrial read is a true positive if it maps to within 50bp of the correct location in the mitogenome, a false positive otherwise. Bacterial and NuMT reads are considered negatives. They are true negatives if they are unmapped, false negatives if they are mapped.

In this context, we can define the metrics of sensitivity, specificity, and *F*_1_ score as follows:

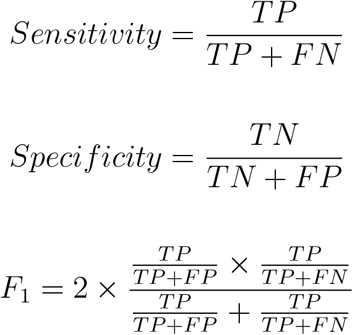

Loosely speaking, sensitivity captures the ability to map reads which should be mapped, while specificity captures the ability to avoid mapping reads which should not be mapped. We can think of the *F*_1_ score as providing a balanced measure taking both specificity and sensitivity into account.

Oftentimes in an aDNA setting, specificity is the priority due to sparsity of data, but mitigating spurious alignments is also very important. Figure 3 plots sensitivity vs. specificity by pangenome tool (`SAFARI` or `vg giraffe`) for double-stranded high-damage and zero-damage samples, respectively, on samples following the Vindija-derived fragment length distribution. Notably, the alignments are exactly the same in the absence of damage, while we can see a distinct improvement in sensitivity for `SAFARI` — with a concomitant slight drop in specificity — for the high damage samples. However, the plot shows that regardless of the minimizer length and window size, `SAFARI` has higher sensitivity. For example, with the samples described in Table 1 at the sensitivity-optimized (*k, w*) = (10, 2) parameters we recover 102 more non-spurious alignments (from 269425 for `vg giraffe` to 269527 for `SAFARI`, out of out of 270163 total). This increase may seem modest, but that is only because sensitivity is near optimal already when the kmer size is this small. At default kmer length (*k* = 29) the number of recovered alignments for the same dataset jumps to 14883 (from 123588 for `vg giraffe` to 138471 for `SAFARI`, again out of out of 270163 total). The reader may wonder why we would not always use smaller kmer lengths — on mitochondria it seems clear that smaller kmer lengths are indeed preferable, but a short *k* will have its disadvantages in terms of runtime and spurious alignments which will be particularly evident for larger reference genomes.

**Table 1:** Raw counts of mapped reads by provenance in the surjected linear alignment experiment for the dataset of *in silico* simulations following the empirically-derived fragment length distribution from Vindija Cave, Croatia under high damage rates. *NuMT* is the number of mapped NuMT reads. For readability, only samples for a single subsampling rate (0.9) are shown. Counts are reported as aligned/total. Note that *k* = 29, *w* = 11 are the default parameters. For more data at different indexing parameter pairs please see Supplementary Table 10.

| program | $k$ | $w$ | Bacterial | NuMTs | Mitochondrial |
| --- | --- | --- | --- | --- | --- |
| vg giraffe | 10 | 2 | 36/90235 | 88/360 | 269425/270163 |
| SAFARI | 10 | 2 | 51/90235 | 89/360 | 269527/270163 |
| vg giraffe | 12 | 3 | 3/90235 | 77/360 | 268211/270163 |
| SAFARI | 12 | 3 | 3/90235 | 80/360 | 268526/270163 |
| vg giraffe | 15 | 5 | 1/90235 | 54/360 | 260175/270163 |
| SAFARI | 15 | 5 | 1/90235 | 54/360 | 260773/270163 |
| vg giraffe | 29 | 11 | 0/90235 | 5/360 | 123588/270163 |
| SAFARI | 29 | 11 | 0/90235 | 8/360 | 138471/270163 |
| SHRiMP |  |  | 0/90235 | 86/360 | 269223/270163 |
| BWA-MEM |  |  | 0/90235 | 27/360 | 237299/270163 |
| BWA aln |  |  | 0/90235 | 17/360 | 228817/270163 |
| BWA aln (anc) |  |  | 0/90235 | 19/360 | 251049/270163 |
| Bowtie2 |  |  | 0/90235 | 15/360 | 206858/270163 |

**Figure 3:**
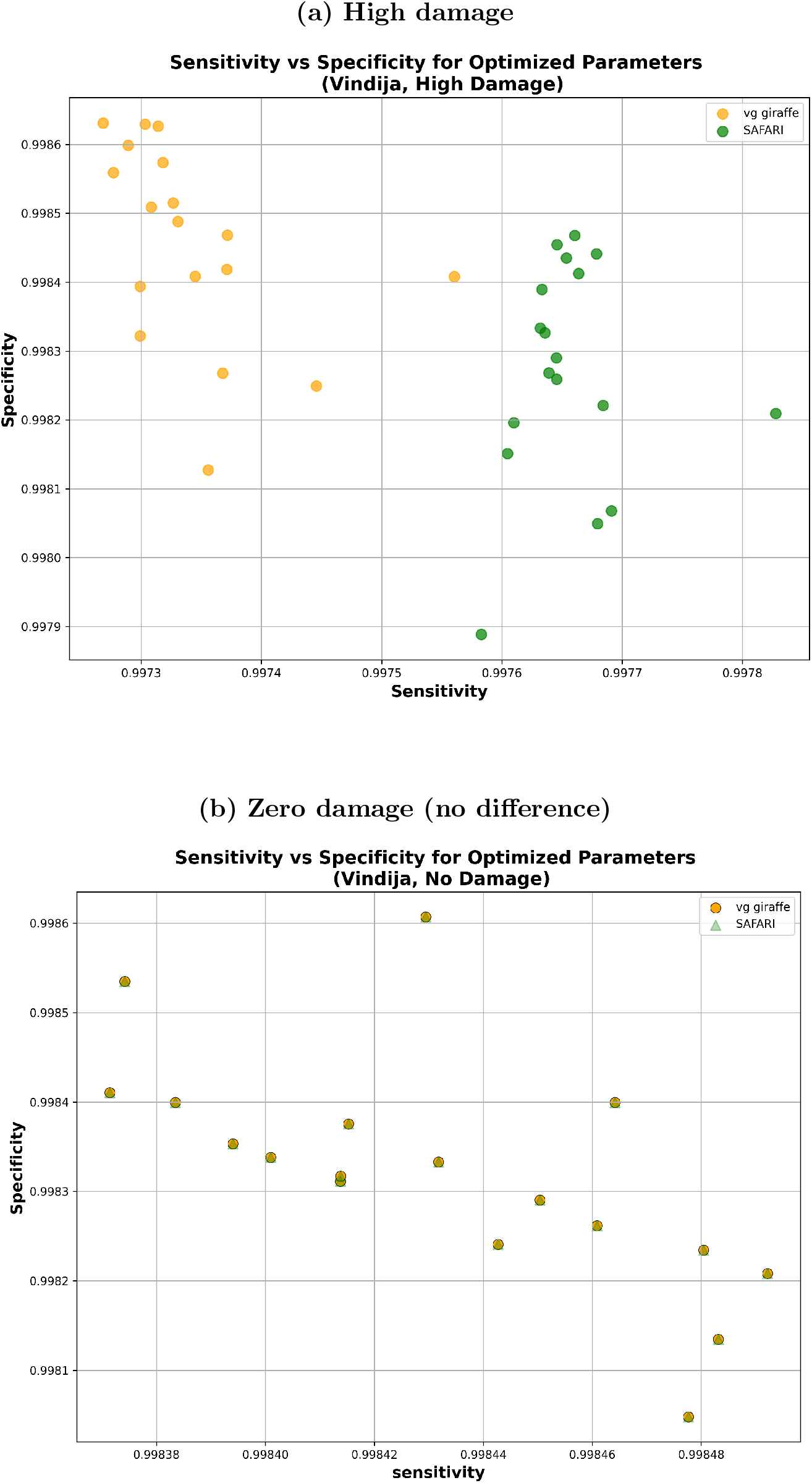
Sensitivity vs. specificity comparison of `SAFARI` and `vg giraffe` on surjected linear alignments using the empirically derived fragment length distribution from Vindija Cave, Croatia. The minimizer length *k* and number of consecutive kmers per window per strand *w* were individually selected to optimize median sensitivity across samples out of all 26 tested parameter pairs. In both cases, the optimized indexing parameter values were *k* = 10, *w* = 2. (a) Samples with high levels of damage. Here we can see that sensitivity has clearly improved with `SAFARI` thanks to the recovered alignments, at the expense of a slight drop in specificity. This is almost always a beneficial tradeoff in the context of aDNA analysis. (b) Samples with zero damage. As expected, there is no differen1c3e between `SAFARI` and `vg giraffe`, as the resulting alignments are identical thus showing that our strategy does not generate spurious alignment in the absence of deamination.

The juxtaposition of results for high damage and zero damage showcase a beneficial tradeoff conferred by the RYmer indexing strategy for aDNA samples, allowing for more non-spurious alignments to be recovered in the presence of damage in the seed. We obtained similar results using the Chagyrskaya Cave distribution dataset; please see Supplementary Section 6.

To further illustrate the point that the damaged seed recovery is most clearly beneficial when damage is at its highest, we show in Figure 4 the median *F*_1_ score for the Vindija Cave distribution dataset for each of the two pangenome aligners, stratified by damage. It is clear that by rescuing deaminated seeds, `SAFARI` improves overall alignment performance, especially on highly damaged samples. The analogous plot showing similar results for the Chagyrskaya Cave distribution dataset can be found in Supplementary Section G.3.3.

**Figure 4:**
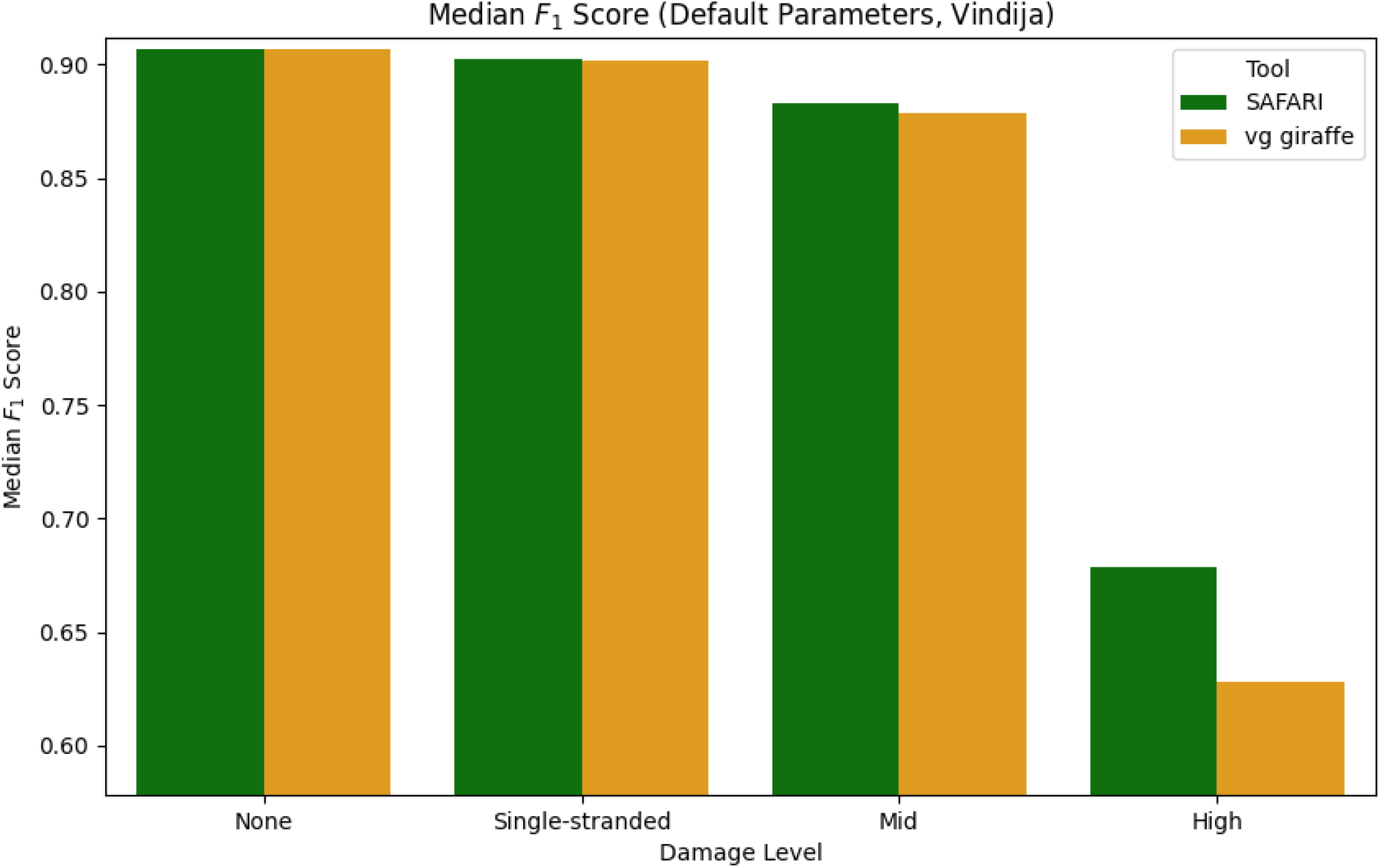
Median *F*_1_ score of each of the two pangenomic alignment tools stratified by level of aDNA damage under default indexing parameters (*k, w*) = (29, 11) for the empirically derived fragment length distribution from Vindija Cave, Croatia. The improvement in median *F*_1_ score conferred by the deaminated seed rescue in `SAFARI` is especially clear at high levels of damage.

The results thus far indicate that `SAFARI` improves upon `vg giraffe` for high-damaged samples. We next sought to determine how these pangenome aligners fare compared to commonly used linear aligners for ancient DNA. To provide such a comparison we show in Figure 5 the median *F*_1_ scores for all tools evaluated in this experiment, again stratified by damage, for the Chagyrskaya Cave distribution dataset. For the pangenome aligners `vg giraffe` and `SAFARI`, the indexing parameters were selected to optimize for median sensitivity. We see that, as expected, all tools perform equally well on zero or low-damage samples. However at medium levels of damage and far more pronouncedly at high levels of damage, only three tools continue to perform well, namely the two pangenome aligners, and also `SHRiMP` (which is no longer being maintained). `SAFARI` narrowly edges out the other tools at high damage with respect to median *F*_1_. This demonstrates that the performance of pangenome aligners such as `SAFARI` is superior to currently maintained linear alignment tools.

**Figure 5:**
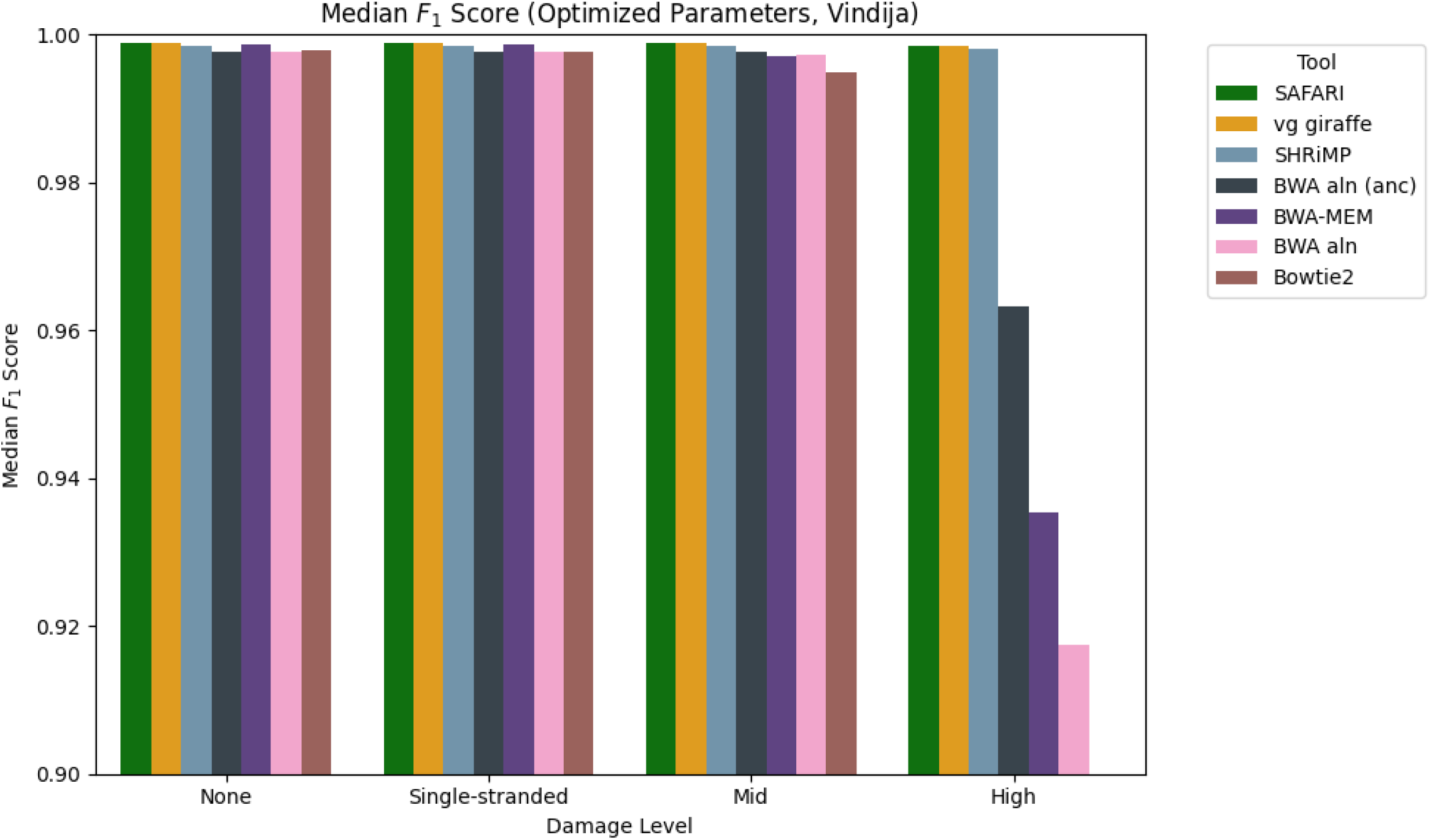
Median *F*_1_ score across samples obeying the fragment length distribution from Vindija Cave, stratified by level of damage. In the case of `SAFARI` and `vg giraffe` we use indexing parameter values (i.e. kmer size *k* and consecutive kmers per forward-strand window *w*) that optimize for maximum sensitivity. The optimized values were (*k, w*) = (10, 2). We see that many tools worsen in performance precipitously once damage reaches a high level, with the two pangenome tools and `SHRiMP` being the three that are relatively robust to high damage overall. For the analogous plot with samples obeying the fragment length distribution from Chagyrskaya Cave please see Supplementary Section 11.

Reporting summary statistics of performance (e.g. sensitivity, specificity, *F*_1_) is informative, but it is important to report the raw counts for the data as well. Below (Table 1) we provide raw counts for aligned reads by provenance (endogenous mitochondria DNA, NuMTs, or bacterial DNA) by pangenome aligner for the Vindija Cave empirical fragment length distribution *in silico* simulation dataset.

#### 3.1.1 Damage Profile Estimation

Accurate estimation of the deamination rate in ancient DNA samples is critical because deamination patterns are one of the signature litmus tests for authentication. However, survivor bias during mapping can lead to systematic underestimation of damage in an aDNA sample. By rescuing and aligning reads affected by deamination events `SAFARI` mitigates survivor bias towards less damaged material in the sample, thereby allowing for more accurate damage rate estimates. To assess the extent of improvement in damage rate estimates for `SAFARI` as compared to `vg giraffe` we looked at the median root-mean-squared error (RMSE) between the ground truth and estimated damage rate matrices for our surjected linear alignment data. Naturally, we expect this quantity to decrease as the alignments improve. The median RMSE was chosen as the metric because it is more robust to outliers than the mean, although the mean RMSE results are very similar and would not affect the main takeaway. We stratified over four different damage rate matrices: zero damage, double-stranded mid-level damage, double-stranded high-level damage, and single-stranded low-level damage (e.g. damage patterns found when using a Santa Cruz library preparation) [21]. Details on these nucleotide misincorporation matrices are provided in the Supplementary Section J.1. We found that there is zero difference when damage is not present in the samples, and very little change when low levels of damage are present (i.e. single-stranded libraries), but at medium and especially high levels of damage the improvement is especially manifest for both fragment length distributions. No doubt the improvement for damaged samples is mainly driven by the greater number of alignments available when using `SAFARI` due to the recovery of reads with deaminated seeds. Full data from this experiment is provided in Supplementary Section G.3.8.

### 3.2 HaploCart with SAFARI

`HaploCart` is a subcommand of the `vgan` suite of tools for panmitogenomics which calls single-source human mtDNA haplotypes using alignment with `vg giraffe` to the first published human mitochondrial pangenome reference graph [44]. The tool uses Bayesian modeling to estimate the maximum likelihood haplotype of a (possibly ancient) sample. In the original `HaploCart` paper, we assessed the performance of the tool on 20 ancient samples despite the fact that we did not have an explicit model for aDNA damage. We found that our tool still outperformed other mtDNA haplogroup classifiers. Nonetheless, the need for a mapper specifically designed for aDNA to pangenomes was clearly indicated. We reran our analysis on these same samples using the RYmer-corrected `HaploCart` with the original reference graph to determine if the performance of the program would improve due to the additional alignments. (Please refer to Supplementary Section F.1 for details). We used high-level damage rate matrices as initial estimates based on the samples’ antiquity. We downsampled to four target coverage depths, namely 0.25X, 0.5X, 1.0X, 2.0X. To account for stochastic effects at low coverage depths, we generated five replicates per sample. We use indexing parameters (*k, w*) = (10, 2) for both the minimizer and RYmer index, as these were the parameter values that we found to optimize for maximum sensitivity for both tools in the linear alignment experiment. Both `HaploCart` versions were run with the same mapping parameters, for all parameters used by both tools. In the case of `SAFARI` we used as input the high-level damage matrices specified in Supplementary Section J.1.

We compared predicted haplogroups to ground truth across samples, using Levenshtein (edit) distance as our metric [24]. Note that Levenshtein distance between haplotype sequences is more meaningful than shortest path distance on the mitochondrial tree because of the possibility of back mutations, and also because named haplotypes can differ from one another by varying numbers of polymorphisms. Ground truth labels were determined by running both tools at full coverage. In the original article, we compared our tool against `HaploGrep2`, one of the most used tools for assigning mitochondrial haplotypes [58]. In certain cases the prediction for `HaploGrep2` disagreed with `HaploCart` and instead reverted to the haplogroup H2a2a1 of the Revised Cambridge Reference Sequence (rCRS), which was used as a reference [4]. This is a reference bias issue idiosyncratic to `HaploGrep2`, and so in these cases we defer to the `HaploCart` classification which does not suffer from this bias by virtue of its pangenomic reference. For one sample (STR266b) the two tools disagreed between J1c and J1c5 at full coverage; we deferred in this case to the `HaploGrep2` prediction of J1c5.

The detailed results of the experiment can be seen in the Supplementary Section G.1. We observe that the number of aligned and processed reads has increased, as expected. More importantly, the improvement conferred by the RYmer correction strategy is also borne out in the sizeable decrease in mean Levenshtein distance between prediction and ground truth (Figure 7). These results strongly suggest that the RYmer index scheme works as intended, facilitating more sensitive alignment and a concomitant gain in accuracy on low-coverage ancient samples by allowing reads with deaminated seeds to be rescued and aligned.

**Figure 6:**
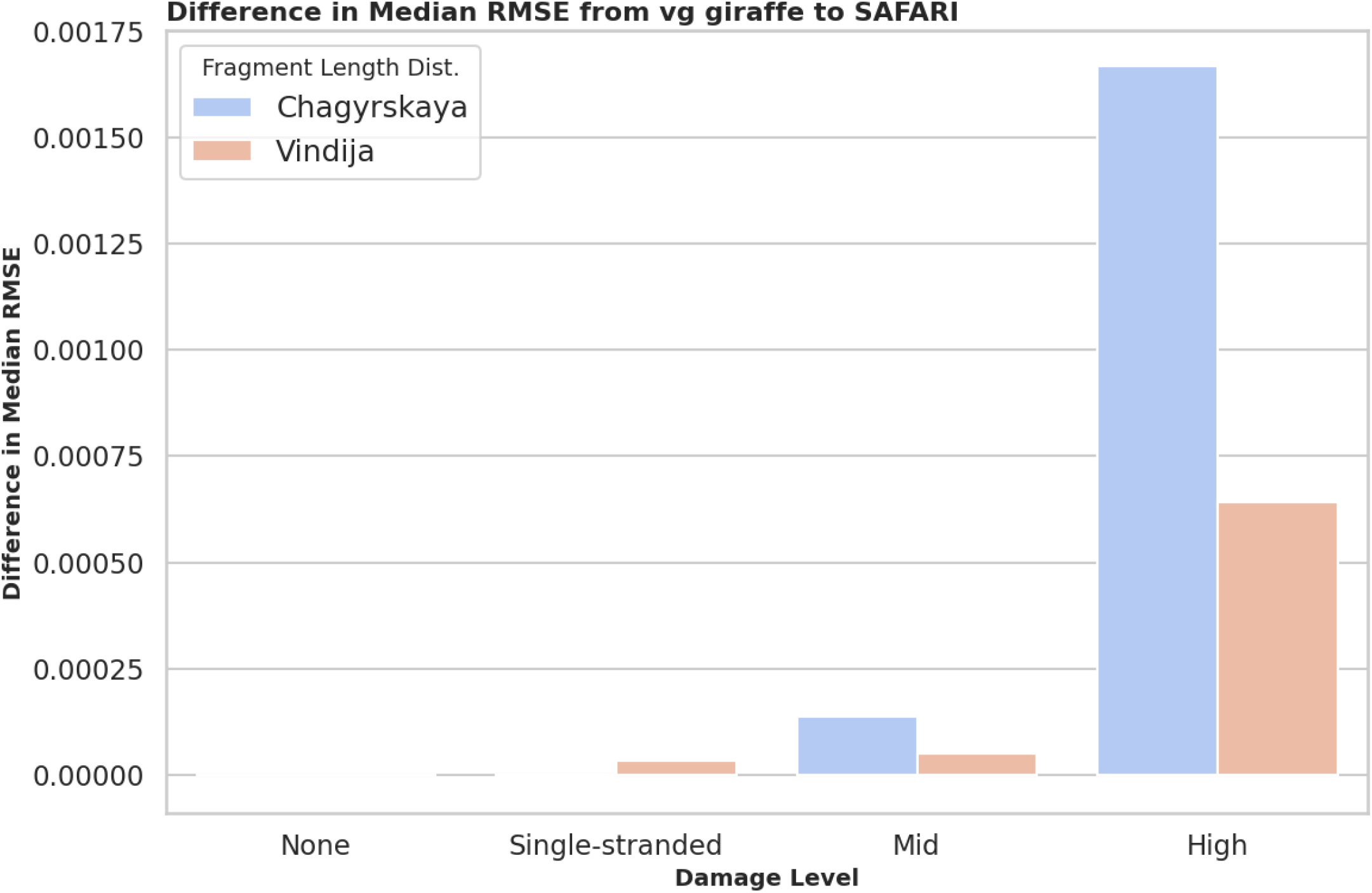
Difference in median root mean-squared error (RMSE) between ground truth and predicted nucleotide misincorporation matrices between `vg giraffe` and `SAFARI` at four distinct damage patterns, stratified by empirically derived fragment length distribution. Lower RMSE implies better performance, so if `vg giraffe` outperformed `SAFARI` the RMSE difference would be negative. We clearly see the improvement in inferred damage rate even for medium levels of damage but the improvement is particularly stark when high levels of damage are present in the input samples, due to the increased amount of information recovered from reads with deaminated seeds. Note that at zero damage the estimated damage matrices are equivalent.

**Figure 7:**
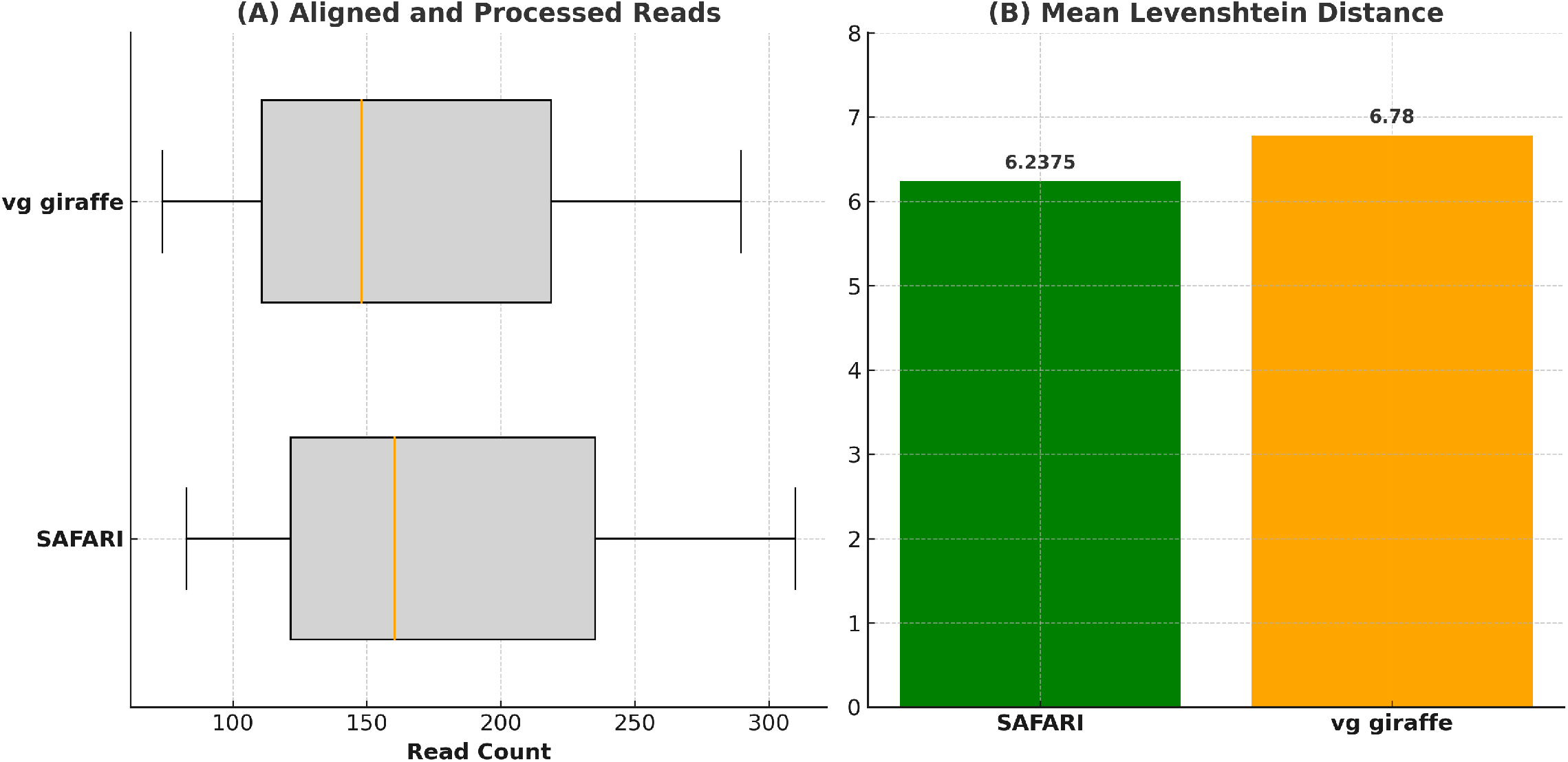
Comparison of performance metrics for `HaploCart` runs on ancient DNA samples before and after RYmer correction to the mapping. The median number of aligned and processed reads with `SAFARI` is 160.5 as compared to 148.0 for `vg giraffe`. The mean Levenshtein distance (edit distance) from the predicted to the ground truth is 6.2375 and 6.78 for `SAFARI` and `vg giraffe`, respectively. We report mean rather than median for the latter metric because the value is often zero.

**Figure 8:**
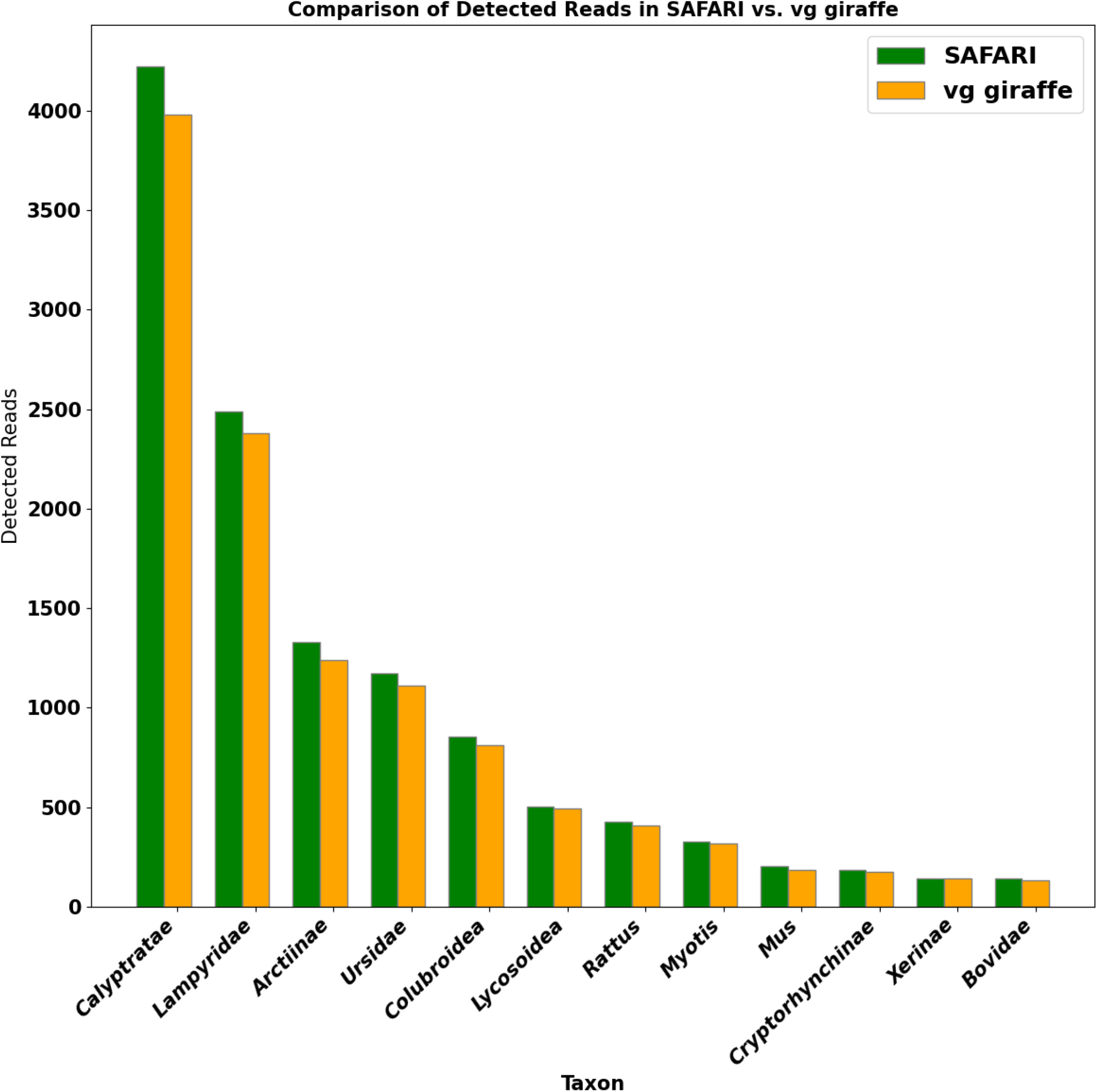
Detected number of fragments per taxon for `vg giraffe` vs. `SAFARI` in the `euka` experiment. In this case `SAFARI` is run with the default posterior threshold of 0.5. No false positive taxa were detected in either case. In almost all instances the number of fragments detected from the taxa increased after the RYmer correction was applied. The sole exception is *Xerinae* for which there was one less fragment (142 for `SAFARI`, 143 for `vg giraffe`).

### 3.3 euka with SAFARI

The second `vgan` subcommand that we modified to incorporate alignment with `SAFARI` was `euka`, a tool which characterizes the bilaterian composition of an ancient environmental DNA sample, using alignment with `vg giraffe` to a curated mtDNA database in conjunction with maximum likelihood methods of inference [56]. Since `euka` is explicitly designed for ancient data, it should benefit from `SAFARI`’s ancientdamage-aware modifications. Notably, for quality control purposes `euka` sets a lower bound on the number of alignments detected from a given taxon, so this aDNA correction to the alignment approach may well lead to detections which would otherwise go unnoticed.

We compared the two versions of the pangenome aligner on a simulated metagenomic cave environment from the Late Pleistocene [36]. In both cases, a minimum fragment count per detected taxon of 100, a minimum entropy score of 0.5, a minimum mapping quality score per read of 10 and a minimum number of six coverage bins per taxon was imposed. The Bayesian filter threshold for `SAFARI` alignment was the default of 0.5. A detailed description of the original environment can be found in [56]. Further details about our simulations are provided in Supplementary Section F.3. Our indexing parameters were *k* = 30, *w* = 20. A large kmer size was set to reduce false positives given the nature of metagenomic environmental samples. Default `SAFARI` parameters were employed, with the exception of the user-specified damage rate matrices which were passed in as the high-level damage profiles detailed in Supplementary Section J.1. The results of this experiment are seen in the Supplementary Section G.2. We found that `euka` in conjunction with `SAFARI` detected more fragments per taxon in almost all cases as compared with the original `vg giraffe`. The sole exception is *Xerinae* for which there was one less fragment (142 for `SAFARI`, 143 for `vg giraffe`). No false positive taxa were detected in either case. Although in this particular case the same number of taxa were detected, the detection threshold of 100 fragments was arbitrarily selected and it could easily be the case that the differential in detected read counts is enough for a taxon to be detected with `SAFARI` but not `vg giraffe` which may be very consequential for the study in question. Therefore this experiment provides another example of our aDNA correction strategy being able to improve inference on ancient samples.

In addition to lending further support to `SAFARI`’s ability to improve downstream analyses on aDNA samples, this experiment demonstrates that the very minor reduction in specificity introduced by the RYmer seed rescue procedure does not preclude its use on environmental or metagenomic samples despite the preponderance of exogenous DNA that have the potential to be spuriously mapped, so long as the user selects the indexing parameters and `SAFARI` posterior threshold sensibly.

Importantly, we note that the experiment also validates the utility of the user-defined posterior threshold in `SAFARI`. Supplementary Figure 5 demonstrates that the number of detected reads per taxon is indeed considerably sensitive to this threshold, with a more stringent (i.e. higher) threshold leading to a drop in the number of detected fragments. At a posterior threshold of 1.0, `SAFARI` and `vg giraffe` have the same behavior and lead to identical alignments, as expected, since in this case no RYmer seeds will pass the Bayesian filter.

### 3.4 Runtime and Memory Requirements

We performed a quick and direct benchmark of runtime and memory requirements using default indexing parameters (*k, w*) = (29, 11) for `SAFARI` as compared to those from the original `vg giraffe`. We did this for three different reference graphs. The first is a reference graph comprising just a single mitochondrial haplotype, namely that of the rCRS. The second is the newly curated panmitogenome for humans and archaic hominins which we describe in the Conclusion. And the third is a single-genome graph of chloroplast DNA from *Arabidopsis thaliana* (NCBI accession NC 000932.1, 154478 bases). In each case Illumina HiSeq 2500 reads (100 bp) at 100X coverage depth were simulated with `ART` [20], and the input was directly mapped by each of the two tools. The results are found in Table 2. For both the rCRS graph and the hominin panmitogenome the input data was generated from the rCRS mitogenome sequence. We see that for all reference graphs the time and memory requirements are considerably larger, and future work is needed to lessen the additional time and memory load, but the requirements should still be reasonable for most use cases. More runtime and memory benchmarks for the experiments conducted herein are provided in Supplementary Section I.

**Table 2:** Runtime and memory metrics comparing vg giraffe and SAFARI under deafult indexing parameters (*k* = 29, *w* = 11) on three different reference graphs of varying sizes with the addition of number of fragments.

| Ref. Graph | Tool | User Time (s) | Sys. Time (s) | Elapsed Time | Max Mem. (KB) | # Fragments |
| --- | --- | --- | --- | --- | --- | --- |
| rCRS | **vg giraffe** | 23.60 | 0.62 | 0:00:77 | 175,380 | 163,988 |
| rCRS | **SAFARI** | 501.46 | 4.18 | 0:10:03 | 556,036 | 163,988 |
| Hominin | **vg giraffe** | 757.56 | 4.79 | 0:15:20 | 907,244 | 163,988 |
| Hominin | **SAFARI** | 830.17 | 522.77 | 0:41:84 | 1,164,780 | 163,988 |
| Chloroplast | **vg giraffe** | 233.48 | 4.21 | 0:05:97 | 487,460 | 1,544,000 |
| Chloroplast | **SAFARI** | 1711.58 | 12.86 | 0:31:15 | 882,992 | 1,544,000 |

## 4 Conclusion

The seed-and-extend paradigm for full-genome nucleotide alignment has been the standard approach to short-read alignment for over thirty years, and has proven very effective for highly accurate reads [29, 2, 35]. However, in the special case of ancient DNA, the requirement for an exact seed match to a minimizer in the index before the extension phase can begin to adversely affect the sensitivity of the aligner due to the potential for deamination events in the seeds. This issue is especially problematic due to the inherently low-coverage and fragmentary nature of aDNA samples, in which even a small loss of data could be very consequential for any downstream inferences drawn from the alignments. By modifying the well-known haplotype-consistent pangenome aligner `vg giraffe`, we have demonstrated that the use of a RYmer index works well to rescue deaminated seeds and thereby improve the alignment of ancient samples. We have demonstrated that although the false positive rate increases somewhat under this indexing scheme, the increase is more than tolerable, in part due to our Bayesian filter which discards low-confidence seeds, and in part due to spurious seed matches failing to extend downstream in the `giraffe` pipeline. The efficacy of our RYmer indexing scheme is demonstrated by the improved performance of extant pangenome tools for ancient DNA (`HaploCart` and `euka`), the improved alignment statistics compared to standard `vg giraffe` in our surjected linear aligner experiment, and also by the improved damage rate estimation on the resulting BAM files. The latter result further shows that researchers who do not use pangenomes in their pipelines can still nonetheless benefit from our work.

We believe that `SAFARI` has the potential to improve our understanding of ancient DNA samples in a wide variety of settings and use cases. In this manuscript, we showcase its utility in pinpointing the fine-grained placement of a human sample’s mitochondrial lineage, and expanding our knowledge of eukaryotic mitodiversity in long-gone macroecological environments. We have even shown how surjection can be used to take full advantage of reference bias mitigation from pangenomes while still being able to work with BAM files to interface with tried and true NGS pipelines. But as alignment is in some sense agnostic to the nature of the DNA data (beyond its ancient damage profile) the applications are far wider than have been presented herein. The RYmer correction is, for example, not specific to mitochondrial DNA or even eukaryotic DNA. Moreover, the modified encoding scheme in its own right allows for the encoding of kmers of twice the length as before using the same hash table, an attractive feature which may have applications entirely beyond the field of ancient DNA.

Future work is essential to enhance the RYmer index concept, particularly its alignment capabilities with both pangenomes and linear genomes. For example it still remains to develop a RYmer implementation for syncmers, an alternate minimizer selection system currently available in `vg giraffe` [11]. Similarly, the introduction of minmers, which are not yet supported by `vg giraffe`, would be a valuable addition [22]. Moreover, our evaluations have been confined to mitochondrial genomes and one chloroplast genome, both of which are considerably smaller than, for instance, entire autosomes. As such, it is crucial to test the RYmer scheme in a nuclear context, where runtime and memory issues may be exacerbated, the adverse effects of false positive alignments may be compounded, and where multiple disconnected subgraphs corresponding to different chromosomes will be encountered. Optimal parameter settings for alignment using RYmer index-ing (e.g. stopping criteria for extensions, cap on the number of hits per RYmer, etc.) will need to be identified.

Finally, as alluded to earlier, we mention briefly that we are releasing as part of this paper a curated panmitogenome for humans and archaic hominins (including ancestrally reconstructed states) which we hope may be of use to researchers with relevant samples. More details, as well as a demonstration of improved alignment recovery when used in conjunction with `SAFARI`, are provided in Supplementary Section D.

## 5 Availability of Data and Materials

All scripts used in the benchmarking experiments are available at https://github.com/JoshuaDanielRubin/SAFARI_paper_data. The raw data underlying the results in the manuscript is available upon reasonable request to the corresponding author. The standalone `SAFARI` codebase can be found at https://github.com/grenaud/SAFARI. The FigShare DOI is 10.6084/m9.figshare.26763313. A static binary is provided for ease of use across operating systems.

## 6 Software Versions

`SAFARI` was modified from `vg` version 1.44.0 (Solara). Our `vgan` version was 2.0.2 (Miritillo). (We made minor modifications to this `vgan` version for the sole purpose of passing the additional `SAFARI` parameters into the internal `vg giraffe` subcommand call). Our `leeHom` version was 1.2.15. Our `Bowtie2` version was 2.4.4. Our `BWA` version was 0.7.17. Our `SHRiMP` version was 2.2.2. Our `BBMap` version was 38.91. Our `SAMtools` version was 1.13. Our `Snakemake` version was 5.10.0. Our `Python` version was 3.8.8. Our `seqtk` version was 1.3-r106.

## 7 Author Contributions

All authors conceived the idea for this paper. JDR implemented `SAFARI` and performed all benchmarking experiments, making use of previous scripts written by GR. JS was the primary contributor to the development of the `vg giraffe` subcommand and the GBWT graph data structure, and looked over the `SAFARI` manuscript and code. LK provided data and guidance on the spurious alignment model, as well as input on the manuscript. PWS provided infrastructure support for the project.

## 9 Acknowledgements

We express our gratitude to the `vg` toolkit development team for all of their efforts in creating the software backbone for this project. We thank Mikkel Schwartz for his assistance in constructing the full panmitogenome released in conjunction with this manuscript. We would also like to thank Nicola Vogel for her input on the manuscript. Finally, we thank Nanna Elmstedt Bild for her help in creating the graphical abstract and Antonio Fernandez-Guerra who coined the term RYmer. We also acknowledge that the large language model `GPT-4` [33] was used to assist with scripting and other tasks in connection with the preparation of this manuscript.

