## Supplementary Material for "SAFARI: Pangenome Alignment of Ancient DNA Using Purine/Pyrimidine Encodings"

October 8, 2024

**1** Department of Health Technology, Section for Bioinformatics, Technical University of Denmark, Kongens Lyngby, Denmark
**2** UC Santa Cruz Genomics Institute, Santa Cruz, USA

#### Contents

**A Variation Graph Overview** **3**

**B Fragment Length Distribution Visualizations** **3**

**C Damage Rate Estimate Visualizations** **4**

**D Human and Archaic Hominin Panmitogenome Release** **5**

### A Variation Graph Overview

For a good high-level overview of the variation graph data structure in general please see the Supplementary Material from [33], Section 1.

### B Fragment Length Distribution Visualizations

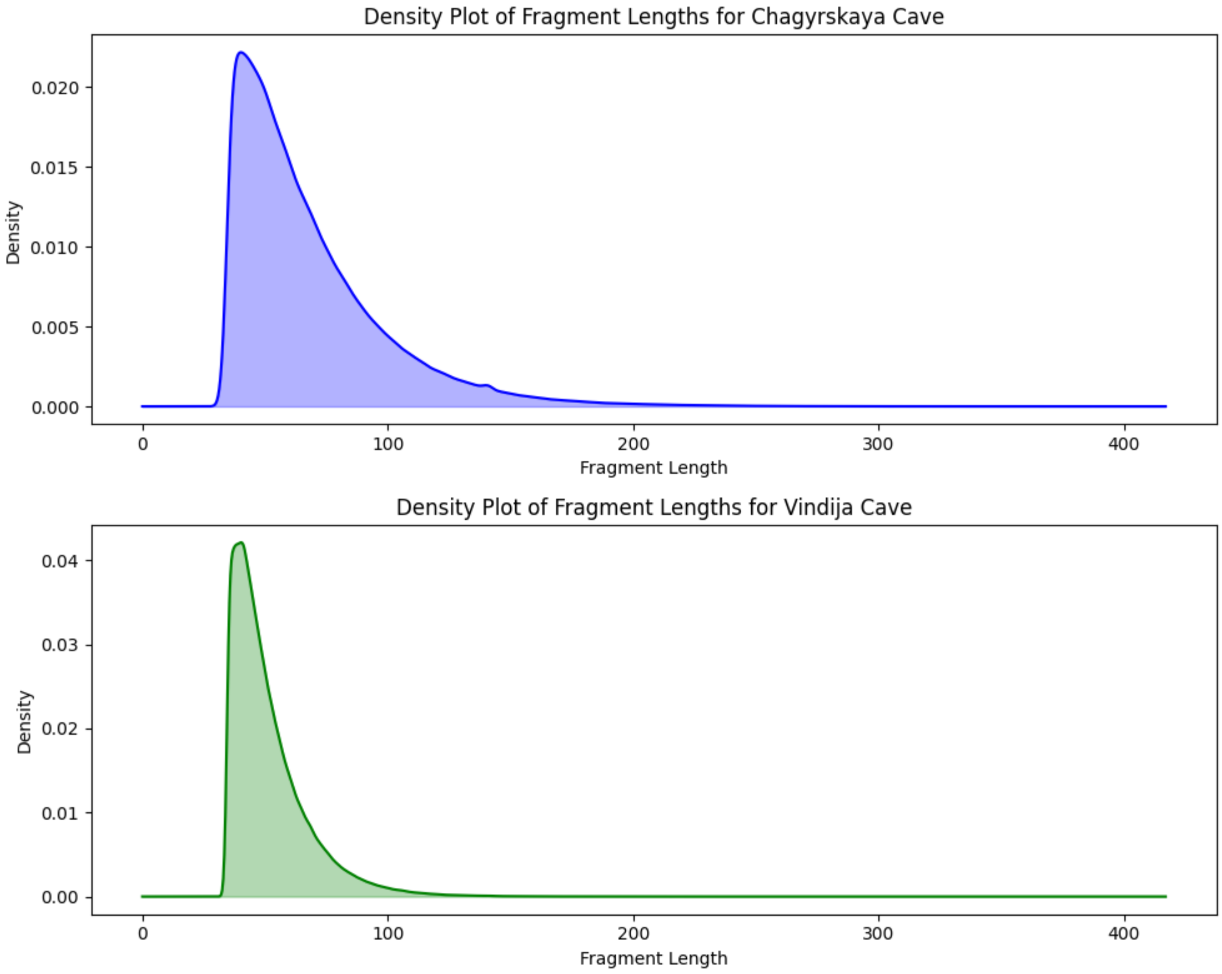

Figure 1: Fragment length distribution for the Chagyrskaya Cave [**TOP**] and Vindija Cave [**BOTTOM**] datasets, respectively.

### C Damage Rate Estimate Visualizations

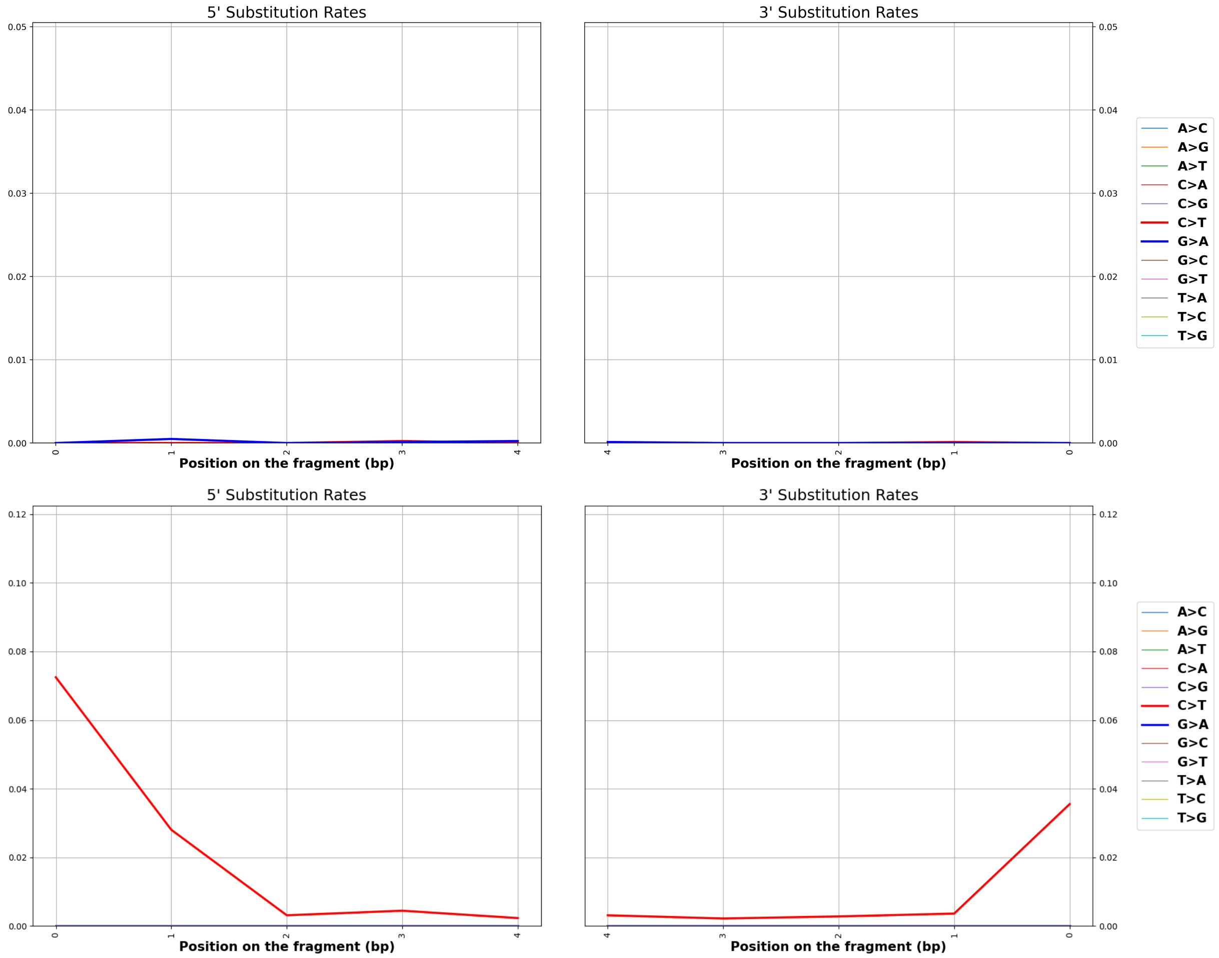

Figure 2: Examples of estimated damage rates for zero damage and single-stranded damage patterns used in our linear alignment experiments. Note that these BAM files were aligned with `SAFARI` but the corresponding BAM files aligned with `vg giraffe` are so similar that it would be difficult to distinguish.

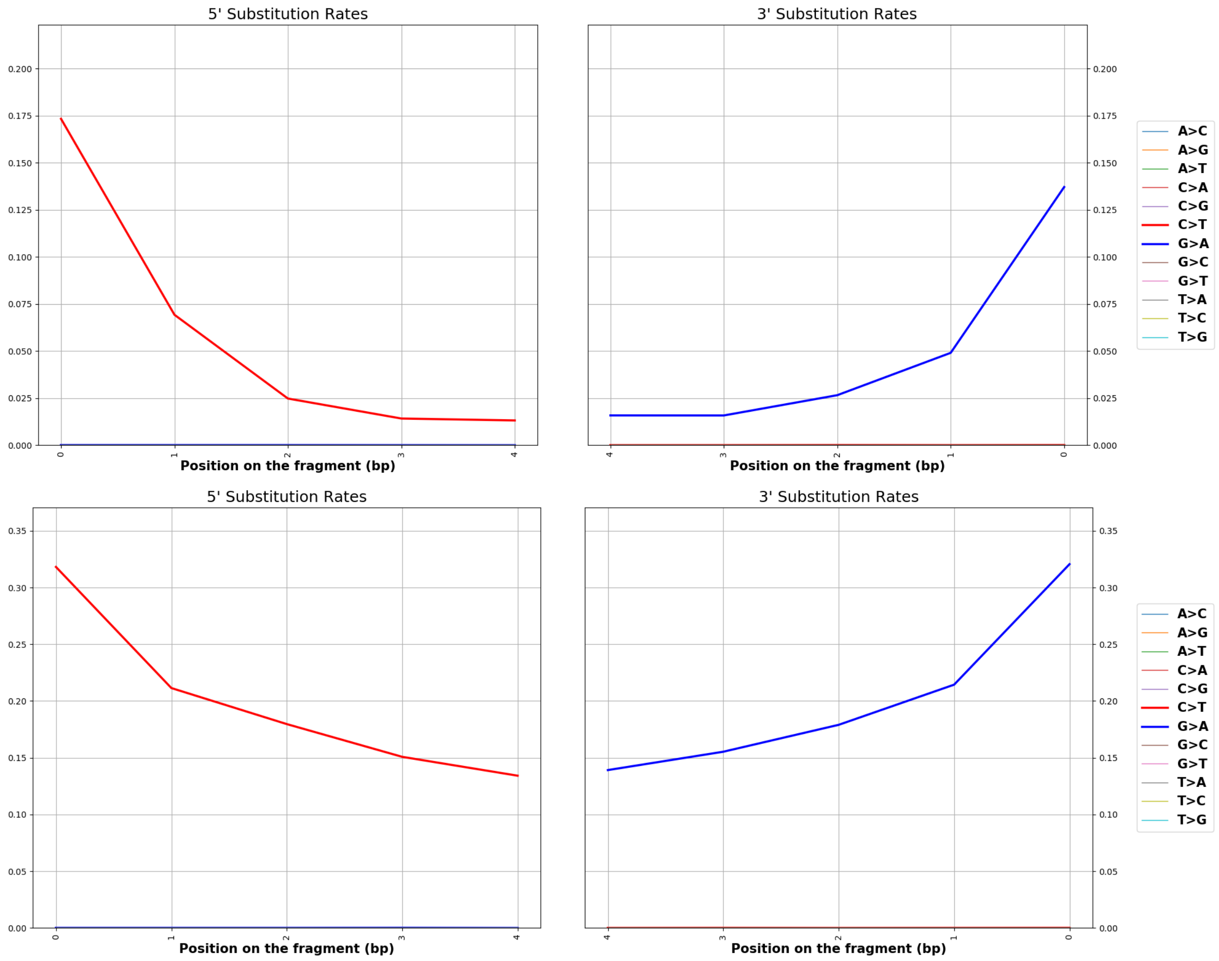

Figure 2: [**CONTINUED**] Examples of estimated damage rates for medium-level and high-level double-stranded damage patterns used in our linear alignment experiments.

### D Human and Archaic Hominin Panmitogenome Release

As part of this paper we are releasing a curated pangenome reference graph for human and archaic hominin mitochondria. We hope this will help some researchers studying relevant samples to make use of our aDNA-aware aligner. The graph contains an update to our `HaploCart` graph with 6,401 human haplogroup sequences (an 18% increase from Phylotree Build 17, based on [6]). The increased resolution will better represent extant human mtDNA haplotype diversity. In addition there are archaic hominin sequences as well as ancestrally reconstructed states in order to better capture the true variation in hominin mitodiversity even for variation which no longer exists. In all the graph comprises 12,852 human and archaic hominin mitogenomes, to the best of our knowledge the largest panmitogenome ever published. These files are hosted in our in-house FTP server. In brief, the files can be retrieved via the Linux command

```
wget -nc -l1 --recursive --no-parent -P .
ftp://ftp.healthtech.dtu.dk:/public/SAFARI_graph/
```

Further details about the graph are provided in the subsections below.

#### D.1 Table of Archaic Hominin Sequences in the Human and Archaic Hominin Panmitogenome Release

Below we list all archaic hominin sequences used in the newly released panmitogenome as part of this paper.

Table 1: Comprehensive list of archaic hominin genomes used in the newly released panmitogenome.

| Accession | Description |
|---|---|
| MT576650.1 | Homo sapiens neanderthalensis isolate Denisova17 mitochondrion, complete genome |
| MT921957.1 | Homo sapiens neanderthalensis mitochondrion, complete genome |
| KU131206.2 | Homo sapiens neanderthalensis isolate DC1227 mitochondrion, complete genome |
| MG025536.1 | Homo sapiens neanderthalensis isolate Les Cottes Z4-1514 mitochondrion, complete genome |
| KX198085.1 | Homo sapiens neanderthalensis isolate GoyetQ374a-1 mitochondrion, complete genome |
| KX198086.1 | Homo sapiens neanderthalensis isolate GoyetQ305-7 mitochondrion, complete genome |
| MG025540.1 | Homo sapiens neanderthalensis isolate Goyet Q56-1 mitochondrion, complete genome |
| MG025538.1 | Homo sapiens neanderthalensis isolate Spy 94a mitochondrion, complete genome |
| MG025539.1 | Homo sapiens neanderthalensis isolate Vindija 87 mitochondrion, complete genome |
| NC_011137.1 | Homo sapiens neanderthalensis mitochondrion, complete genome |
| KX198088.1 | Homo sapiens neanderthalensis isolate GoyetQ57-2 mitochondrion, complete genome |
| MT677921.1 | Homo sapiens neanderthalensis mitochondrion, complete genome |
| MG025537.1 | Homo sapiens neanderthalensis isolate Mezmaiskaya 2 mitochondrion, complete genome |
| KX198087.1 | Homo sapiens neanderthalensis isolate GoyetQ305-4 mitochondrion, complete genome |
| MK388903.1 | Homo sapiens neanderthalensis isolate Chagyrskaya8 mitochondrion, complete genome |
| MT795654.1 | Homo sapiens neanderthalensis isolate Stajnia S5000 mitochondrion, complete genome |
| OM062614.1 | Homo sapiens neanderthalensis isolate Mezmaiskaya 3 mitochondrion, complete genome |
| MK033602.1 | Homo sapiens neanderthalensis isolate Denisova15 mitochondrion, complete genome |
| MK123269.1 | Homo sapiens neanderthalensis isolate Scladina I-4A mitochondrion, complete genome |
| KY751400.2 | Homo sapiens neanderthalensis mitochondrion, complete genome |
| NC_023100.1 | Homo heidelbergensis mitochondrion, complete genome |
| KT780370.1 | Homo sapiens subsp. 'Denisova' isolate Denisova8 mitochondrion, complete genome |
| KX663333.1 | Homo sapiens subsp. 'Denisova' isolate Denisova2 mitochondrion, complete genome |

Continued on next page

| Accession | Description |
|---|---|
| MT576651.1 | Homo sapiens subsp. 'Denisova' isolate Denisova19 mitochondrion, complete genome |
| MT576653.1 | Homo sapiens subsp. 'Denisova' isolate Denisova21 mitochondrion, complete genome |
| MT576652.1 | Homo sapiens subsp. 'Denisova' isolate Denisova20 mitochondrion, complete genome |

#### D.2 Phylogeny Inference and Ancestral State Reconstruction

We used `IQ-TREE` [23] to infer the phylogeny including ancestral states from our database of 6,401 modern human mitogenomes plus archaic hominin sequences (see above subsection). The choice of tool was based on feasibility given the inordinately large tree size; `IQ-TREE` is especially good at optimizing memory consumption, and for this reason previous researchers building large human mitochondrial phylogenies have also made use of this tool [2]. The program infers ancestral states in conjunction with inference of the phylogeny using a maximum likelihood approach. We construct the tree under a HKY DNA substitution model which allows for variable base frequencies (all bases) and a variable transition/transversion ratio, both essential features when dealing with mtDNA [12]. Specifically, we used the command

```
nice -19 iqtree -s [INPUT FILE] -m HKY -nt AUTO -redo -pre tree_tmp/haps -asr
```

#### D.3 Panmitogenome Construction

The hominin panmitogenome was constructed with `PGGB` [7]. The exact command used was as follows:

```
nice -19 parallel -j 55 --bar "nice pggb -i [INPUT MULTIFASTA] -t 50 -o final_s -p 70
-s  -H 12810 -n 5000 -k 300 -G 3079,3559 -l 20" :::  1100
```

We then used the `vg mod` subcommand to impose a maximum sequence size on each node of six nucleobases, as we found this empirically to help improve results. The final graph has 15722 nodes, 21551 edges, and 12852 embedded haplotype paths, and forms a single circularized connected component to reflect the biology of the eukaryotic mitogenome.

#### D.4 Improved Mapping of aDNA Reads from Archaic Hominins

Figure 3 shows how using `SAFARI` in conjunction with the released human and archaic hominin panmitogenome translates into more alignments for archaic samples in the presence of damage. The Denisovan sample used here has NCBI accession KT780370.1. The high-damage profiles are taken from [8]. With the exception of the user-inputted damage profiles for `SAFARI`, for which the appropriate ATP2 profile matrices were passed in, default parameters were used throughout for both tools. The number of aligned reads for each `in silico` sample was determined via the command `vg stats -a`. The data is presented in tabular form in Table 2 below.

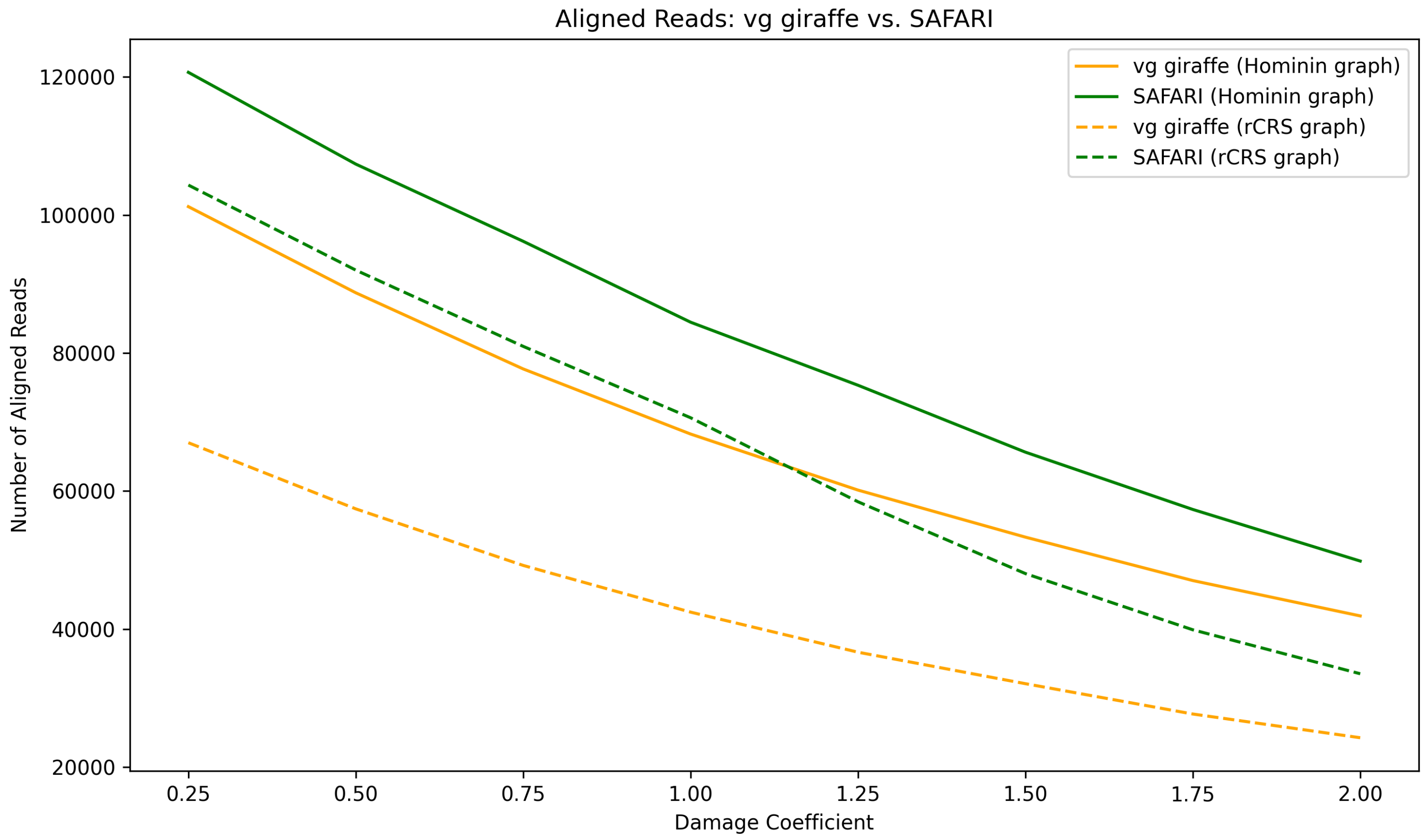

Figure 3: Number of aligned reads mapping with `vg giraffe` vs. `SAFARI` at various deamination rates for a Denisovan sample (generated *in silico*). The damage coefficient is a constant factor by which the deamination rates vary (in other words if the damage coefficient doubles then deamination events are twice as frequent). The dashed lines show data mapped to the linear reference graph containing only the rCRS reference genome, while the solid lines show data mapped to the full panmitogenome.

Table 2: Number of aligned reads mapping with `vg giraffe` vs. `SAFARI` at various deamination rates for a Denisovan sample (generated *in silico*).

| Deamination factor | `vg giraffe` Hominin | `SAFARI` Hominin | `vg giraffe` rCRS | `SAFARI` rCRS |
|---|---:|---:|---:|---:|
| 0.25 | 101,200 | 120,660 | 66,992 | 104,326 |
| 0.50 | 88,719 | 107,364 | 57,406 | 92,001 |
| 0.75 | 77,684 | 96,179 | 49,214 | 80,955 |
| 1.00 | 68,250 | 84,455 | 42,445 | 70,610 |
| 1.25 | 60,127 | 75,319 | 36,638 | 58,435 |
| 1.50 | 53,311 | 65,623 | 32,076 | 48,029 |
| 1.75 | 47,019 | 57,320 | 27,681 | 39,885 |
| 2.00 | 41,896 | 49,846 | 24,244 | 33,515 |

##### D.5 Effect of Misspecification of Damage Rate Matrices into `SAFARI`

As part of its model `SAFARI` requires the user to input estimated nucleotide misincorporation rates for the sample. It is therefore interesting to ask how sensitive the performance of the program is to these estimates, and what the effect would be of underestimation or overestimation of damage in the input sample.

We generated 500,000 endogenous reads from a Denisovan sample (NCBI accession KT780370.1) [31]. These reads were contaminated (*in silico*) with 100,000 bacterial reads from the `RefSoil` database [5]. We then aligned the data with `SAFARI` at varying input estimated damage rate matrices. We did this by taking the ATP2_mt_1.00 rate matrices and multiplying the rates by a constant multiplicative factor which we term the damage coefficient. We vary the damage coefficient from 0.25 to 2.00, inclusive, at a step size of 0.25. This procedure was performed twice, once aligning to the full hominin panmitogenome and once to the single-haplotype graph comprising solely the rCRS haplotype H2a2a1.

We can see from Table 3 that, first and foremost, the program is not entirely insensitive to the damage rate estimates, in that varying the estimates will have an effect on the resulting alignments. However in general the program is quite robust to misspecifications of the damage rates. Furthermore, it is interesting to see that underestimation of damage seems to adversely affect performance more severely than overestimation of damage for both reference graphs, but this difference in sensitivity to error in the estimation is minor. Interestingly in both cases the number of resulting alignments as well as $F_1$ score peaks at a damage coefficient of 1.25, indicating that the `SAFARI` model is fairly well-calibrated, but could still benefit from minor adjustments in future work, as we would expect the performance to be optimal when the estimated rate matrices are closest to the true rate matrices.

Table 3: Alignment statistics and performance metrics for `SAFARI` mapping bacterially-contaminated Denisovan reads to both panmitogenomes at varying damage coefficient values (which affects the user-defined damage rate estimates passed into the program). All decimals are rounded to six figures.

| Panmitogenome | Damage Coeff | TP | FP | TN | FN | Sensitivity | Specificity | $F_1$ |
|---|---|---|---|---|---|---|---|---|
| Hominin | 0.25 | 82890 | 245 | 24988 | 12 | 0.999855 | 0.990290 | 0.998452 |
| Hominin | 0.5 | 83023 | 248 | 24988 | 12 | 0.999855 | 0.990173 | 0.998437 |
| Hominin | 0.75 | 83191 | 248 | 24988 | 12 | 0.999856 | 0.990173 | 0.998440 |
| Hominin | 1.0 | 83333 | 248 | 24988 | 12 | 0.999856 | 0.990173 | 0.998442 |
| Hominin | 1.25 | 83482 | 248 | 24988 | 12 | 0.999856 | 0.990173 | 0.998445 |
| Hominin | 1.5 | 83377 | 248 | 24988 | 12 | 0.999856 | 0.990173 | 0.998443 |
| Hominin | 1.75 | 83406 | 248 | 24988 | 12 | 0.999856 | 0.990173 | 0.998444 |
| Hominin | 2.0 | 83457 | 249 | 24988 | 12 | 0.999856 | 0.990134 | 0.998439 |
| rCRS | 0.25 | 75892 | 210 | 24998 | 2 | 0.999974 | 0.991669 | 0.998605 |
| rCRS | 0.5 | 76224 | 210 | 24998 | 2 | 0.999974 | 0.991669 | 0.998611 |
| rCRS | 0.75 | 76457 | 210 | 24998 | 2 | 0.999974 | 0.991669 | 0.998616 |
| rCRS | 1.0 | 76707 | 210 | 24998 | 2 | 0.999974 | 0.991669 | 0.998620 |
| rCRS | 1.25 | 76961 | 210 | 24998 | 2 | 0.999974 | 0.991669 | 0.998625 |
| rCRS | 1.5 | 76878 | 210 | 24998 | 2 | 0.999974 | 0.991669 | 0.998623 |
| rCRS | 1.75 | 76921 | 210 | 24998 | 2 | 0.999974 | 0.991669 | 0.998624 |
| rCRS | 2.0 | 76921 | 210 | 24998 | 2 | 0.999974 | 0.991669 | 0.998624 |

#### D.6 Surjection After Mapping to a Pangenome Can Benefit from Variation Even If the Linear Reference Is Identical

Many ancient DNA researchers are accustomed to working with NGS pipelines which make use of a linear reference genome for alignment, and do processing on BAM files. Such researchers may well wonder if there is much point in mapping to a graph if they will need to surject alignments back to their reference haplotype downstream anyway, or whether all advantages accrued by the pangenome will be nullified after collapsing all the variation in the reference.

To test this we ran a quick experiment to show that indeed performance can improve by mapping to the full graph first and then surjecting afterwords. We took the same bacterially-contaminated Denisovan sample as in Section D.5 above. This sample was mapped by both aligners under identical parameters, with the ATP2_mt_1.00 profiles being input into `SAFARI`. The mapping was performed twice, once to the single-haplotype rCRS reference graph, and then again to the full hominin panmitogenome, followed by surjection to the rCRS reference (H2a2a1).

Table 4: Comparison of performance metrics between aligning to the full hominin panmitogenome first and then surjecting onto the rCRS haplotype **[LEFT]** and directly aligning to the rCRS-only pangenome **[RIGHT]**. The former strategy can improve sensitivity, which is often critical for aDNA studies.

| **Metric** | **Full Panmitogenome** | **rCRS Graph** |
|---|---|---|
| Corrected Alignments | 87398 | 76707 |
| Unmapped Bacteria | 24988 | 24998 |
| Mapped Bacteria | 12 | 2 |
| Mismapped Alignments | 258 | 210 |

We see in Table 4 that considerably more reads can be aligned by first aligning to the full graph and then surjecting to the linear reference, despite the fact that the linear reference haplotype is very divergent from the true underlying haplotype in the sample. This is because if the panmitogenome contains reference genomes less divergent from the sample than the linear reference, it will store kmers in the index from those more similar genomes and extend and produce graphical alignments based on information from those more similar genomes. Then when surjection happens there will be mismatches etc. due to dissimilarity between the sample and the linear reference being surjected onto, but the alignments will still be produced, unlike if mapping directly where more alignments may fail either at the seed matching stage or the extension stage.

Although this procedure of pangenome mapping followed by surjection does lead to an increase in false positive mappings this issue is largely mitigated by the Bayesian filter in `SAFARI`.

### E Inference Model Details

#### E.1 User-defined Model Parameters in SAFARI

We briefly discuss here some advice and general comments about user-defined parameters in our model. There are three user-defined parameters for SAFARI, namely the estimated nucleotide misincorporation matrices, the prior on a correct alignment of a base, and the posterior threshold. A discussion about the former (the estimated rate matrices) can be found in Section D.5.

The prior probability that a user should have on the likelihood that an arbitrary seed from their ancient sample will correctly align to the reference, conditionalized on a match of that seed to some indexed RYmer in the RYmer index (the flag "-V" in SAFARI) is heavily dependent on the nature of the sample. The degree of preservation, depth and breadth of coverage, and the specific methods used for DNA extraction, possible treatment, and sequencing are all factors which would have a substantial impact on this value. The user is therefore in a better position to provide a more informative prior than we are. Accordingly, we allow her to provide this information as a command-line argument. Similarly, the user can also control the posterior threshold (the flag "-j") to be able to adjust the sensitivity/specificity tradeoff to her requirements.

Unfortunately, even if the user has a good understanding of the sample in question, the choice of these parameters may still be difficult to fine-tune. This is because the probability will be very much dependent on the pangenome reference being used, the indexing parameters, and the nature of the sample, and future work is required to better understand and develop a model for this. Nonetheless the user may be able to use something like a grid search to identify optimal parameters for their sample, or perhaps best practices will be decided over time, similar to how optimal parameter settings for standard linear alignment tools for aDNA samples have reached broad consensus in the community.

#### E.2 Determination of Likelihood Function for Spurious Alignment

The question of what likelihood function to assign to model 2 (i.e. a spurious seed hit to the RYmer index) is a difficult one because in theory it will depend heavily on factors such as contamination rate and composition, which cannot be known *a priori*. In order to obtain a somewhat reasonable distribution, therefore, we generated a basic *in silico* simulation in Python to understand at a very coarse-grained level how the number of potentially deaminated mismatches between the seed and the candidate RYmer hit varies with the choice of kmer size $k$. Here we describe how we determined our likelihood function for a spurious alignment, conditionalized on the fact that there is a match in RYmer space between a seed and a hit.

We used the standard human mitochondrial reference, called the revised Cambridge Reference Sequence (rCRS), as our reference genome [1]. We indexed the minimizers and RYmers in the reference (https://github.com/JoshuaDanielRubin/SAFARI_paper_data/blob/main/spurious_model/main.py). The bacterial contamination FASTA file (https://github.com/JoshuaDanielRubin/SAFARI_data/blob/main/spurious_model/refSoilSmall.fa) comprising twenty bacterial genomes was obtained via accession numbers from the RefSoil database [5]. The mean genome length for the original file of 766 reference sequences was 4,396,003.8 bp. For reasons of space we used only the first twenty reference sequences to infer our spurious alignment model parameters.

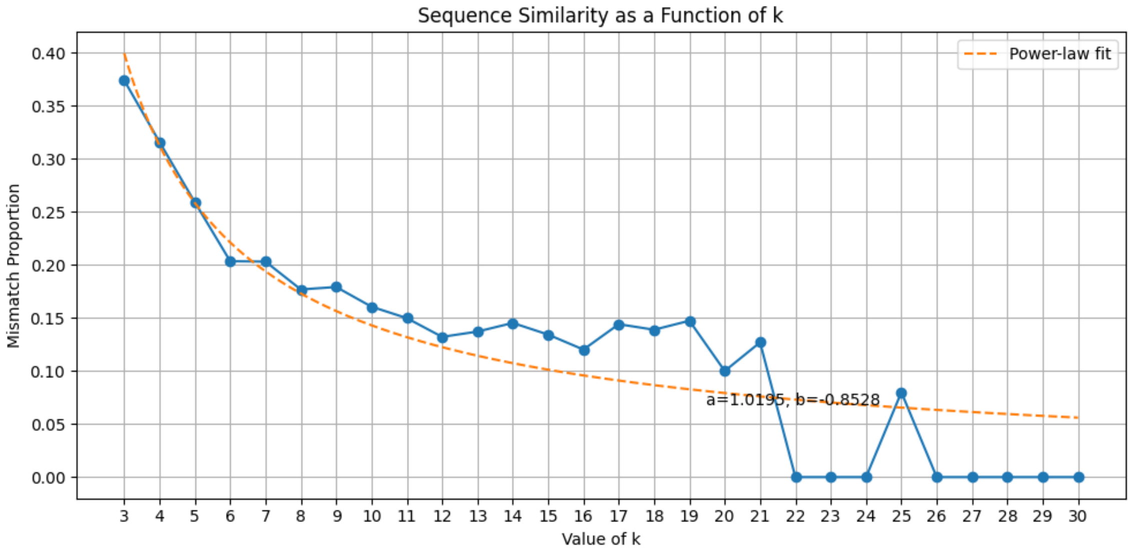

Figure 4: *in silico* `Python` simulations of aDNA RYmers and minimizers at varying values of $k$. The distribution roughly obeys a power law behavior which plateaus at high $k$ values.

We emphasize that this likelihood function is very coarse-grained due to aleatoric uncertainties about the input sample which cannot in principle be known to us at runtime. However, we can confirm that this model is considerably more accurate in practice than a model assuming random DNA of equal base frequency for both the reference and the input data (where each base has probability 0.5 of mismatching in kmer space).

### F Experiment Details

#### F.1 HaploCart Experiment

The script for downloading twenty ENA samples for the HaploCart experiment can be found at https://github.com/JoshuaDanielRubin/SAFARI_paper_data/blob/main/haplocart_ancient/download.sh. The script for downsampling samples can be found at https://github.com/JoshuaDanielRubin/SAFARI_paper_data/blob/main/haplocart_ancient/downsample.sh.

Note as well that for this experiment, we run with the -np flag, i.e. forgoing the phylogeny-aware confidence estimation as it is not relevant. We use the high-level damage matrices described in Section J.1 as input estimates.

#### F.2 Linear Aligner Benchmarking Experiment

Samples for the linear aligner benchmarking experiment were generated using the workflow manager Snakemake [13]. The Snakefile can be found at https://github.com/JoshuaDanielRubin/SAFARI_paper_data/blob/main/linear_experiment/human_mito/Snakefile.

In addition to our primary goal of comparing standard vg giraffe to SAFARI we also wanted to understand how well these tools perform with respect to commonly-used linear alignment tools. We therefore benchmarked against SHRiMP [30], BWA aln (both standard and ancient-specific parameters) [18, 21], BWA-MEM [16], BBMap [4], and Bowtie2 [14]. Though it is seldom used in the aDNA community and no longer maintained, SHRiMP uses a spaced kmer scheme which allows more tolerance of mismatches than standard kmer indexing systems, and we have internally found it to be highly effective for aligning aDNA samples [30].

Our reference genome was the rCRS (https://github.com/JoshuaDanielRubin/SAFARI_paper_data/blob/main/linear_experiment/human_mito/simulations/gen_0.fa). We stratified by four separate damage matrices. Three represent zero damage, medium-level damage, and high-level damage rates (i.e. nucleotide misincorporation patterns) for double-stranded DNA libraries. The fourth represents damage rates for single-stranded (otherwise know as Santa Cruz) libraries, which have been shown to be optimal in many cases for aDNA samples [10]. Further information about these matrices is provided in Section J.1.

We generated 300,000 circular (i.e. the end wraps around to the beginning, reflecting the circularity of mtDNA) mitochondrial fragments and 1,000 NuMT fragments using the fragSim subcommand of gargammel [32, 11, 28]. The fragment length distribution passed into fragSim via the -s flag was from either the empirically derived distribution from Vindija Cave, or that from Chagyrskaya Cave, depending on which dataset was being generated [20, 25]. aDNA damage was simulated using the deamSim subcommand of gargammel [28].

NuMTs are nuclear pseudogenes originating from the mitochondria which often mismap to the mitochondria [35]. They are found at low abundance relative to bona fide mtDNA, but in order to be extra charitable in our benchmarking and to ensure that our RYmer correction procedure is relatively robust to a high amount of false positives the proportion of NuMTs in our samples was well above the biological expectation. These fragments were contaminated *in silico* with 100,000 bacterial fragments from a *Campylobacter col* reference genome (NCBI accession NC_022660.1) [3] retrieved from the RefSoil database discussed in Section E.2. Fragment lengths for both the endogenous and contaminant material obeyed an empirically derived distribution from a high-coverage 80,000-year-old Neanderthal sample from Chagyrskaya Cave [20]. Samples were subsampled at eighteen subsampling rates (0.1 to 0.95 inclusive, step size 0.05) using seqtk [17]. Adapter sequences were added using the adptSim subcommand of gargammel. Illumina HiSeq 2500 short reads were then simulated using ART [9] at a fixed read length of 125 bp. Adapter sequences were trimmed and these reads were merged using leeHom with the --ancientdna flag enabled [27]. Exact command and parameter settings can be found in the Snakefile. A total of 568 BAM files were produced by each of the eight alignment tools under evaluation, yielding a total sample size of 4544 alignment files.

##### F.2.1 Exact Commands for the Linear Benchmarking Experiment

For `BWA aln` we follow [21] in setting the aDNA-specific parameters. For `BWA-MEM` we follow [34]. For `Bowtie2` we follow [14]. Parameters for the other tools were decided upon from prior experience/experimentation.

Table 5: Exact commands used in the linear aligner benchmarking experiment.

| Rule | Command |
|---|---|
| BWA-MEM | interleavefastq.sh {input_1} {input_2} {input_3} \| bwa mem -r 2.5 -k 19 -p simulations/gen_0.fa /dev/stdin \| samtools sort /dev/stdin > {output} |
| BWA aln | cat <(bwa aln simulations/gen_0.fa {input_1} \| bwa samse simulations/gen_0.fa /dev/stdin {input_1}) <(bwa sampe simulations/gen_0.fa <(bwa aln simulations/gen_0.fa {input_2}) <(bwa aln simulations/gen_0.fa {input_3}) {input_2} {input_3} \| samtools view -S /dev/stdin) \| samtools sort /dev/stdin > {output} |
| BWA aln (anc) | cat <(bwa aln -n 0.02 -o 2 -l 1024 simulations/gen_0.fa {input_1} \| bwa samse simulations/gen_0.fa /dev/stdin {input_1}) <(bwa sampe simulations/gen_0.fa <(bwa aln -n 0.02 -o 2 -l 1024 simulations/gen_0.fa {input_2}) <(bwa aln -n 0.02 -o 2 -l 1024 simulations/gen_0.fa {input_3}) {input_2} {input_3} \| samtools view -S /dev/stdin) \| samtools sort /dev/stdin > {output} |
| Bowtie2 | interleavefastq.sh {input_1} {input_2} {input_3} \| bowtie2 --end-to-end --sensitive -x bowtie2index/gen_0.fa --interleaved /dev/stdin \| samtools sort /dev/stdin > {output} |
| SHRiMP | cat <(gmapper -N 1 -o 1 --single-best-mapping --sam-unaligned --fastq --sam --no-qv-check --qv-offset 33 {input_1} simulations/gen_0.fa) <(gmapper -N 1 -o 1 --single-best-mapping --sam-unaligned --fastq --sam --no-qv-check --qv-offset 33 -1 {input_2} -2 {input_3} simulations/gen_0.fa \| samtools view -S /dev/stdin) \| samtools sort /dev/stdin > {output} |
| BBMap | cat <(bbmap.sh sam=1.3 qin=33 ref=simulations/gen_0.fa nodisk=t in={input.input_1} out=/dev/stdout) <(bbmap.sh sam=1.3 qin=33 ref=simulations/gen_0.fa nodisk=t in={input.input_2} in2={input.input_3} out=/dev/stdout \| samtools view -S /dev/stdin) \| samtools sort /dev/stdin > {output} |
| SAFARI | nice -19 vg safari -f /dev/stdin -m graph.min -d graph.dist -Z graph.giraffe.gbz --deam-3p [3p DEAM PROFILE] --deam-5p [5p DEAM PROFILE] -j 0.2 \| vg surject -x graph.xg -P -F PATHS -b /dev/stdin \| samtools sort /dev/stdin > {output} |
| vg giraffe | nice -19 interleavefastq.sh input_1} {input_2} {input_3} \| vg giraffe -f /dev/stdin -m graph.min -d graph.dist -Z graph.giraffe.gbz \| vg surject -x graph.xg -P -F PATHS -b /dev/stdin \| samtools sort /dev/stdin > {output} |

##### F.2.2 $(k, w)$ Indexing Parameter Pairs Evaluated

In addition to the default indexing parameters we also aligned all samples under 25 other distinct indexing parameter pairs for both `vg giraffe` and `SAFARI` in order to understand how varying these parameter values affect the behavior of these tools. Below (Table 6) is a table listing the parameter pairs that were evaluated.

Table 6: Indexing parameter pair values for `SAFARI` and `vg giraffe` evaluated in the linear alignment experiment.

| $k$ | $w$ |
|---|---|
| 10 | 10 |
| 10 | 15 |
| 10 | 2 |
| 10 | 5 |
| 10 | 7 |
| 10 | 8 |
| 11 | 6 |
| 11 | 7 |
| 12 | 3 |
| 12 | 4 |
| 12 | 5 |
| 12 | 6 |
| 12 | 7 |
| 15 | 10 |
| 15 | 15 |
| 15 | 5 |
| 18 | 8 |
| 20 | 10 |
| 20 | 15 |
| 20 | 2 |
| 20 | 5 |
| 29 | 11 |
| 30 | 10 |
| 30 | 15 |
| 30 | 2 |
| 30 | 5 |

##### F.2.3 Damage Rate Estimation Experiment

**Populating the MD Field in `SAFARI` and `vg giraffe` Alignments** It is a minor annoyance that natively `SAFARI` and `vg giraffe` surjected alignments in BAM file format do not populate the MD field required to represent mismatches between the read and the reference (although mismatches are directly reported in GAM file format). Therefore these fields were populated afterwards via the `samtools calmd` subcommand [19]. For the script that does this please see `https://github.com/JoshuaDanielRubin/SAFARI_paper_data/blob/main/linear_experiment/human_mito/make_MD.sh`.

**Damage Profile Estimates with `bam2prof`** `bam2prof` is a small script for estimating nucleotide misincorporation rates from BAM file input [26]. For the samples with damage patterns arising from a single-stranded library preparation we ran `bam2prof` with flags `-minl 20 -q -single -minq 0 -length 5`. For all other samples we run with the flags `-minl 20 -q -both -minq 0 -length 5`.

#### F.3 Simulated Metagenomic Cave Environment Details

For full details of the simulated metagenomic cave environment used in the `euka` experiment please see https://github.com/nicolaavogel/eukaPaperData/tree/master/CreateSimulatedEnv. There you will find the precise bacteria, fungi, archaea, and viruses included in the sample. A table of the present eukaryotic taxa is provided below. In total there were 1500000 reads in the full sample.

| |
|---|
| *Bovidae* |
| *Calyptratae* |
| *Colubroidea* |
| *Cryptorhynchinae* |
| *Lampyridae* |
| *Lycosoidea* |
| *Mus* |
| *Myotis* |
| *Neognathae* |
| *Rattus* |
| *Sciurinae* |
| *Ursidae* |
| *Xerinae* |

Table 7: Eukaryotic taxa present in the simulated metagenomic environment.

### G Experiment Figures and Tables

#### G.1 HaploCart Experiment

Table 8: Comparison of predicted human mtDNA haplogroup and number of processed reads between the original and modified HaploCart versions at four target mean coverage depths (0.25X, 0.5X, 1X, and 2X) over five replicates. Full coverage predictions for HaploCart and HaploGrep2, respectively, are provided in the third column. Across the board, the total number of processed reads increases. More importantly, the predictions are more accurate with respect to edit distance between predicted and ground truth haplogroup labels as described in the main article Section 3.2.

| Sample | Rate | Full_Cov_Pred | Replicate | HG_corr | HG_uncorr | Reads_corr | Reads_uncorr |
|---|---|---|---|---|---|---|---|
| AED204 | 0.25X | X2b11 | 1 | X1’2’3 | B4a2 | 49 | 42 |
| AED204 | 0.25X | X2b11 | 2 | X2b1 | X2b | 64 | 52 |
| AED204 | 0.25X | X2b11 | 3 | X2 | X2 | 54 | 48 |
| AED204 | 0.25X | X2b11 | 4 | N1a1b | X4 | 59 | 51 |
| AED204 | 0.25X | X2b11 | 5 | X2b12 | X2b12 | 42 | 36 |
| AED92b | 0.25X | U4a1 | 1 | U4a2d | U2b | 45 | 36 |
| AED92b | 0.25X | U4a1 | 2 | U4a1 | U4a1 | 68 | 62 |
| AED92b | 0.25X | U4a1 | 3 | U4a1 | U4d1 | 58 | 49 |
| AED92b | 0.25X | U4a1 | 4 | U4 | U4 | 66 | 57 |
| AED92b | 0.25X | U4a1 | 5 | U4a1 | U4a1 | 65 | 60 |
| Alh10 | 0.25X | I1 | 1 | I1 | I1 | 46 | 44 |
| Alh10 | 0.25X | I1 | 2 | H27a | B4c1a1a1 | 44 | 35 |
| Alh10 | 0.25X | I1 | 3 | A1 | A1 | 42 | 38 |
| Alh10 | 0.25X | I1 | 4 | I6a | I6a | 61 | 52 |
| Alh10 | 0.25X | I1 | 5 | I1 | I1 | 39 | 36 |
| DA100 | 0.25X | C4b1 | 1 | C4 | C4 | 41 | 39 |
| DA100 | 0.25X | C4b1 | 2 | C4 | C4 | 54 | 51 |
| DA100 | 0.25X | C4b1 | 3 | C4b | C4b | 36 | 35 |
| DA100 | 0.25X | C4b1 | 4 | C4 | C4 | 43 | 40 |
| DA100 | 0.25X | C4b1 | 5 | C4b1 | C4b1 | 34 | 28 |
| DA101 | 0.25X | U5a1b1e | 1 | U5a | U5a | 50 | 47 |
| DA101 | 0.25X | U5a1b1e | 2 | U5a1b1 | U5a1b1 | 49 | 46 |
| DA101 | 0.25X | U5a1b1e | 3 | U5a | U5a | 42 | 38 |
| DA101 | 0.25X | U5a1b1e | 4 | U5a1b1e | U5a1b1e | 48 | 45 |
| DA101 | 0.25X | U5a1b1e | 5 | U6c | H2a1f2 | 43 | 41 |
| DA15 | 0.25X | C4d | 1 | C1 | C1 | 54 | 48 |
| DA15 | 0.25X | C4d | 2 | C4e | C4e | 55 | 49 |
| DA15 | 0.25X | C4d | 3 | C4e | C4e | 58 | 54 |
| DA15 | 0.25X | C4d | 4 | C4d | C4d | 61 | 56 |
| DA15 | 0.25X | C4d | 5 | C4e | C4e | 48 | 45 |
| DA171 | 0.25X | H2a1 | 1 | H3u | H1bd | 57 | 50 |
| DA171 | 0.25X | H2a1 | 2 | M35a | M35a | 59 | 54 |
| DA171 | 0.25X | H2a1 | 3 | U5a2e | U5a2e | 61 | 53 |
| DA171 | 0.25X | H2a1 | 4 | H5p | H5p | 55 | 47 |
| DA171 | 0.25X | H2a1 | 5 | B5 | B4m | 63 | 58 |
| FN2 | 0.25X | H3 | 1 | H24a1 | H24a1 | 49 | 44 |
| FN2 | 0.25X | H3 | 2 | H1aw | L3 | 52 | 48 |
| FN2 | 0.25X | H3 | 3 | H56 | H1 | 53 | 46 |
| FN2 | 0.25X | H3 | 4 | H27c | H27c | 51 | 41 |

Continued on next page

| Sample | Rate | Full_Cov_Pred | Replicate | HG_corr | HG_uncorr | Reads_corr | Reads_uncorr |
|---|---|---|---|---|---|---|---|
| FN2 | 0.25X | H3 | 5 | H27 | H10 | 32 | 25 |
| I10899 | 0.25X | U5b | 1 | U5b1 | U5b1 | 88 | 86 |
| I10899 | 0.25X | U5b | 2 | U5a | U5a | 92 | 91 |
| I10899 | 0.25X | U5b | 3 | L3e2a | L3e2a | 107 | 103 |
| I10899 | 0.25X | U5b | 4 | U5a'b | U5a'b | 82 | 78 |
| I10899 | 0.25X | U5b | 5 | U5b1 | U5b1 | 108 | 104 |
| I11300 | 0.25X | J2a1a1 | 1 | J2a1a1 | J2a1a1 | 80 | 76 |
| I11300 | 0.25X | J2a1a1 | 2 | J2a1a1 | J2a1a1 | 110 | 107 |
| I11300 | 0.25X | J2a1a1 | 3 | J2a1a1 | J2a1a1 | 83 | 80 |
| I11300 | 0.25X | J2a1a1 | 4 | J2a1a1 | J2a1a1 | 103 | 98 |
| I11300 | 0.25X | J2a1a1 | 5 | J2 | J2 | 88 | 84 |
| I7645 | 0.25X | R7b2 | 1 | B4'5 | B4'5 | 35 | 32 |
| I7645 | 0.25X | R7b2 | 2 | B2 | B2 | 61 | 59 |
| I7645 | 0.25X | R7b2 | 3 | T2b22 | T2b22 | 49 | 49 |
| I7645 | 0.25X | R7b2 | 4 | B2 | B2 | 52 | 49 |
| I7645 | 0.25X | R7b2 | 5 | B4'5 | B4'5 | 68 | 64 |
| I7646 | 0.25X | H1e1c | 1 | H1bi | H1bi | 68 | 66 |
| I7646 | 0.25X | H1e1c | 2 | H1e1c | H1e1c | 68 | 63 |
| I7646 | 0.25X | H1e1c | 3 | J1d | J1d | 61 | 54 |
| I7646 | 0.25X | H1e1c | 4 | H1bi | H1bi | 70 | 69 |
| I7646 | 0.25X | H1e1c | 5 | H1 | H1 | 78 | 72 |
| I8132 | 0.25X | D1a2 | 1 | U8a1b | U8a1b | 49 | 47 |
| I8132 | 0.25X | D1a2 | 2 | J1c2f | J1c2f | 42 | 40 |
| I8132 | 0.25X | D1a2 | 3 | B2 | B2 | 40 | 39 |
| I8132 | 0.25X | D1a2 | 4 | H1aq | H1aq | 51 | 48 |
| I8132 | 0.25X | D1a2 | 5 | H1aq | H1aq | 48 | 48 |
| I8569 | 0.25X | H1ah | 1 | H1 | H1 | 64 | 64 |
| I8569 | 0.25X | H1ah | 2 | B4 | B4 | 69 | 66 |
| I8569 | 0.25X | H1ah | 3 | B4'5 | B4'5 | 65 | 65 |
| I8569 | 0.25X | H1ah | 4 | B4 | B4 | 71 | 68 |
| I8569 | 0.25X | H1ah | 5 | H1 | H1 | 63 | 62 |
| NW54 | 0.25X | C4a1a1 | 1 | C4b5 | C1d1 | 56 | 46 |
| NW54 | 0.25X | C4a1a1 | 2 | C1b8a | C1b8a | 45 | 38 |
| NW54 | 0.25X | C4a1a1 | 3 | C4d | C4d | 59 | 47 |
| NW54 | 0.25X | C4a1a1 | 4 | C4a1a1 | C4a1a1 | 65 | 57 |
| NW54 | 0.25X | C4a1a1 | 5 | C4a1a | M18 | 56 | 48 |
| STR266b | 0.25X | J1c5 | 1 | F1 | U4a2h1 | 34 | 29 |
| STR266b | 0.25X | J1c5 | 2 | J1 | J1 | 46 | 41 |
| STR266b | 0.25X | J1c5 | 3 | U5b2a1b | U5b2a1b | 44 | 40 |
| STR266b | 0.25X | J1c5 | 4 | J1c9 | J1c9 | 56 | 51 |
| STR266b | 0.25X | J1c5 | 5 | J1c7 | J1c7 | 59 | 53 |
| STR393b | 0.25X | H5a1 | 1 | H2a1f | H2a1f | 50 | 43 |
| STR393b | 0.25X | H5a1 | 2 | H1c4a1 | H44b | 56 | 42 |
| STR393b | 0.25X | H5a1 | 3 | H5s | H5s | 54 | 41 |
| STR393b | 0.25X | H5a1 | 4 | V25 | V25 | 52 | 43 |
| STR393b | 0.25X | H5a1 | 5 | H3w | H3w | 55 | 44 |
| STR486 | 0.25X | T2b | 1 | T2f5 | T1 | 50 | 43 |
| STR486 | 0.25X | T2b | 2 | T2 | T2 | 50 | 41 |
| STR486 | 0.25X | T2b | 3 | T1 | T1 | 47 | 42 |
| STR486 | 0.25X | T2b | 4 | T2 | T2 | 56 | 49 |

Continued on next page

| Sample | Rate | Full_Cov_Pred | Replicate | HG_corr | HG_uncorr | Reads_corr | Reads_uncorr |
|---|---|---|---|---|---|---|---|
| STR486 | 0.25X | T2b | 5 | T2b | T2b | 56 | 45 |
| STR491 | 0.25X | T2b | 1 | T2b | T2b | 55 | 47 |
| STR491 | 0.25X | T2b | 2 | T2b33 | T2b33 | 42 | 37 |
| STR491 | 0.25X | T2b | 3 | T2b | T2e1a1b1 | 52 | 44 |
| STR491 | 0.25X | T2b | 4 | T2b | T1 | 44 | 36 |
| STR491 | 0.25X | T2b | 5 | T2 | T2 | 46 | 43 |
| Vim2b | 0.25X | H7 | 1 | H1 | H72 | 46 | 38 |
| Vim2b | 0.25X | H7 | 2 | H2a1f1 | H2a1f1 | 47 | 38 |
| Vim2b | 0.25X | H7 | 3 | HV17 | HV17 | 43 | 40 |
| Vim2b | 0.25X | H7 | 4 | H1 | H1 | 47 | 34 |
| Vim2b | 0.25X | H7 | 5 | H1c7 | H1c7 | 42 | 36 |
| AED204 | 0.5X | X2b11 | 1 | X2b | B4 | 99 | 81 |
| AED204 | 0.5X | X2b11 | 2 | X2b6 | X2b6 | 104 | 88 |
| AED204 | 0.5X | X2b11 | 3 | X2b9 | A2e | 115 | 102 |
| AED204 | 0.5X | X2b11 | 4 | X2b4 | X2b4 | 96 | 80 |
| AED204 | 0.5X | X2b11 | 5 | X2b | X2b | 123 | 102 |
| AED92b | 0.5X | U4a1 | 1 | U4a1 | U4a1 | 116 | 103 |
| AED92b | 0.5X | U4a1 | 2 | U4a1d | N22a | 120 | 101 |
| AED92b | 0.5X | U4a1 | 3 | U4a1 | U4a1 | 115 | 101 |
| AED92b | 0.5X | U4a1 | 4 | U4a1 | U4a1 | 124 | 103 |
| AED92b | 0.5X | U4a1 | 5 | U4a1 | U4a1 | 144 | 128 |
| Alh10 | 0.5X | I1 | 1 | I1 | I1 | 84 | 69 |
| Alh10 | 0.5X | I1 | 2 | I1 | I1 | 91 | 72 |
| Alh10 | 0.5X | I1 | 3 | I1 | I1 | 100 | 89 |
| Alh10 | 0.5X | I1 | 4 | I1c | I1 | 97 | 85 |
| Alh10 | 0.5X | I1 | 5 | I1 | I1 | 84 | 76 |
| DA100 | 0.5X | C4b1 | 1 | C4b1 | C4b1 | 100 | 91 |
| DA100 | 0.5X | C4b1 | 2 | C4b5 | C4b5 | 90 | 87 |
| DA100 | 0.5X | C4b1 | 3 | C4 | C4 | 76 | 69 |
| DA100 | 0.5X | C4b1 | 4 | C4b1 | C4b1 | 92 | 84 |
| DA100 | 0.5X | C4b1 | 5 | C4 | C4 | 75 | 68 |
| DA101 | 0.5X | U5a1b1e | 1 | U5a1b1e | U5a1b1e | 90 | 82 |
| DA101 | 0.5X | U5a1b1e | 2 | U5a1b1e | U5a1b1e | 110 | 103 |
| DA101 | 0.5X | U5a1b1e | 3 | U5a2a1a | U5a2a1a | 103 | 94 |
| DA101 | 0.5X | U5a1b1e | 4 | U5a1b1e | U5a1b1e | 103 | 97 |
| DA101 | 0.5X | U5a1b1e | 5 | U5a1 | U5a1 | 95 | 84 |
| DA15 | 0.5X | C4d | 1 | C1 | C1 | 110 | 98 |
| DA15 | 0.5X | C4d | 2 | C1 | C1 | 120 | 112 |
| DA15 | 0.5X | C4d | 3 | C4 | C4 | 120 | 109 |
| DA15 | 0.5X | C4d | 4 | C4 | C4 | 132 | 114 |
| DA15 | 0.5X | C4d | 5 | C4d | C4d | 98 | 86 |
| DA171 | 0.5X | H2a1 | 1 | H2a1 | H2a1 | 116 | 102 |
| DA171 | 0.5X | H2a1 | 2 | H107 | H1an2 | 118 | 111 |
| DA171 | 0.5X | H2a1 | 3 | H5p | H5p | 113 | 98 |
| DA171 | 0.5X | H2a1 | 4 | H1an2 | H1an2 | 104 | 96 |
| DA171 | 0.5X | H2a1 | 5 | H1an2 | H1an2 | 114 | 104 |
| FN2 | 0.5X | H3 | 1 | H48 | H48 | 114 | 89 |
| FN2 | 0.5X | H3 | 2 | H104 | H104 | 117 | 95 |
| FN2 | 0.5X | H3 | 3 | H5a9 | H5a9 | 104 | 90 |
| FN2 | 0.5X | H3 | 4 | H16e | H16e | 116 | 88 |

Continued on next page

| Sample | Rate | Full_Cov_Pred | Replicate | HG_corr | HG_uncorr | Reads_corr | Reads_uncorr |
|---|---|---|---|---|---|---|---|
| FN2 | 0.5X | H3 | 5 | H3at1 | H105 | 90 | 80 |
| I10899 | 0.5X | U5b | 1 | U5b1 | U5b1 | 210 | 207 |
| I10899 | 0.5X | U5b | 2 | U5b1 | U5b1 | 161 | 157 |
| I10899 | 0.5X | U5b | 3 | U5b1 | U5b1 | 201 | 197 |
| I10899 | 0.5X | U5b | 4 | U5b1h | U5b1h | 172 | 169 |
| I10899 | 0.5X | U5b | 5 | U5b | U5b | 172 | 169 |
| I11300 | 0.5X | J2a1a1 | 1 | J2a1a | J2a1a | 175 | 169 |
| I11300 | 0.5X | J2a1a1 | 2 | J2a1a | J2a1 | 186 | 179 |
| I11300 | 0.5X | J2a1a1 | 3 | J2a1a1 | J2a1a | 189 | 186 |
| I11300 | 0.5X | J2a1a1 | 4 | J2a1a1 | J2a1a1 | 181 | 175 |
| I11300 | 0.5X | J2a1a1 | 5 | J2a1a1 | J2a1a1 | 182 | 174 |
| I7645 | 0.5X | R7b2 | 1 | F | F | 105 | 102 |
| I7645 | 0.5X | R7b2 | 2 | HV0d | HV0d | 108 | 103 |
| I7645 | 0.5X | R7b2 | 3 | HV0d | HV0d | 103 | 96 |
| I7645 | 0.5X | R7b2 | 4 | F | F | 114 | 107 |
| I7645 | 0.5X | R7b2 | 5 | HV0d | HV0d | 110 | 105 |
| I7646 | 0.5X | H1e1c | 1 | H1bi | H1bi | 157 | 147 |
| I7646 | 0.5X | H1e1c | 2 | H1e | H1e | 139 | 133 |
| I7646 | 0.5X | H1e1c | 3 | H58 | H58 | 152 | 141 |
| I7646 | 0.5X | H1e1c | 4 | H1e1c | H1e1c | 167 | 162 |
| I7646 | 0.5X | H1e1c | 5 | H1 | H1 | 148 | 139 |
| I8132 | 0.5X | D1a2 | 1 | H1 | H1 | 102 | 97 |
| I8132 | 0.5X | D1a2 | 2 | H1 | H1 | 85 | 79 |
| I8132 | 0.5X | D1a2 | 3 | H1aq | H1aq | 88 | 87 |
| I8132 | 0.5X | D1a2 | 4 | H1aq | H1aq | 84 | 77 |
| I8132 | 0.5X | D1a2 | 5 | H1aq | H1aq | 85 | 80 |
| I8569 | 0.5X | H1ah | 1 | H1ah | H1ah | 130 | 126 |
| I8569 | 0.5X | H1ah | 2 | H11a6 | H11a6 | 155 | 151 |
| I8569 | 0.5X | H1ah | 3 | H3r | H3r | 140 | 133 |
| I8569 | 0.5X | H1ah | 4 | H1ah | H1ah | 130 | 125 |
| I8569 | 0.5X | H1ah | 5 | H1ah | H1ah | 160 | 158 |
| NW54 | 0.5X | C4a1a1 | 1 | C4a1a | C4a1a | 92 | 82 |
| NW54 | 0.5X | C4a1a1 | 2 | C4a1a | C4a1a | 97 | 84 |
| NW54 | 0.5X | C4a1a1 | 3 | C4a1a1 | C4a1a1 | 93 | 86 |
| NW54 | 0.5X | C4a1a1 | 4 | C4a1 | C4a1a | 102 | 86 |
| NW54 | 0.5X | C4a1a1 | 5 | C4a1 | C4a1a1 | 115 | 94 |
| STR266b | 0.5X | J1c5 | 1 | J1c5 | J1c5 | 84 | 76 |
| STR266b | 0.5X | J1c5 | 2 | J1c | J1c | 91 | 80 |
| STR266b | 0.5X | J1c5 | 3 | J1c2a3 | J1c2a3 | 78 | 70 |
| STR266b | 0.5X | J1c5 | 4 | J1c1b1 | J1c1b1 | 85 | 74 |
| STR266b | 0.5X | J1c5 | 5 | J1d | J1c | 100 | 91 |
| STR393b | 0.5X | H5a1 | 1 | H5a1 | H5a1 | 121 | 104 |
| STR393b | 0.5X | H5a1 | 2 | H1bc | B4a1a1u | 111 | 93 |
| STR393b | 0.5X | H5a1 | 3 | H5a1p | H5a1p | 101 | 80 |
| STR393b | 0.5X | H5a1 | 4 | H1ac | H1ac | 98 | 78 |
| STR393b | 0.5X | H5a1 | 5 | H1ab | H1ab | 116 | 98 |
| STR486 | 0.5X | T2b | 1 | T2 | T2 | 90 | 78 |
| STR486 | 0.5X | T2b | 2 | T2 | T2 | 96 | 81 |
| STR486 | 0.5X | T2b | 3 | T2 | T2 | 96 | 75 |
| STR486 | 0.5X | T2b | 4 | T2b | T2b | 103 | 89 |

Continued on next page

| Sample | Rate | Full_Cov_Pred | Replicate | HG_corr | HG_uncorr | Reads_corr | Reads_uncorr |
|---|---|---|---|---|---|---|---|
| STR486 | 0.5X | T2b | 5 | T2b | T2b | 100 | 90 |
| STR491 | 0.5X | T2b | 1 | T2b33 | T2b33 | 90 | 75 |
| STR491 | 0.5X | T2b | 2 | T2b | T2b | 86 | 73 |
| STR491 | 0.5X | T2b | 3 | T2b | T2b | 77 | 67 |
| STR491 | 0.5X | T2b | 4 | T2b | T2 | 109 | 91 |
| STR491 | 0.5X | T2b | 5 | T2b | T2b | 88 | 72 |
| Vim2b | 0.5X | H7 | 1 | H27d | H41 | 96 | 81 |
| Vim2b | 0.5X | H7 | 2 | H56a | H2a | 98 | 89 |
| Vim2b | 0.5X | H7 | 3 | H5a1c | H8c | 113 | 104 |
| Vim2b | 0.5X | H7 | 4 | HV6a | HV6a | 95 | 85 |
| Vim2b | 0.5X | H7 | 5 | H30a | H30a | 103 | 92 |
| AED204 | 1.0X | X2b11 | 1 | X2b6 | X2b6 | 199 | 175 |
| AED204 | 1.0X | X2b11 | 2 | X2b | X2b | 221 | 191 |
| AED204 | 1.0X | X2b11 | 3 | X2b3 | X2b3 | 214 | 178 |
| AED204 | 1.0X | X2b11 | 4 | X2b | X2b | 179 | 149 |
| AED204 | 1.0X | X2b11 | 5 | X2b | X2b | 212 | 187 |
| AED92b | 1.0X | U4a1 | 1 | U4a1 | U4a1e | 248 | 205 |
| AED92b | 1.0X | U4a1 | 2 | U4a1d | U4a1 | 221 | 185 |
| AED92b | 1.0X | U4a1 | 3 | U4a1 | U4a1 | 240 | 205 |
| AED92b | 1.0X | U4a1 | 4 | U4a1 | U4a1 | 259 | 227 |
| AED92b | 1.0X | U4a1 | 5 | U4a1a | U4a1 | 246 | 207 |
| Alh10 | 1.0X | I1 | 1 | I1 | I1 | 188 | 165 |
| Alh10 | 1.0X | I1 | 2 | I1d | I1d | 185 | 160 |
| Alh10 | 1.0X | I1 | 3 | I1 | I1 | 192 | 162 |
| Alh10 | 1.0X | I1 | 4 | I2e | I1 | 190 | 173 |
| Alh10 | 1.0X | I1 | 5 | I1 | I1 | 183 | 166 |
| DA100 | 1.0X | C4b1 | 1 | C4b1 | C4b1 | 199 | 183 |
| DA100 | 1.0X | C4b1 | 2 | C4b | C4b | 177 | 161 |
| DA100 | 1.0X | C4b1 | 3 | C4b1 | C4b1 | 199 | 184 |
| DA100 | 1.0X | C4b1 | 4 | C4b1 | C4b1 | 227 | 218 |
| DA100 | 1.0X | C4b1 | 5 | C4b1 | C4b1 | 203 | 190 |
| DA101 | 1.0X | U5a1b1e | 1 | U5a1b1e | U5a1b1e | 195 | 184 |
| DA101 | 1.0X | U5a1b1e | 2 | U5a1b1e | U5a1b1e | 189 | 175 |
| DA101 | 1.0X | U5a1b1e | 3 | U5a1b1e | U5a1b1e | 176 | 164 |
| DA101 | 1.0X | U5a1b1e | 4 | U5a1b1e | U5a1b1e | 181 | 168 |
| DA101 | 1.0X | U5a1b1e | 5 | U5a1b1c1 | U5a1b1e | 173 | 163 |
| DA15 | 1.0X | C4d | 1 | C4e | C4e | 211 | 185 |
| DA15 | 1.0X | C4d | 2 | C4e | C4 | 217 | 201 |
| DA15 | 1.0X | C4d | 3 | C4 | C4 | 230 | 202 |
| DA15 | 1.0X | C4d | 4 | C4 | C4e | 223 | 201 |
| DA15 | 1.0X | C4d | 5 | C4e | C4e | 233 | 204 |
| DA171 | 1.0X | H2a1 | 1 | H3r | H3r | 219 | 204 |
| DA171 | 1.0X | H2a1 | 2 | H2a1 | H2a1 | 213 | 199 |
| DA171 | 1.0X | H2a1 | 3 | H2a1d | H2a1 | 237 | 217 |
| DA171 | 1.0X | H2a1 | 4 | H2a1 | H76a | 226 | 210 |
| DA171 | 1.0X | H2a1 | 5 | H2a1 | H2a1 | 221 | 198 |
| FN2 | 1.0X | H3 | 1 | H3ar | H3ar | 208 | 183 |
| FN2 | 1.0X | H3 | 2 | H3aq | H7e | 218 | 190 |
| FN2 | 1.0X | H3 | 3 | H7b | H3z1 | 246 | 201 |
| FN2 | 1.0X | H3 | 4 | H25 | H50 | 210 | 171 |

Continued on next page

| Sample | Rate | Full_Cov_Pred | Replicate | HG_corr | HG_uncorr | Reads_corr | Reads_uncorr |
|---|---|---|---|---|---|---|---|
| FN2 | 1.0X | H3 | 5 | H66 | H66 | 222 | 194 |
| I10899 | 1.0X | U5b | 1 | U5b1 | U5b1c | 367 | 356 |
| I10899 | 1.0X | U5b | 2 | U5b | U5b | 388 | 375 |
| I10899 | 1.0X | U5b | 3 | U5b1 | U5b1 | 367 | 350 |
| I10899 | 1.0X | U5b | 4 | U5b | U5b | 402 | 394 |
| I10899 | 1.0X | U5b | 5 | U5b1 | U5b1 | 357 | 347 |
| I11300 | 1.0X | J2a1a1 | 1 | J2a1a1 | J2a1a1 | 362 | 354 |
| I11300 | 1.0X | J2a1a1 | 2 | J2a1a1 | J2a1a1 | 325 | 320 |
| I11300 | 1.0X | J2a1a1 | 3 | J2a1a1 | J2a1a1 | 361 | 347 |
| I11300 | 1.0X | J2a1a1 | 4 | J2a1a1 | J2a1a1 | 364 | 348 |
| I11300 | 1.0X | J2a1a1 | 5 | J2a1a1 | J2a1a1 | 370 | 353 |
| I7645 | 1.0X | R7b2 | 1 | HV0d | HV0d | 214 | 205 |
| I7645 | 1.0X | R7b2 | 2 | B4'5 | B4'5 | 200 | 185 |
| I7645 | 1.0X | R7b2 | 3 | HV0d | HV0d | 199 | 187 |
| I7645 | 1.0X | R7b2 | 4 | HV0d | HV0d | 208 | 198 |
| I7645 | 1.0X | R7b2 | 5 | H10 | H10 | 202 | 196 |
| I7646 | 1.0X | H1e1c | 1 | H1e1c | H1e1c | 305 | 289 |
| I7646 | 1.0X | H1e1c | 2 | H1bi | H1bi | 297 | 284 |
| I7646 | 1.0X | H1e1c | 3 | H1bi | H1bi | 287 | 274 |
| I7646 | 1.0X | H1e1c | 4 | H1bi | H1bi | 289 | 272 |
| I7646 | 1.0X | H1e1c | 5 | H1bi | H1bi | 297 | 283 |
| I8132 | 1.0X | D1a2 | 1 | H1aq | H1aq | 186 | 177 |
| I8132 | 1.0X | D1a2 | 2 | H1aq | H1aq | 196 | 186 |
| I8132 | 1.0X | D1a2 | 3 | H1aq | H1aq | 180 | 170 |
| I8132 | 1.0X | D1a2 | 4 | H1aq | H1aq | 172 | 164 |
| I8132 | 1.0X | D1a2 | 5 | H1aq | H1aq | 186 | 175 |
| I8569 | 1.0X | H1ah | 1 | H1 | H1 | 249 | 240 |
| I8569 | 1.0X | H1ah | 2 | H1 | H1 | 292 | 285 |
| I8569 | 1.0X | H1ah | 3 | H1ah | H1ah | 275 | 267 |
| I8569 | 1.0X | H1ah | 4 | H1ah | H1ah | 254 | 246 |
| I8569 | 1.0X | H1ah | 5 | H1 | H1 | 281 | 271 |
| NW54 | 1.0X | C4a1a1 | 1 | C4a1a | C4a1a | 234 | 189 |
| NW54 | 1.0X | C4a1a1 | 2 | C4a1a | C4a1a | 205 | 166 |
| NW54 | 1.0X | C4a1a1 | 3 | C4a1a | C4a1a | 195 | 177 |
| NW54 | 1.0X | C4a1a1 | 4 | C4a1a1 | C4a1a1 | 238 | 206 |
| NW54 | 1.0X | C4a1a1 | 5 | C4a1a1 | C4a1a1 | 221 | 186 |
| STR266b | 1.0X | J1c5 | 1 | J1c | J1c | 167 | 153 |
| STR266b | 1.0X | J1c5 | 2 | J1c9 | J1c9 | 161 | 143 |
| STR266b | 1.0X | J1c5 | 3 | J1c4c | J1c4c | 181 | 159 |
| STR266b | 1.0X | J1c5 | 4 | J1c9 | J1c9 | 173 | 154 |
| STR266b | 1.0X | J1c5 | 5 | J1 | J1 | 176 | 158 |
| STR393b | 1.0X | H5a1 | 1 | H5a | H5a | 232 | 194 |
| STR393b | 1.0X | H5a1 | 2 | H5a1 | H5a1 | 218 | 184 |
| STR393b | 1.0X | H5a1 | 3 | H5a1 | H5a1 | 197 | 169 |
| STR393b | 1.0X | H5a1 | 4 | H5c | H5c | 247 | 209 |
| STR393b | 1.0X | H5a1 | 5 | H5a1 | H5a1 | 225 | 179 |
| STR486 | 1.0X | T2b | 1 | T2 | T2b33 | 183 | 157 |
| STR486 | 1.0X | T2b | 2 | T2a3 | T2a3 | 209 | 183 |
| STR486 | 1.0X | T2b | 3 | T2b | T2b | 197 | 172 |
| STR486 | 1.0X | T2b | 4 | T2a3 | T2a3 | 170 | 152 |

Continued on next page

| Sample | Rate | Full_Cov_Pred | Replicate | HG_corr | HG_uncorr | Reads_corr | Reads_uncorr |
|---|---|---|---|---|---|---|---|
| STR486 | 1.0X | T2b | 5 | T2b17 | T2b17 | 207 | 192 |
| STR491 | 1.0X | T2b | 1 | T2b | T2b | 177 | 154 |
| STR491 | 1.0X | T2b | 2 | T2b | T2b | 176 | 145 |
| STR491 | 1.0X | T2b | 3 | T2b | T2b | 192 | 157 |
| STR491 | 1.0X | T2b | 4 | T2b | T2b | 187 | 154 |
| STR491 | 1.0X | T2b | 5 | T2b | T2b | 158 | 134 |
| Vim2b | 1.0X | H7 | 1 | H1 | H1 | 209 | 191 |
| Vim2b | 1.0X | H7 | 2 | R0 | R0 | 194 | 168 |
| Vim2b | 1.0X | H7 | 3 | H53 | H53 | 185 | 161 |
| Vim2b | 1.0X | H7 | 4 | H57 | H57 | 187 | 159 |
| Vim2b | 1.0X | H7 | 5 | H102 | H102 | 210 | 177 |
| AED204 | 2.0X | X2b11 | 1 | X2b1 | X2b1 | 391 | 331 |
| AED204 | 2.0X | X2b11 | 2 | X2b7 | X2b7 | 392 | 341 |
| AED204 | 2.0X | X2b11 | 3 | X2b13 | X2b7 | 411 | 346 |
| AED204 | 2.0X | X2b11 | 4 | X2b7 | X2b1 | 433 | 369 |
| AED204 | 2.0X | X2b11 | 5 | X2b3 | X2b3 | 411 | 358 |
| AED92b | 2.0X | U4a1 | 1 | U4a1a | U4a1 | 458 | 397 |
| AED92b | 2.0X | U4a1 | 2 | U4a1 | U4a1 | 511 | 436 |
| AED92b | 2.0X | U4a1 | 3 | U4a1a | U4a1e | 500 | 440 |
| AED92b | 2.0X | U4a1 | 4 | U4a1 | U4a1 | 490 | 427 |
| AED92b | 2.0X | U4a1 | 5 | U4a1a | U4a1 | 515 | 439 |
| Alh10 | 2.0X | I1 | 1 | I1 | I1 | 343 | 299 |
| Alh10 | 2.0X | I1 | 2 | I1 | I1 | 363 | 322 |
| Alh10 | 2.0X | I1 | 3 | I1 | I1 | 383 | 330 |
| Alh10 | 2.0X | I1 | 4 | I1 | I1 | 382 | 331 |
| Alh10 | 2.0X | I1 | 5 | I1 | I1 | 375 | 315 |
| DA100 | 2.0X | C4b1 | 1 | C4b1 | C4b1 | 382 | 358 |
| DA100 | 2.0X | C4b1 | 2 | C4b1 | C4b1 | 381 | 358 |
| DA100 | 2.0X | C4b1 | 3 | C4b1 | C4b1 | 396 | 358 |
| DA100 | 2.0X | C4b1 | 4 | C4b1 | C4b1 | 371 | 348 |
| DA100 | 2.0X | C4b1 | 5 | C4b1 | C4b1 | 443 | 410 |
| DA101 | 2.0X | U5a1b1e | 1 | U5a1b1a1 | U5a1b1a1 | 394 | 363 |
| DA101 | 2.0X | U5a1b1e | 2 | U5a1b1e | U5a1b1e | 349 | 321 |
| DA101 | 2.0X | U5a1b1e | 3 | U5a1b1e | U5a1b1e | 405 | 375 |
| DA101 | 2.0X | U5a1b1e | 4 | U5a1b1e | U5a1b1e | 411 | 382 |
| DA101 | 2.0X | U5a1b1e | 5 | U5a1b1e | U5a1b1e | 398 | 372 |
| DA15 | 2.0X | C4d | 1 | C4d | C4d | 468 | 427 |
| DA15 | 2.0X | C4d | 2 | C4 | C4 | 412 | 358 |
| DA15 | 2.0X | C4d | 3 | C4d | C4d | 409 | 370 |
| DA15 | 2.0X | C4d | 4 | C4d | C4d | 475 | 420 |
| DA15 | 2.0X | C4d | 5 | C4d | C4d | 465 | 425 |
| DA171 | 2.0X | H2a1 | 1 | H2a1 | H2a1 | 431 | 398 |
| DA171 | 2.0X | H2a1 | 2 | H2a1 | H3i | 493 | 449 |
| DA171 | 2.0X | H2a1 | 3 | H76a | H76a | 440 | 397 |
| DA171 | 2.0X | H2a1 | 4 | H2a1 | H2a1 | 453 | 415 |
| DA171 | 2.0X | H2a1 | 5 | H2a1 | H2a1 | 453 | 413 |
| FN2 | 2.0X | H3 | 1 | H3 | H3 | 457 | 401 |
| FN2 | 2.0X | H3 | 2 | H2 | H2 | 403 | 348 |
| FN2 | 2.0X | H3 | 3 | H104 | H58 | 414 | 357 |
| FN2 | 2.0X | H3 | 4 | H66 | H66 | 414 | 361 |

Continued on next page

| Sample | Rate | Full_Cov_Pred | Replicate | HG_corr | HG_uncorr | Reads_corr | Reads_uncorr |
|---|---|---|---|---|---|---|---|
| FN2 | 2.0X | H3 | 5 | H3 | H3 | 465 | 390 |
| I10899 | 2.0X | U5b | 1 | U5b1 | U5b1 | 731 | 713 |
| I10899 | 2.0X | U5b | 2 | U5b1 | U5b1 | 785 | 753 |
| I10899 | 2.0X | U5b | 3 | U5b1 | U5b1 | 691 | 657 |
| I10899 | 2.0X | U5b | 4 | U5b1 | U5b1 | 732 | 705 |
| I10899 | 2.0X | U5b | 5 | U5b1 | U5b1 | 729 | 707 |
| I11300 | 2.0X | J2a1a1 | 1 | J2a1a1 | J2a1a1 | 694 | 673 |
| I11300 | 2.0X | J2a1a1 | 2 | J2a1a1 | J2a1a1 | 690 | 662 |
| I11300 | 2.0X | J2a1a1 | 3 | J2a1a1 | J2a1a1 | 708 | 690 |
| I11300 | 2.0X | J2a1a1 | 4 | J2a1a1 | J2a1a1 | 699 | 670 |
| I11300 | 2.0X | J2a1a1 | 5 | J2a1a1 | J2a1a1 | 696 | 670 |
| I7645 | 2.0X | R7b2 | 1 | F | F | 74 | 70 |
| I7645 | 2.0X | R7b2 | 2 | B2 | B2 | 79 | 74 |
| I7645 | 2.0X | R7b2 | 3 | HV0d | HV0d | 79 | 77 |
| I7645 | 2.0X | R7b2 | 4 | H1 | H1 | 93 | 90 |
| I7645 | 2.0X | R7b2 | 5 | CZ | CZ | 73 | 69 |
| I7646 | 2.0X | H1e1c | 1 | H1e1c | H1e1c | 549 | 520 |
| I7646 | 2.0X | H1e1c | 2 | H1e1c | H1e1c | 544 | 513 |
| I7646 | 2.0X | H1e1c | 3 | H1e1c | H1e1c | 547 | 513 |
| I7646 | 2.0X | H1e1c | 4 | H1bi | H1bi | 555 | 530 |
| I7646 | 2.0X | H1e1c | 5 | H1e1c | H1bi | 562 | 534 |
| I8132 | 2.0X | D1a2 | 1 | H1aq | H1aq | 89 | 84 |
| I8132 | 2.0X | D1a2 | 2 | H1aq | H1aq | 105 | 99 |
| I8132 | 2.0X | D1a2 | 3 | H1aq | H1aq | 97 | 91 |
| I8132 | 2.0X | D1a2 | 4 | H1aq | H1aq | 103 | 97 |
| I8132 | 2.0X | D1a2 | 5 | H1 | H1 | 105 | 100 |
| I8569 | 2.0X | H1ah | 1 | H1ah | H1ah | 520 | 503 |
| I8569 | 2.0X | H1ah | 2 | H1ah | H1ah | 522 | 502 |
| I8569 | 2.0X | H1ah | 3 | H1ah | H1ah | 532 | 514 |
| I8569 | 2.0X | H1ah | 4 | H1ah | H1ah | 525 | 508 |
| I8569 | 2.0X | H1ah | 5 | H1ah | H1ah | 538 | 527 |
| NW54 | 2.0X | C4a1a1 | 1 | C4a1a1 | C4a1a1 | 456 | 379 |
| NW54 | 2.0X | C4a1a1 | 2 | C4a1a1 | C4a1a1 | 445 | 378 |
| NW54 | 2.0X | C4a1a1 | 3 | C4a1a | C4a1a | 452 | 385 |
| NW54 | 2.0X | C4a1a1 | 4 | C4a1a1 | C4a1a1 | 419 | 363 |
| NW54 | 2.0X | C4a1a1 | 5 | C4a1a1 | C4a1a1 | 417 | 364 |
| STR266b | 2.0X | J1c5 | 1 | J1c3g | J1c3g | 355 | 311 |
| STR266b | 2.0X | J1c5 | 2 | J1c | J1c | 400 | 355 |
| STR266b | 2.0X | J1c5 | 3 | J1c | J1c | 356 | 315 |
| STR266b | 2.0X | J1c5 | 4 | J1c5 | J1c5 | 347 | 310 |
| STR266b | 2.0X | J1c5 | 5 | J1c | J1c | 363 | 319 |
| STR393b | 2.0X | H5a1 | 1 | H5v | H5a | 450 | 384 |
| STR393b | 2.0X | H5a1 | 2 | H5a1 | H5a1 | 430 | 359 |
| STR393b | 2.0X | H5a1 | 3 | H5a1 | H5a1 | 428 | 352 |
| STR393b | 2.0X | H5a1 | 4 | H5a1 | H5a1 | 465 | 385 |
| STR393b | 2.0X | H5a1 | 5 | H5a1 | H5a8 | 428 | 370 |
| STR486 | 2.0X | T2b | 1 | T2b | T2 | 387 | 334 |
| STR486 | 2.0X | T2b | 2 | T2b | T2b | 387 | 343 |
| STR486 | 2.0X | T2b | 3 | T2i | T2i | 374 | 316 |
| STR486 | 2.0X | T2b | 4 | T2b | T2b | 388 | 336 |

Continued on next page

| Sample | Rate | Full_Cov_Pred | Replicate | HG_corr | HG_uncorr | Reads_corr | Reads_uncorr |
|---|---|---|---|---|---|---|---|
| STR486 | 2.0X | T2b | 5 | T2b | T2b | 404 | 353 |
| STR491 | 2.0X | T2b | 1 | T2b | T2b | 346 | 302 |
| STR491 | 2.0X | T2b | 2 | T2b | T2b | 333 | 292 |
| STR491 | 2.0X | T2b | 3 | T2b | T2b | 364 | 299 |
| STR491 | 2.0X | T2b | 4 | T2b | T2b | 377 | 326 |
| STR491 | 2.0X | T2b | 5 | T2b | T2b | 374 | 313 |
| Vim2b | 2.0X | H7 | 1 | R0 | R0 | 404 | 360 |
| Vim2b | 2.0X | H7 | 2 | H105 | H105 | 395 | 356 |
| Vim2b | 2.0X | H7 | 3 | H7 | H7 | 401 | 354 |
| Vim2b | 2.0X | H7 | 4 | H3am | H3am | 381 | 334 |
| Vim2b | 2.0X | H7 | 5 | HV | HV | 384 | 335 |

#### G.2 euka Experiment

##### G.2.1 Table of Detected Reads Per Taxon

| *j* | Arc *tiinae* | Bov *idae* | Caly *tratae* | Colu *broidea* | Cryp *rhynchinae* | Lamp *ridae* | Lyco *soidea* | Mus | Myo *tis* | Rat *tus* | Urs *idae* | Xeri *inae* |
|---|---|---|---|---|---|---|---|---|---|---|---|---|
| 0.0 | 1346 | 157 | 4300 | 891 | 185 | 2518 | 506 | 216 | 351 | 455 | 1241 | 152 |
| 0.1 | 1343 | 148 | 4293 | 876 | 185 | 2516 | 506 | 212 | 347 | 445 | 1209 | 148 |
| 0.2 | 1342 | 145 | 4271 | 875 | 185 | 2511 | 505 | 208 | 344 | 441 | 1203 | 147 |
| 0.3 | 1341 | 143 | 4257 | 866 | 185 | 2508 | 505 | 205 | 337 | 434 | 1193 | 146 |
| 0.4 | 1340 | 141 | 4247 | 860 | 185 | 2499 | 503 | 205 | 331 | 430 | 1179 | 144 |
| 0.5 | 1331 | 140 | 4222 | 855 | 184 | 2489 | 502 | 203 | 326 | 426 | 1173 | 142 |
| 0.6 | 1326 | 138 | 4203 | 851 | 183 | 2485 | 501 | 199 | 326 | 422 | 1167 | 142 |
| 0.7 | 1323 | 134 | 4195 | 842 | 182 | 2476 | 501 | 197 | 324 | 416 | 1153 | 142 |
| 0.8 | 1316 | 132 | 4178 | 826 | 181 | 2460 | 499 | 191 | 323 | 410 | 1132 | 142 |
| 0.9 | 1302 | 131 | 4141 | 817 | 178 | 2434 | 498 | 188 | 319 | 409 | 1125 | 142 |
| 1.0 | 1237 | 132 | 3980 | 813 | 177 | 2380 | 495 | 185 | 316 | 409 | 1114 | 143 |
| *Unc* | 1237 | 132 | 3980 | 813 | 177 | 2380 | 495 | 185 | 316 | 409 | 1113 | 143 |

Table 9: Reads Detected per Taxon on the euka experiment simulated metagenomic environmental aDNA sample with SAFARI as aligner at varying posterior threshold values. Note that at posterior threshold of 1.0 SAFARI and giraffe are identical. As the posterior threshold lowers more deaminated seeds are rescued and aligned, leading to an increase in detected reads across the board.

##### G.2.2 Detected Reads Per Taxon by SAFARI Posterior Threshold

Below (Figure 5) is simply a pictorial representation of the above table, showing how number of detected reads depends on the user-defined posterior threshold for the SAFARI RYmer seed filter. The horizontal dashed line at $y = 100$ shows the minimum number of reads required by euka to consider a taxon as detected in the sample.

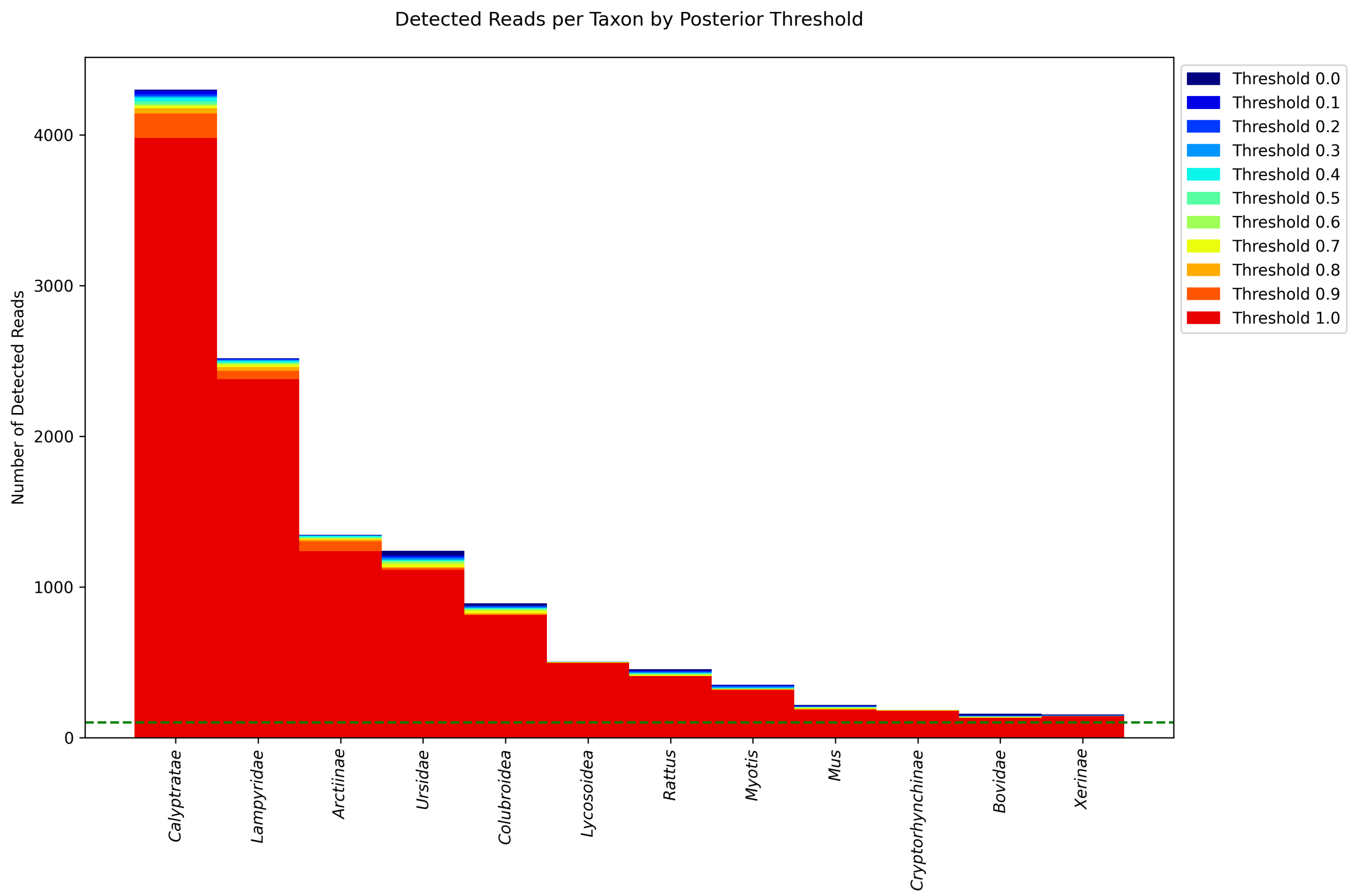

Figure 5: Number of detected reads per taxon as a function of the user-defined posterior threshold (flag `-j`) for the `SAFARI` RYmer seed filter. As the threshold decreases, more reads per taxon are detected. At a threshold of 1.0 the number of reads per taxon is identical to the results from the standard `vg giraffe`, as expected. The dashed horizontal line indicates the minimum number of fragments required to consider a taxon detected.

#### G.3 Linear Alignment Experiment

##### G.3.1 Selected Alignment Statistics

Table 10: Raw counts of mapped reads by provenance in the surjected linear alignment experiment for the dataset of *in silico* simulations following the empirically-derived fragment length distribution from Vindija Cave, Croatia under high damage rates. $NuMT$ is the number of mapped NuMT reads. For readability, only samples for a single subsampling rate (0.9) are shown. Counts are reported as aligned/total. Note that $k = 29, w = 11$ are the default parameters.

| $k$ | $w$ | Bacterial | | NuMTs | | Mitochondrial | |
|---|---|---|---|---|---|---|---|
| | | `SAF` | `gir` | `SAF` | `gir` | `SAF` | `gir` |
| 10 | 2 | 51/90235 | 36/90235 | 89/360 | 88/360 | 269527/270163 | 269425/270163 |
| 10 | 5 | 14/90235 | 10/90235 | 83/360 | 83/360 | 269045/270163 | 268908/270163 |
| 10 | 7 | 8/90235 | 7/90235 | 81/360 | 81/360 | 268319/270163 | 268130/270163 |
| 10 | 8 | 7/90235 | 6/90235 | 79/360 | 79/360 | 267792/270163 | 267589/270163 |
| 10 | 10 | 7/90235 | 6/90235 | 72/360 | 72/360 | 266372/270163 | 266118/270163 |
| 10 | 15 | 3/90235 | 3/90235 | 65/360 | 65/360 | 258594/270163 | 258273/270163 |
| 11 | 6 | 5/90235 | 5/90235 | 75/360 | 73/360 | 267889/270163 | 267640/270163 |
| 11 | 7 | 4/90235 | 4/90235 | 72/360 | 71/360 | 267268/270163 | 266991/270163 |
| 12 | 3 | 3/90235 | 3/90235 | 80/360 | 77/360 | 268526/270163 | 268211/270163 |
| 12 | 4 | 3/90235 | 3/90235 | 79/360 | 75/360 | 268007/270163 | 267703/270163 |
| 12 | 5 | 3/90235 | 3/90235 | 76/360 | 74/360 | 267348/270163 | 267044/270163 |
| 12 | 6 | 3/90235 | 3/90235 | 75/360 | 74/360 | 266611/270163 | 266274/270163 |
| 12 | 7 | 3/90235 | 3/90235 | 71/360 | 70/360 | 265709/270163 | 265395/270163 |
| 15 | 5 | 1/90235 | 1/90235 | 54/360 | 54/360 | 260773/270163 | 260175/270163 |
| 15 | 10 | 0/90235 | 0/90235 | 45/360 | 45/360 | 248752/270163 | 248314/270163 |
| 15 | 15 | 0/90235 | 0/90235 | 33/360 | 33/360 | 231642/270163 | 231284/270163 |
| 18 | 8 | 0/90235 | 0/90235 | 36/360 | 36/360 | 236890/270163 | 236358/270163 |
| 20 | 2 | 0/90235 | 0/90235 | 40/360 | 35/360 | 244702/270163 | 243564/270163 |
| 20 | 5 | 0/90235 | 0/90235 | 29/360 | 28/360 | 233688/270163 | 233095/270163 |
| 20 | 10 | 0/90235 | 0/90235 | 19/360 | 19/360 | 213783/270163 | 213541/270163 |
| 20 | 15 | 0/90235 | 0/90235 | 14/360 | 14/360 | 194248/270163 | 194051/270163 |
| 29 | 11 | 0/90235 | 0/90235 | 8/360 | 5/360 | 138471/270163 | 123588/270163 |
| 30 | 2 | 0/90235 | 0/90235 | 23/360 | 6/360 | 247983/270163 | 166057/270163 |
| 30 | 5 | 0/90235 | 0/90235 | 9/360 | 5/360 | 200888/270163 | 154073/270163 |
| 30 | 10 | 0/90235 | 0/90235 | 6/360 | 5/360 | 138022/270163 | 120769/270163 |
| 30 | 15 | 0/90235 | 0/90235 | 6/360 | 5/360 | 95669/270163 | 88174/270163 |
| **SHRiMP** | | | | | | | |
| | | 0/90235 | | 86/360 | | 269223/270163 | |
| **BWA-MEM** | | | | | | | |
| | | 0/90235 | | 27/360 | | 237299/270163 | |
| **BWA aln** | | | | | | | |
| | | 0/90235 | | 17/360 | | 228817/270163 | |
| **BWA aln (anc)** | | | | | | | |
| | | 0/90235 | | 19/360 | | 251049/270163 | |
| **Bowtie2** | | | | | | | |
| | | 0/90235 | | 15/360 | | 206858/270163 | |

##### G.3.2 Sensitivity vs. Specificity Plots

Figures 6b and 6a show sensitivity vs. specificity of the two pangenome aligners `SAFARI` and `vg giraffe` on the Chagyrskaya Cave distribution datasets for high-damage and zero-damage samples, respectively. The indexing parameters $(k, w) = (10, 2)$ were selected to (individually) optimize median sensitivity. It is evident that sensitivity benefits from `SAFARI`'s rescue of alignments containing deaminated seeds when samples are highly damaged, at the expense of a drop in specificity.

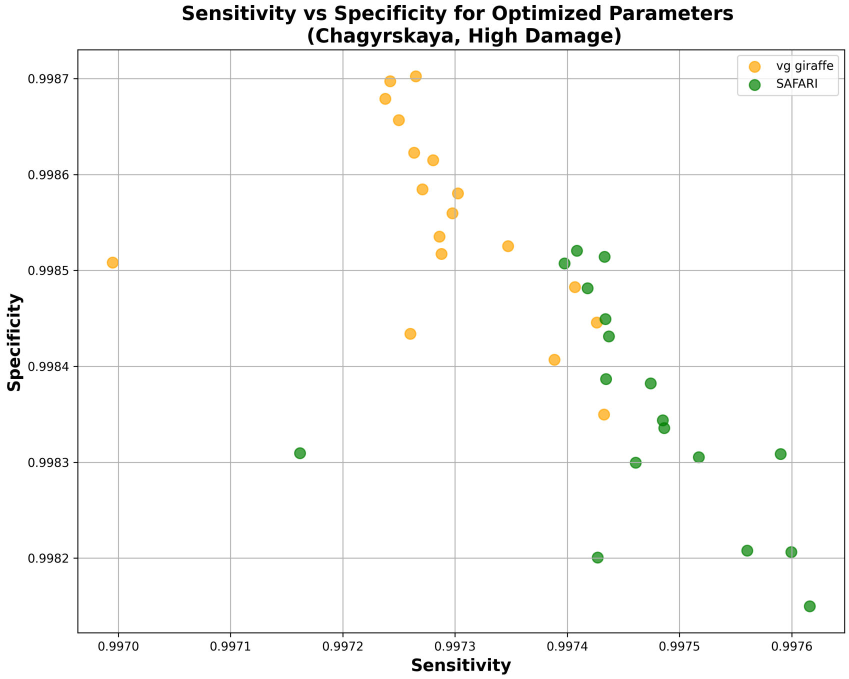

(a) High damage

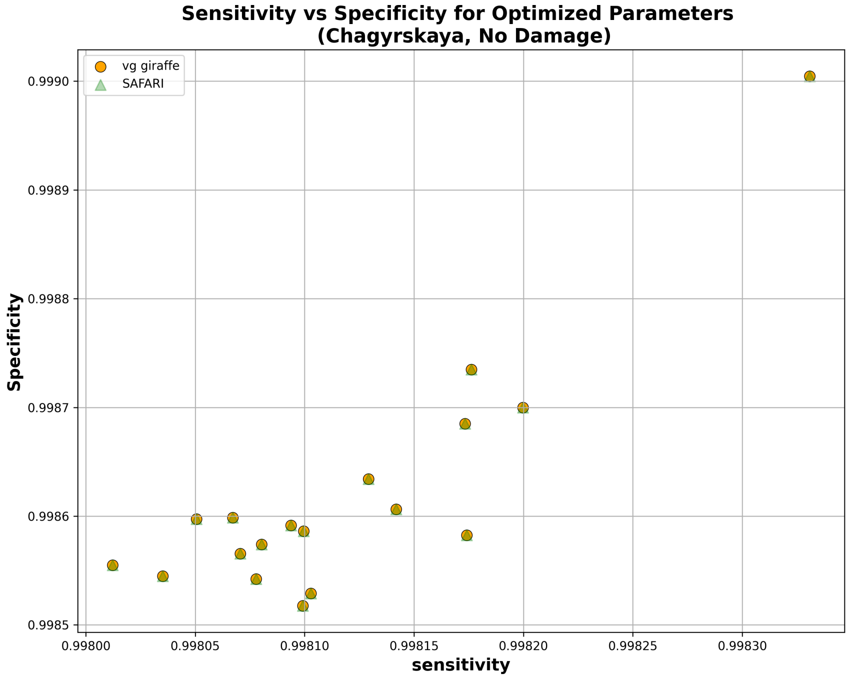

(b) Zero damage (no difference)

Figure 6: Sensitivity vs. specificity comparison of `SAFARI` and `vg giraffe` on surjected linear alignments using the empirically derived fragment length distribution from Chagyrskaya Cave, Siberia. The $(k, w)$ indexing parameters were individually selected to optimize median sensitivity across samples, out of all tested parameter pairs. (a) Samples with high levels of damage. Here we can see that sensitivity has clearly improved with `SAFARI` thanks to the recovered alignments, at the expense of a drop in specificity. This is almost always a beneficial tradeoff in the context of aDNA analysis. (b) Samples with zero damage. As expected, there is no difference between `SAFARI` and `vg giraffe` alignments, as the resulting alignments are identical thus showing that our strategy does not generate spurious alignment in the absence of deamination.

##### G.3.3 Median $F_1$ Scores for Pangenome Aligners Using Default Indexing Parameter Values, Chagyrskaya Cave Distribution

Figure 7 shows median $F_1$ score for the Chagyrskaya Cave distribution dataset for each of the two pangenome aligners, stratified by damage. It is clear that by rescuing deaminated seeds, `SAFARI` improves overall alignment performance, especially on highly damaged samples.

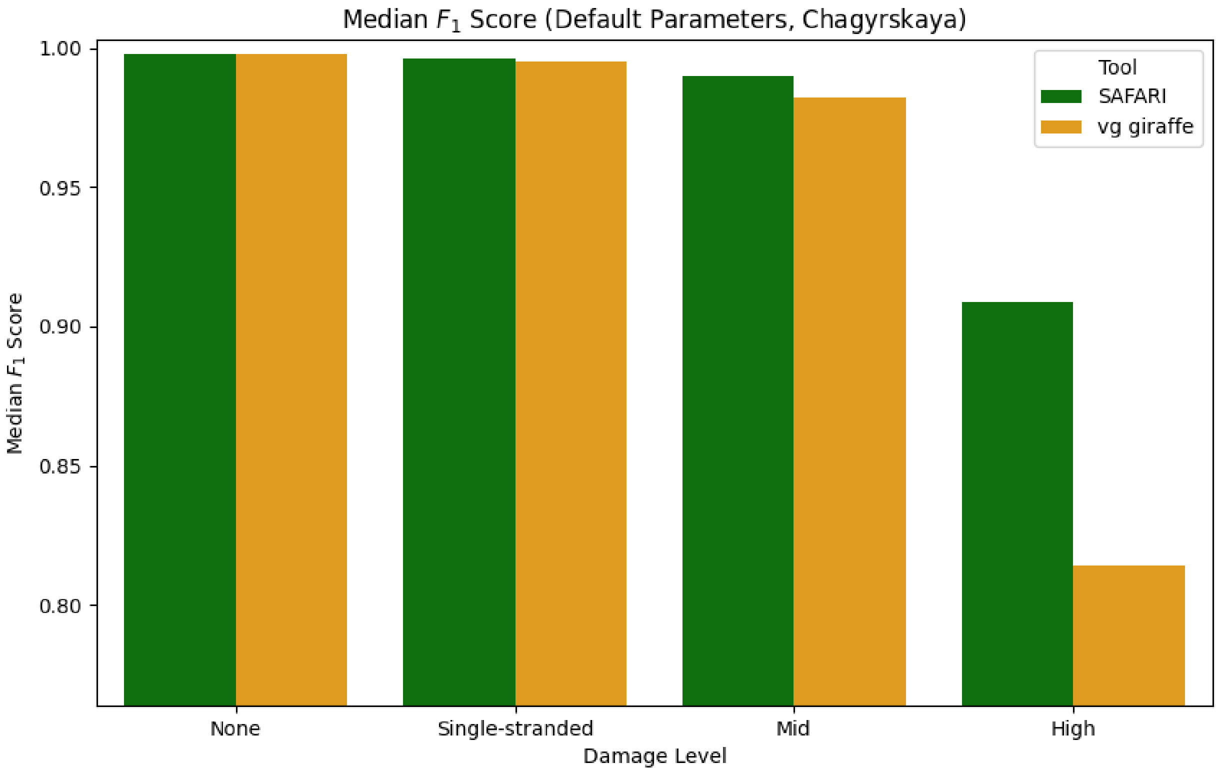

Figure 7: Median $F_1$ score of each of the two pangenomic alignment tools stratified by level of aDNA damage under default indexing parameters $(k, w) = (29, 11)$ for the empirically derived fragment length distribution from Chagyrskaya Cave, Croatia. The improvement in median $F_1$ score conferred by the deaminated seed rescue in `SAFARI` is especially clear at high levels of damage.

###### G.3.4 Median $F_1$ Scores for Pangenome Aligners Using Sensitivity-optimized Indexing Parameter Values

Figures 8a and 8b show median $F_1$ scores for `vg giraffe` vs. `SAFARI` on the Chagyrskaya Cave and Vindija Cave empirically-derived fragment length distribution datasets, respectively, using $k, w$ indexing parameter values optimized for maximum sensitivity.

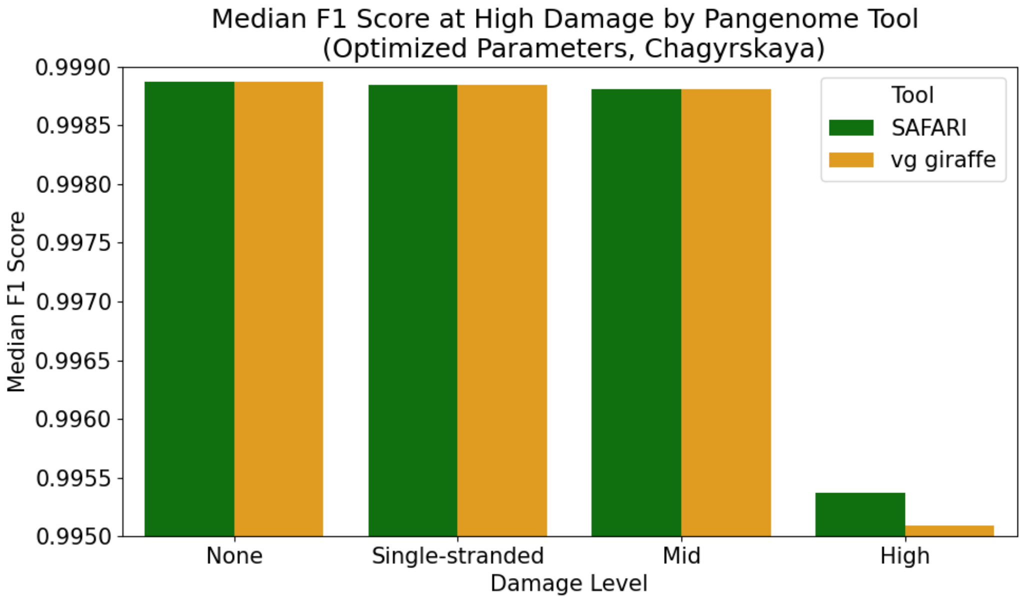

(a) Chagyrskaya Cave

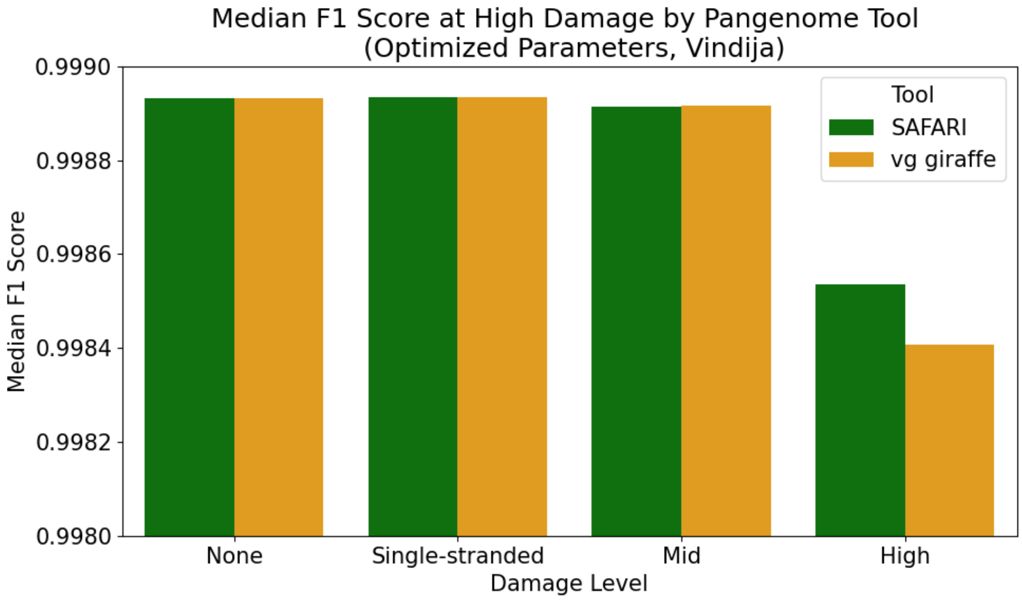

(b) Vindija Cave

Figure 8: Median $F_1$ scores across samples by pangenome aligner when using sensitivity-optimized $k, w$ indexing parameter values. The optimized parameters were $(k, w) = (10, 2)$ for both tools in both cases. We show both *in silico* simulation datasets corresponding to each of the two empirically-derived fragment length distributions.

##### G.3.5 Differences in Median Sensitivity/Specificity Between Pangenome Aligners By Indexing Parameter Values

It is interesting to explore how altering the $(k, w)$ indexing parameter values - that is to say the kmer size and density of the minimizer and (in the case of `SAFARI`) RYmer indexes affects the aligner's behavior in terms of where it falls along the tradeoff between sensitivity and specificity. Below (Figures 9 and 10) we plot differences in these metrics between tools by parameter values for both datasets of samples obeying empirically-derived fragment length distributions.

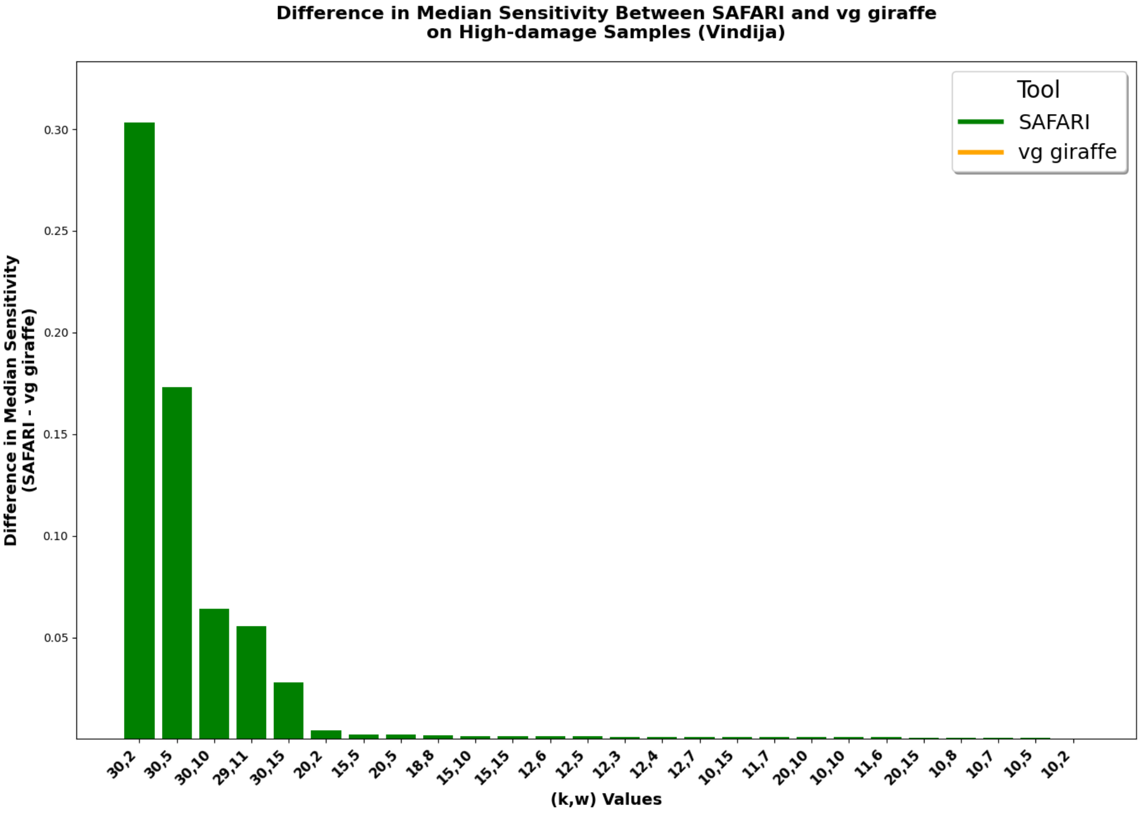

(a) Difference in median sensitivity between `SAFARI` and `vg giraffe` by indexing parameter values.

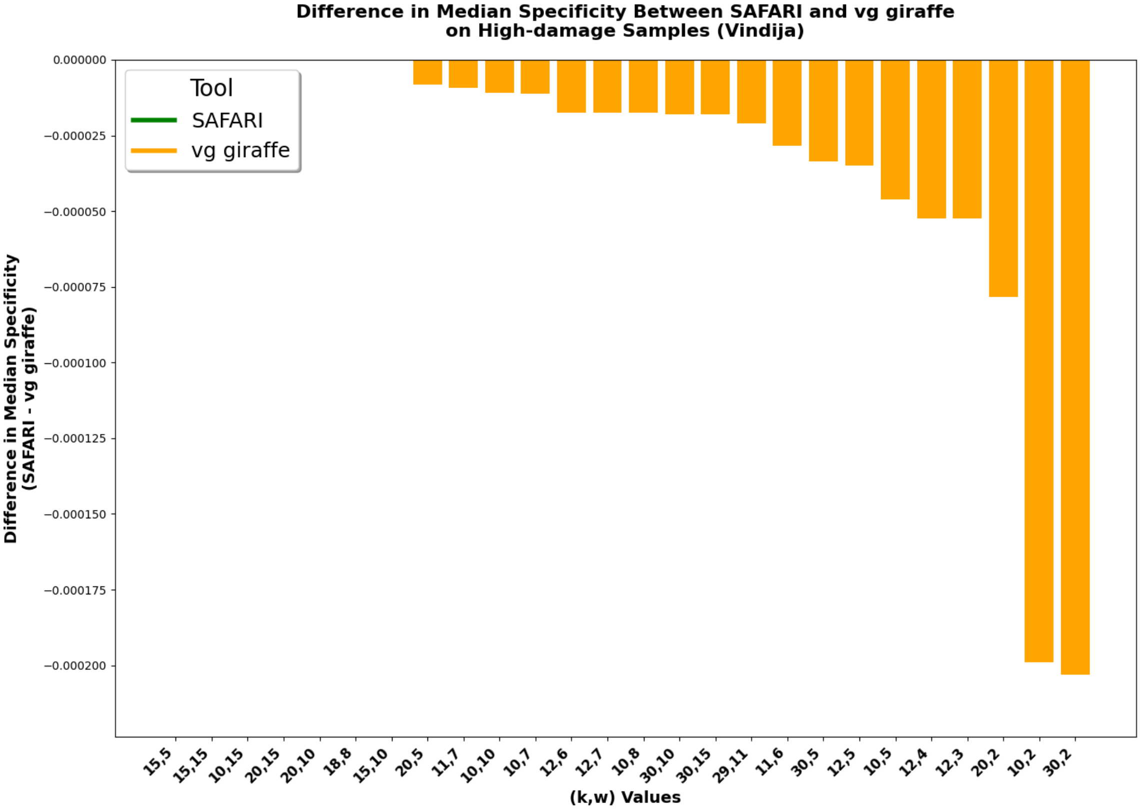

(b) Difference in median specificity between `SAFARI` and `vg giraffe` by indexing parameter values.

Figure 9: Difference in median sensitivity and median specificity between `SAFARI` and `vg giraffe` by indexing parameter values.

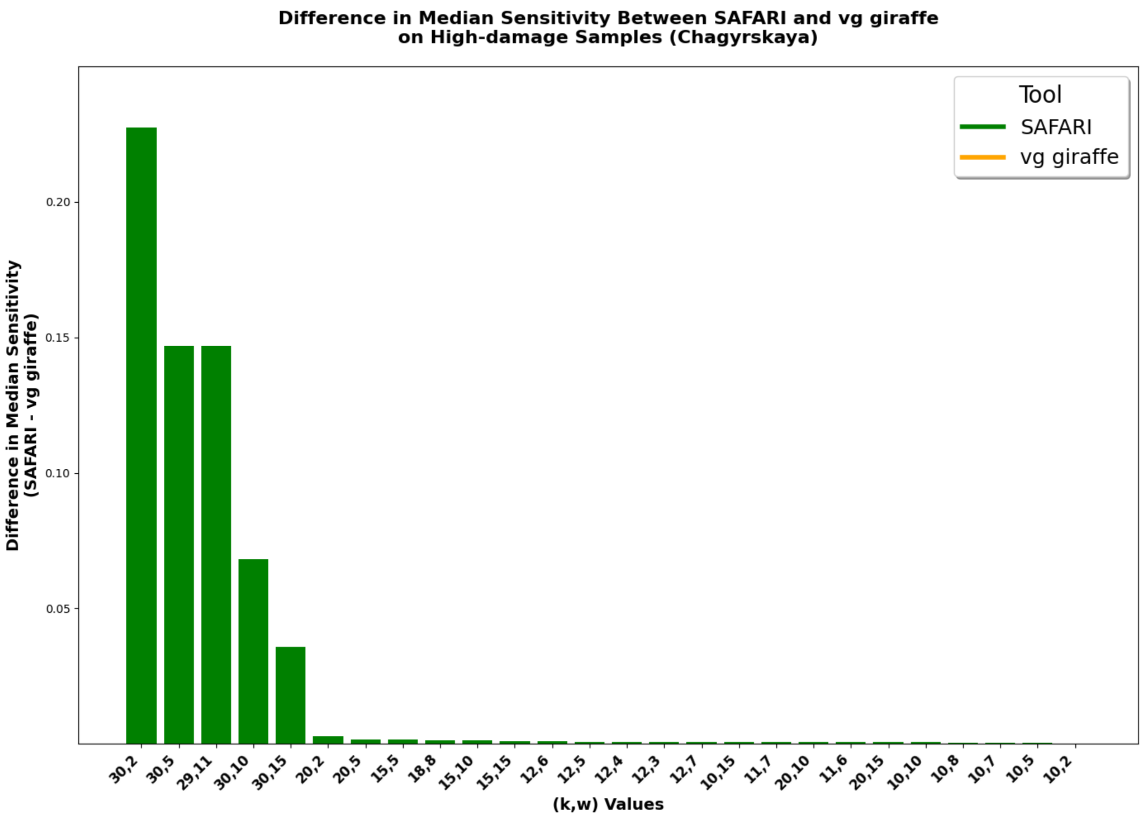

(a) Difference in median sensitivity between SAFARI and vg giraffe by indexing parameter values.

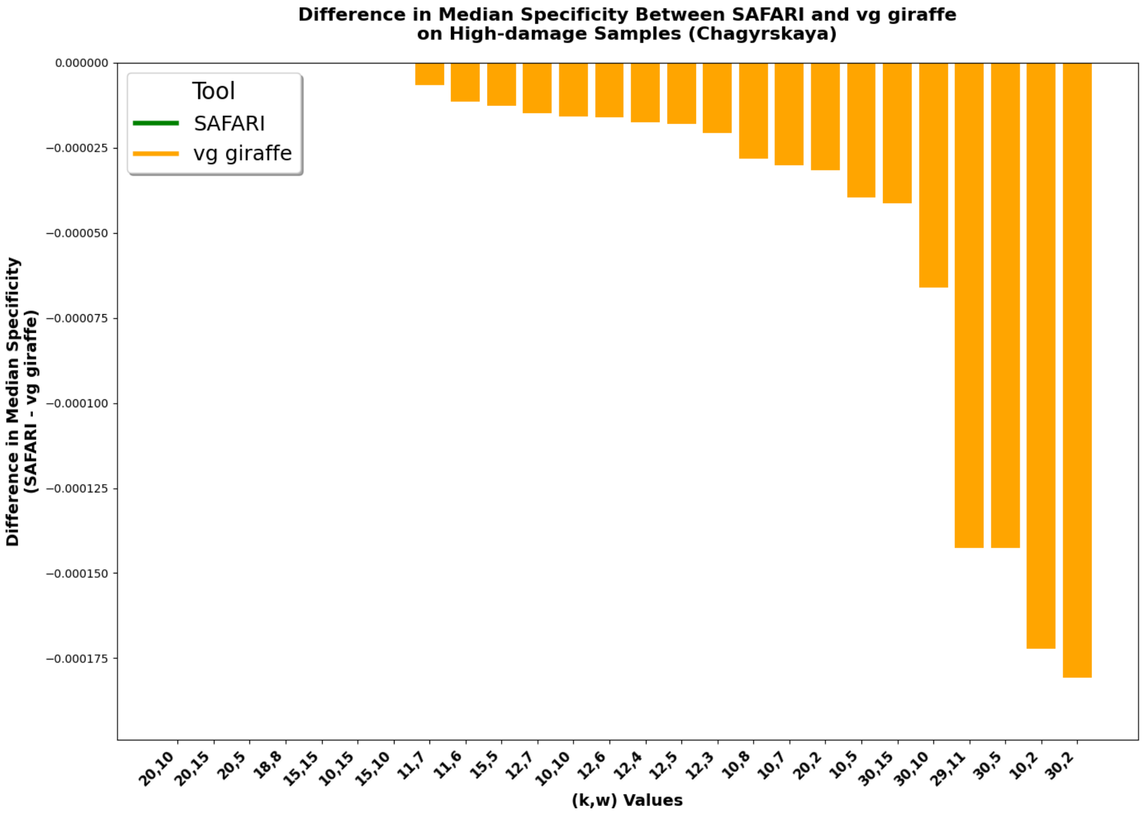

(b) Difference in median specificity between SAFARI and vg giraffe by indexing parameter values.

Figure 10: Difference in median sensitivity and median specificity between SAFARI and vg giraffe by indexing parameter values.

##### G.3.6 Median $F_1$ Scores Across All Tools Using Chagyrskaya Cave Fragment Length Distribution

Figure 11 plots median $F_1$ score for all tools on the Chagyrskaya Cave fragment length distribution dataset for the surjected linear alignment experiment. Similar to that of the Vindija Cave distribution we see that the two pangenome tools `vg giraffe` and `SAFARI`, in addition to `SHRiMP`, are the most robust to high levels of damage.

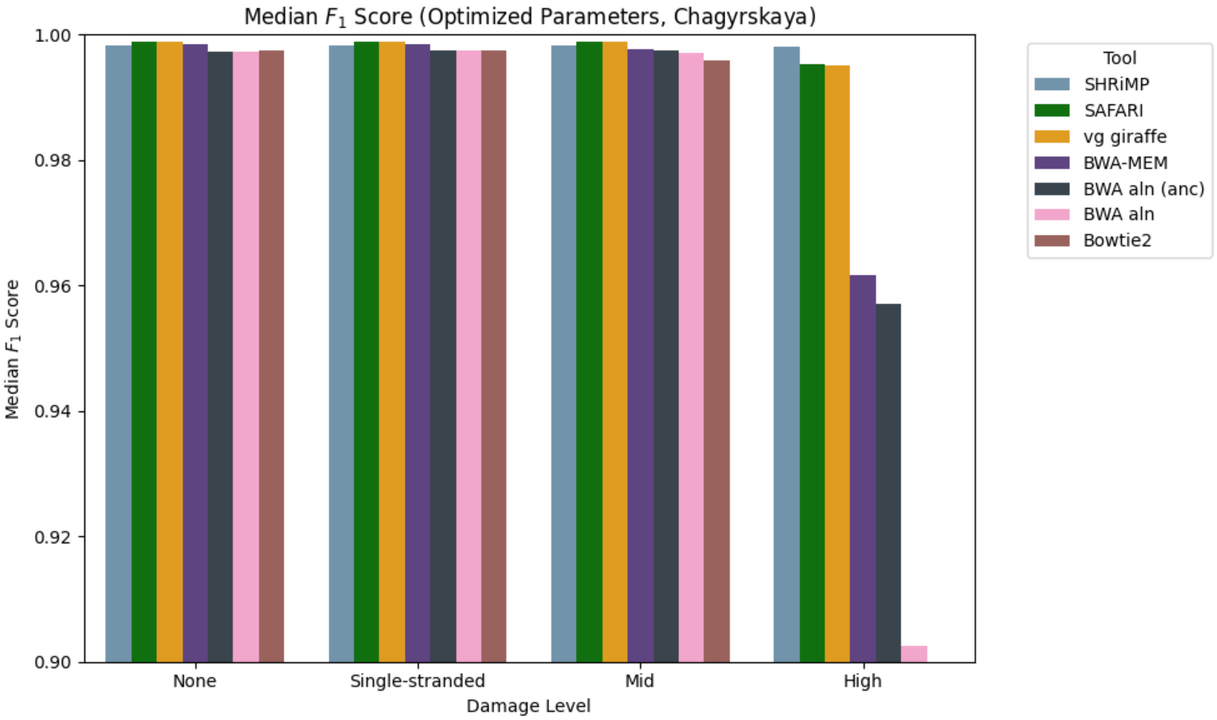

Figure 11: Median $F_1$ score across samples following the fragment length distribution from Chagyrskaya Cave, Siberia, stratified by level of damage. In the case of `SAFARI` and `vg giraffe` we use indexing parameter values that optimize for maximum sensitivity. We see that many tools worsen in performance precipitously once damage reaches a high level, with the two pangenome tools and `SHRiMP` being the three that are relatively robust to high damage overall.

##### G.3.7 Raw Counts for Pangenome Aligners on Chagyrskaya Cave Fragment Length Distribution Dataset

Reporting sensitivity, specificity, $F_1$, etc. is informative, but it is important to report the raw counts for the data as well. Below we provide raw counts for aligned reads by provenance (endogenous mitochondria DNA, NuMTs, or bacterial DNA) by pangenome aligner for the Chagyrskaya Cave empirical fragment length distribution *in silico* simulation dataset.

Table 11: Raw counts of mapped reads by provenance in the surjected linear alignment experiment for the dataset of `in silico` simulations following the empirically-derived fragment length distribution from Chagyrskaya Cave, Siberia, at high-level damage rates. *Bact* stands for number of mapped bacterial reads. *NuMT* is the number of mapped NuMT reads. *Mito* is the number of correctly mapped mitochondrial reads. For readability only samples with subsampling rate 0.9 are shown.

| $k$ | $w$ | Bact | | NuMT | | Mito | |
|---|---|---|---|---|---|---|---|
| | | `SAF` | `gir` | `SAF` | `gir` | `SAF` | `gir` |
| 10 | 2 | 51/90235 | 36/90235 | 96/390 | 95/390 | 269814/270545 | 269769/270545 |
| 10 | 5 | 14/90235 | 10/90235 | 90/390 | 90/390 | 269644/270545 | 269574/270545 |
| 10 | 7 | 8/90235 | 7/90235 | 86/390 | 85/390 | 269257/270545 | 269158/270545 |
| 10 | 8 | 7/90235 | 6/90235 | 81/390 | 80/390 | 268941/270545 | 268834/270545 |
| 10 | 10 | 7/90235 | 6/90235 | 77/390 | 77/390 | 268141/270545 | 267990/270545 |
| 10 | 15 | 3/90235 | 3/90235 | 68/390 | 68/390 | 263509/270545 | 263311/270545 |
| 11 | 6 | 5/90235 | 5/90235 | 85/390 | 84/390 | 268997/270545 | 268841/270545 |
| 11 | 7 | 4/90235 | 4/90235 | 82/390 | 81/390 | 268653/270545 | 268490/270545 |
| 12 | 3 | 3/90235 | 3/90235 | 91/390 | 89/390 | 269371/270545 | 269165/270545 |
| 12 | 4 | 3/90235 | 3/90235 | 90/390 | 88/390 | 269064/270545 | 268852/270545 |
| 12 | 5 | 3/90235 | 3/90235 | 84/390 | 83/390 | 268729/270545 | 268516/270545 |
| 12 | 6 | 3/90235 | 3/90235 | 80/390 | 79/390 | 268314/270545 | 268107/270545 |
| 12 | 7 | 3/90235 | 3/90235 | 76/390 | 75/390 | 267759/270545 | 267560/270545 |
| 15 | 5 | 1/90235 | 1/90235 | 65/390 | 64/390 | 264671/270545 | 264263/270545 |
| 15 | 10 | 0/90235 | 0/90235 | 49/390 | 49/390 | 257044/270545 | 256730/270545 |
| 15 | 15 | 0/90235 | 0/90235 | 41/390 | 41/390 | 245827/270545 | 245582/270545 |
| 18 | 8 | 0/90235 | 0/90235 | 35/390 | 35/390 | 249089/270545 | 248726/270545 |
| 20 | 2 | 0/90235 | 0/90235 | 36/390 | 33/390 | 254092/270545 | 253326/270545 |
| 20 | 5 | 0/90235 | 0/90235 | 30/390 | 30/390 | 246975/270545 | 246532/270545 |
| 20 | 10 | 0/90235 | 0/90235 | 25/390 | 25/390 | 233162/270545 | 232977/270545 |
| 20 | 15 | 0/90235 | 0/90235 | 21/390 | 21/390 | 218578/270545 | 218427/270545 |
| 29 | 11 | 0/90235 | 0/90235 | 24/390 | 11/390 | 225364/270545 | 185539/270545 |
| 30 | 2 | 0/90235 | 0/90235 | 30/390 | 12/390 | 256899/270545 | 195465/270545 |
| 30 | 5 | 0/90235 | 0/90235 | 24/390 | 11/390 | 225364/270545 | 185539/270545 |
| 30 | 10 | 0/90235 | 0/90235 | 15/390 | 9/390 | 179719/270545 | 161369/270545 |
| 30 | 15 | 0/90235 | 0/90235 | 11/390 | 8/390 | 147130/270545 | 137522/270545 |

##### G.3.8 Damage Rate Estimation

The below table (Table 12) shows the root mean square distance between ground truth and predicted damage rate matrices from our linear alignment experiment across all tools, both pangenomic and linear. For `SAFARI` and `vg giraffe`, the reported values are over samples with the default indexing parameters $k = 29, w = 11$. Unfortunately for reasons we were unable to determine (but which fall outside the scope of this paper) the `bam2prof` estimates failed to provide meaningful profiles for alignments generated with `Bowtie2`, instead producing profiles containing "-nan" values. This is unfortunate, but the purpose here is merely to roughly show how the pangenome tools compare to extant aligners, and discarding `Bowtie2` for this particular experiment does not negate the improvement of `SAFARI` over `vg giraffe` which is the main point.

Table 12: Median RMSE by Aligner, Stratified by Damage Matrix and Fragment Length Distribution

| Fragment Length Dist | Damage Level | Aligner | Median RMSE |
|---|---|---|---|
| Chagyrskaya | None | ANC | 0.000254039314235 |
| Chagyrskaya | None | BBMap | 0.000216725954602 |
| Chagyrskaya | None | BWA aln | 0.000249590309279 |
| Chagyrskaya | None | BWA-MEM | 0.000245662099981 |
| Chagyrskaya | None | SAFARI | 0.000129801168650 |
| Chagyrskaya | None | SHRiMP | 0.000240953533138 |
| Chagyrskaya | None | vg giraffe | 0.000129801168650 |
| Chagyrskaya | Single-stranded | ANC | 0.000700009013038 |
| Chagyrskaya | Single-stranded | BBMap | 0.000662971080891 |
| Chagyrskaya | Single-stranded | BWA aln | 0.000682883702541 |
| Chagyrskaya | Single-stranded | BWA-MEM | 0.000374914755779 |
| Chagyrskaya | Single-stranded | SAFARI | 0.000781440569890 |
| Chagyrskaya | Single-stranded | SHRiMP | 0.000584915345703 |
| Chagyrskaya | Single-stranded | vg giraffe | 0.000782318258900 |
| Chagyrskaya | Mid | ANC | 0.001090007922072 |
| Chagyrskaya | Mid | BBMap | 0.001061515685155 |
| Chagyrskaya | Mid | BWA aln | 0.000796128304939 |
| Chagyrskaya | Mid | BWA-MEM | 0.002429977014382 |
| Chagyrskaya | Mid | SAFARI | 0.000621920167702 |
| Chagyrskaya | Mid | SHRiMP | 0.001011073756105 |
| Chagyrskaya | Mid | vg giraffe | 0.000625510373220 |
| Chagyrskaya | High | ANC | 0.020048194493544 |
| Chagyrskaya | High | BBMap | 0.001679988895036 |
| Chagyrskaya | High | BWA aln | 0.038637014215471 |
| Chagyrskaya | High | BWA-MEM | 0.040239909990277 |
| Chagyrskaya | High | SAFARI | 0.012330074318948 |
| Chagyrskaya | High | SHRiMP | 0.002421109681100 |
| Chagyrskaya | High | vg giraffe | 0.012478861693922 |
| Vindija | None | ANC | 0.000210624637848 |
| Vindija | None | BBMap | 0.000183636136553 |
| Vindija | None | BWA aln | 0.000203554096122 |
| Vindija | None | BWA-MEM | 0.000203369211957 |
| Vindija | None | SAFARI | 0.000115545385702 |
| Vindija | None | SHRiMP | 0.000249101805604 |
| Vindija | None | vg giraffe | 0.000115545385702 |
| Vindija | Single-stranded | ANC | 0.000634061586861 |
| Vindija | Single-stranded | BBMap | 0.000614000750481 |
| Vindija | Single-stranded | BWA aln | 0.000623136335772 |
| Vindija | Single-stranded | BWA-MEM | 0.000847970960142 |

Continued on next page

**Table 12 – continued from previous page**

| Fragment Length Dist | Damage Level | Aligner | Median RMSE |
|---|---|---|---|
| Vindija | Single-stranded | SAFARI | 0.000561150818737 |
| Vindija | Single-stranded | SHRiMP | 0.000639254455797 |
| Vindija | Single-stranded | vg giraffe | 0.000554107280269 |
| Vindija | Mid | ANC | 0.000742789194146 |
| Vindija | Mid | BBMap | 0.001124676539555 |
| Vindija | Mid | BWA aln | 0.000596203034631 |
| Vindija | Mid | BWA-MEM | 0.003097568199287 |
| Vindija | Mid | SAFARI | 0.000976162207646 |
| Vindija | Mid | SHRiMP | 0.000717332776321 |
| Vindija | Mid | vg giraffe | 0.000979666873226 |
| Vindija | High | ANC | 0.020780023588946 |
| Vindija | High | BBMap | 0.002682043168663 |
| Vindija | High | BWA aln | 0.036673222311489 |
| Vindija | High | BWA-MEM | 0.042771379640544 |
| Vindija | High | SAFARI | 0.008651229171550 |
| Vindija | High | SHRiMP | 0.003082623667806 |
| Vindija | High | vg giraffe | 0.008684805382982 |

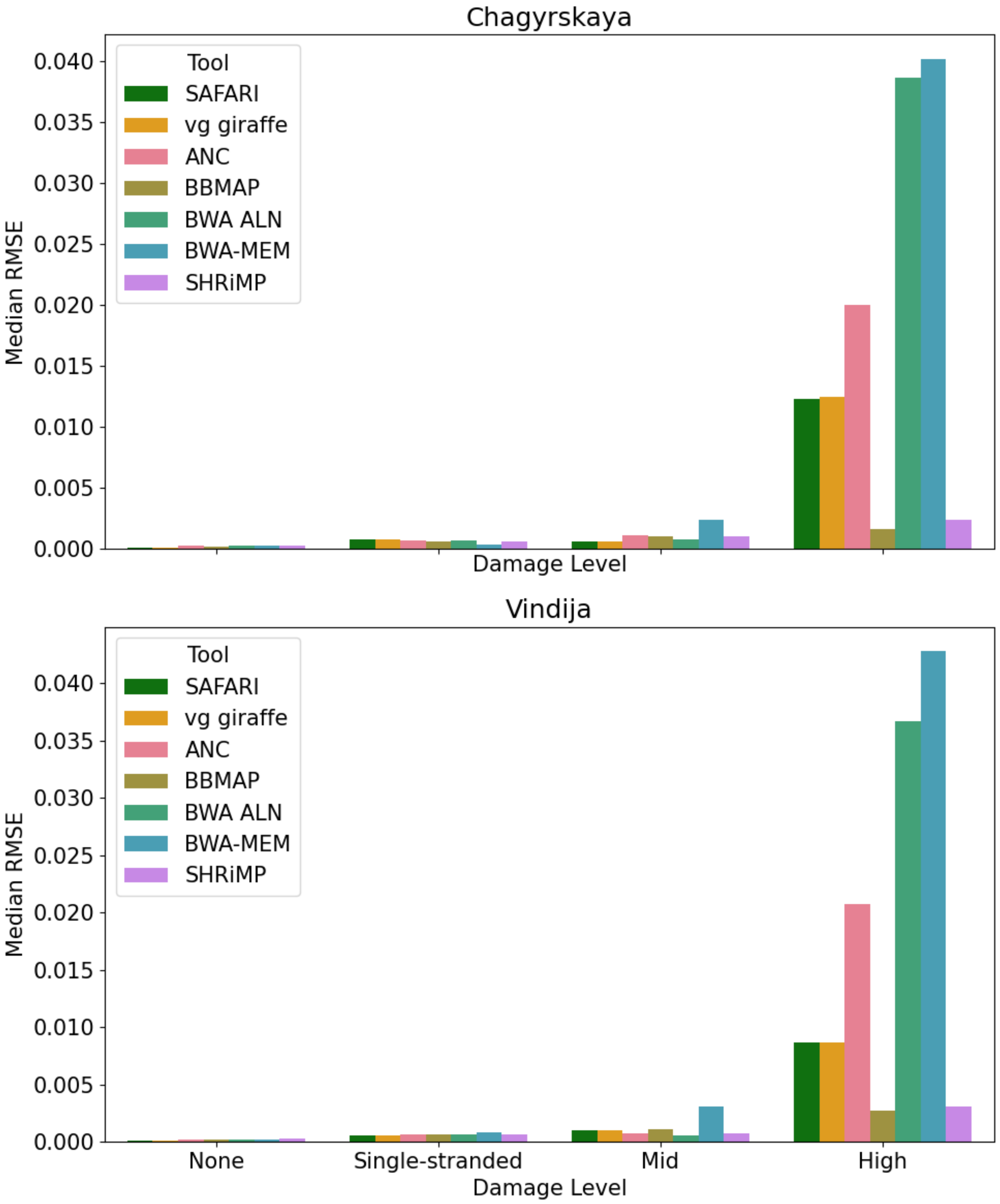

Figure 12: Median RMSE across samples for all alignment tools (both linear and pangenomic) in the surjected linear alignment experiment. **[TOP]** Samples following the empirical fragment length distribution from Chagyrskaya Cave. **[BOTTOM]** Samples following the empirical fragment length distribution from Vindija Cave. Note that the RMSE differences between `SAFARI` and `vg giraffe` are too minor to be discernible from this figure.

**Damage Estimate Visualization on a High-damage Sample for Both Pangenome Aligners** Although there is clear improvement, the damage estimates overall remain very similar. To illustrate how similar the damage estimates from both tools are, even for highly damaged samples, we show in Figure 13 the `bam2prof` damage rate estimates from the same high-damage input sample using both `vg giraffe` [**TOP**] and `SAFARI` [**BOTTOM**].

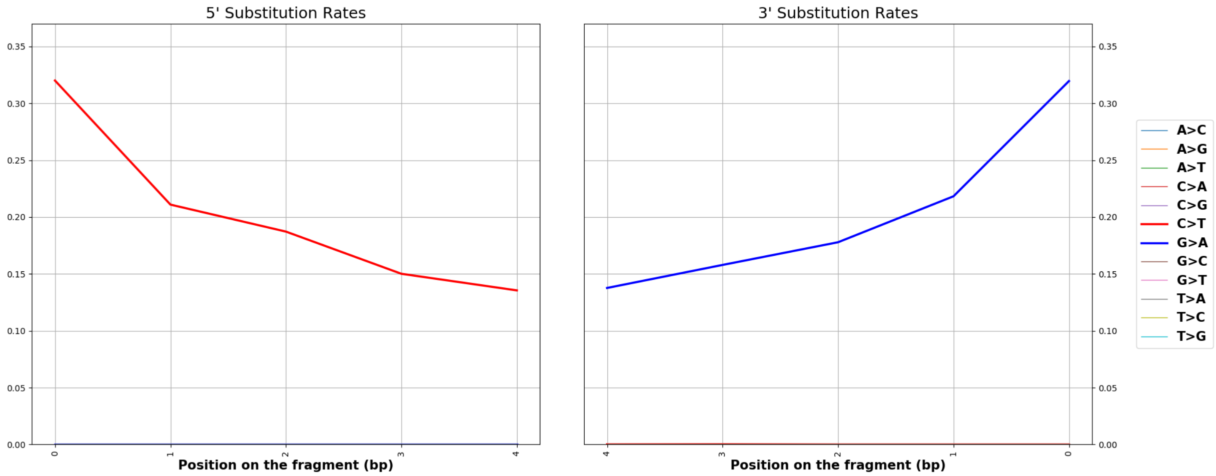

(a) `vg giraffe`

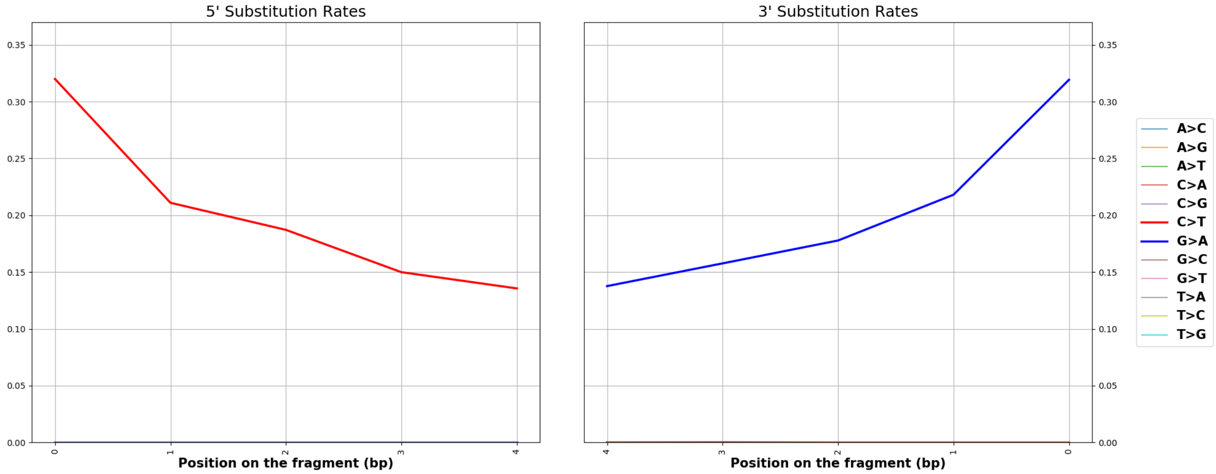

(b) `SAFARI`

Figure 13: Deamination patterns for one high-damage sample in our surjected linear alignment experiment after alignment with `vg giraffe` [**TOP**] and `SAFARI` [**BOTTOM**] obeying the empirically-derived fragment length distribution from Chagyrskaya Cave. The estimates are very nearly (though not entirely) identical.

### H Pseudocode

In this section we provide some brief pseudocode to supplement our description of the RYmer-specific modification to the `vg giraffe` codebase. Of course, for full details it is best to peruse the source code itself as found in our data repository on Github.

#### H.1 Converting Kmer Keys to RYmer Keys

Below we demonstrate the method for converting a kmer-encoded key to its unique RYmer-encoded representation.

**Algorithm 1** Kmer to RYmer Conversion

```
procedure ConvertKmerToRYmer(kmerKey, kmerLength)
    RYmerKey ← new Key()                                ▷ Initialize an empty RYmerKey
    for i ← 0 to kmerLength − 1 do
        baseType ← GetBaseType(kmerKey, i)              ▷ Determine if purine or pyrimidine
        if baseType = purine then
            RYmerKey ← RYmerKey ∘ 0                     ▷ Append 0 for purine (A or G)
        else
            RYmerKey ← RYmerKey ∘ 1                     ▷ Append 1 for pyrimidine (C or T)
    return RYmerKey
```

#### H.2 Filtering RYmers

Below is a thumbnail sketch of the Bayesian filter for putative Rymers described in the main manuscript. For each seed the posterior probability of genuine alignment vs. spurious alignment is computed, and putative RYmers which do not exceed the predefined threshold are discarded before clustering and extension of seeds even begin.

**Algorithm 2** Filter RYmers

```
procedure FilterAndProcessRYmers
    for RYmer ∈ RYmers do
        probModel1 ← P(RYmer | model 1)
        probModel2 ← P(RYmer | model 2)
        priorModel1 ← P(model 1)
        priorModel2 ← P(model 2)
        posteriorModel1 ← (probModel1×priorModel1) / (probModel1×priorModel1+probModel2×priorModel2)
        if posteriorModel1 > threshold then
            minimizerIndexOffset ← RYmer payload
            minimizer ← minimizerIndex[minimizerIndexOffset]
            Keep minimizer
        else
            Remove RYmer
```

### I Experiment Memory and Runtime

Tables 13 and 14 provide memory and runtime details for our experiments. All such data are obtained with the Linux command `/usr/bin/time -v`, except for the linear surjection experiment, which was benchmarking internally using `Snakemake`. Table 15 provides runtime information for mapping to the full hominin panmitogenome for varying $(k, w)$ parameter pairs.

#### I.1 `HaploCart` and `euka`

Table 13: Comparison of runtime and memory consumption between `SAFARI` for the `HaploCart` and `euka` experiments. During these experiments some other light processes were being run on the same server node which may introduce a small amount of added noise.

| | | **SAFARI** | **vg giraffe** |
|---|---|---|---|
| `HaploCart` | User time (s) | 10897.03 | 10453.30 |
| | System time (s) | 1115.41 | 1000.11 |
| | Wall clock time (h:mm:ss) | 18:39.25 | 8:14.21 |
| | Maximum resident set size (Kb) | 1879608 | 1335592 |
| `euka` | User time (s) | 18.49 | 19.64 |
| | System time (s) | 23.37 | 29.70 |
| | Wall clock time (h:mm:ss) | 1:06.55 | 0:55.93 |
| | Maximum resident set size (Kb) | 46053840 | 46054324 |

#### I.2 Direct Comparison (`SAFARI` vs. `vg giraffe`)

Table 14: Comparison of key performance metrics between `SAFARI` and `vg giraffe` as measured by `Snakemake` internal benchmarks.

| Metric | `SAFARI` | `vg giraffe` |
|---|---|---|
| s | 165.446 | **156.658** |
| max RSS | 615.659 | **359.881** |
| Mean load | 144.140 | **137.505** |

#### I.3 Runtime as a Function of Minimizer/RYmer Index Values

Table 15: Runtime for `SAFARI` Mapping to the Full Hominin Panmitogenome at Varying Minimizer/Rymer Index Parameters

| $k$ | $w$ | **User Time (s)** | **System Time (s)** | **Elapsed Time (h:mm:ss)** |
|---|---|---|---|---|
| 10 | 2 | 4433.88 | 23.76 | 1:45.21 |
| 12 | 3 | 3438.49 | 20.41 | 1:13.80 |
| 15 | 5 | 2575.69 | 16.08 | 0:52.76 |
| 17 | 10 | 1639.84 | 13.65 | 0:47.92 |
| 20 | 10 | 1649.12 | 13.92 | 0:48.89 |
| 20 | 30 | 791.52 | 7.74 | 0:22.19 |
| 29 | 11 | 1225.73 | 7.61 | 0:23.17 |
| 29 | 30 | 625.39 | 3.79 | 0:12.29 |

### J Experiment Data Details

#### J.1 Nucleotide Misincorporation Matrices for Simulated Data

Nucleotide misincorporation matrices for high rates of damage are taken from [8]. Matrices for medium rates of damage are taken from [24]. Matrices for single-stranded rates of damage are taken from [22]. ATP2 matrices from a Denisovan sample (used for the experiments in Section D.5 and D.6) are taken from [31].

Table 16: Zero damage nucleotide misincorporation rate matrix for the 3' end.

| | A | | | C | | | G | | | T | | |
|---|---|---|---|---|---|---|---|---|---|---|---|---|
| | →C | →G | →T | →A | →G | →T | →A | →C | →T | →A | →C | →G |
| -4 | 0 | 0 | 0 | 0 | 0 | 0 | 0 | 0 | 0 | 0 | 0 | 0 |
| -3 | 0 | 0 | 0 | 0 | 0 | 0 | 0 | 0 | 0 | 0 | 0 | 0 |
| -2 | 0 | 0 | 0 | 0 | 0 | 0 | 0 | 0 | 0 | 0 | 0 | 0 |
| -1 | 0 | 0 | 0 | 0 | 0 | 0 | 0 | 0 | 0 | 0 | 0 | 0 |
| 0 | 0 | 0 | 0 | 0 | 0 | 0 | 0 | 0 | 0 | 0 | 0 | 0 |

Table 17: Zero damage nucleotide misincorporation rate matrix for the 5' end.

| | A | | | C | | | G | | | T | | |
|---|---|---|---|---|---|---|---|---|---|---|---|---|
| | →C | →G | →T | →A | →G | →T | →A | →C | →T | →A | →C | →G |
| 0 | 0 | 0 | 0 | 0 | 0 | 0 | 0 | 0 | 0 | 0 | 0 | 0 |
| 1 | 0 | 0 | 0 | 0 | 0 | 0 | 0 | 0 | 0 | 0 | 0 | 0 |
| 2 | 0 | 0 | 0 | 0 | 0 | 0 | 0 | 0 | 0 | 0 | 0 | 0 |
| 3 | 0 | 0 | 0 | 0 | 0 | 0 | 0 | 0 | 0 | 0 | 0 | 0 |
| 4 | 0 | 0 | 0 | 0 | 0 | 0 | 0 | 0 | 0 | 0 | 0 | 0 |

Table 18: Double-stranded medium damage nucleotide misincorporation rate matrix for the 3' end.

| | A | | | C | | | G | | | T | | |
|---|---|---|---|---|---|---|---|---|---|---|---|---|
| | →C | →G | →T | →A | →G | →T | →A | →C | →T | →A | →C | →G |
| -4 | 0 | 0 | 0 | 0 | 0 | 0 | 0.0160588 | 0 | 0 | 0 | 0 | 0 |
| -3 | 0 | 0 | 0 | 0 | 0 | 0 | 0.0171117 | 0 | 0 | 0 | 0 | 0 |
| -2 | 0 | 0 | 0 | 0 | 0 | 0 | 0.0260759 | 0 | 0 | 0 | 0 | 0 |
| -1 | 0 | 0 | 0 | 0 | 0 | 0 | 0.0483092 | 0 | 0 | 0 | 0 | 0 |
| 0 | 0 | 0 | 0 | 0 | 0 | 0 | 0.13679 | 0 | 0 | 0 | 0 | 0 |

Table 19: Double-stranded medium damage nucleotide misincorporation rate matrix for the 5' end.

| | A | | | C | | | G | | | T | | |
|---|---|---|---|---|---|---|---|---|---|---|---|---|
| | →C | →G | →T | →A | →G | →T | →A | →C | →T | →A | →C | →G |
| 0 | 0 | 0 | 0 | 0 | 0 | 0.173982 | 0 | 0 | 0 | 0 | 0 | 0 |
| 1 | 0 | 0 | 0 | 0 | 0 | 0.0703105 | 0 | 0 | 0 | 0 | 0 | 0 |
| 2 | 0 | 0 | 0 | 0 | 0 | 0.0259287 | 0 | 0 | 0 | 0 | 0 | 0 |
| 3 | 0 | 0 | 0 | 0 | 0 | 0.0145445 | 0 | 0 | 0 | 0 | 0 | 0 |
| 4 | 0 | 0 | 0 | 0 | 0 | 0.0135411 | 0 | 0 | 0 | 0 | 0 | 0 |

Table 20: Double-stranded high damage nucleotide misincorporation rate matrix for the 3' end.

| | A | | | C | | | G | | | T | | |
|---|---|---|---|---|---|---|---|---|---|---|---|---|
| | →C | →G | →T | →A | →G | →T | →A | →C | →T | →A | →C | →G |
| -4 | 0 | 0 | 0 | 0 | 0 | 0 | 0.146352 | 0 | 0 | 0 | 0 | 0 |
| -3 | 0 | 0 | 0 | 0 | 0 | 0 | 0.164419 | 0 | 0 | 0 | 0 | 0 |
| -2 | 0 | 0 | 0 | 0 | 0 | 0 | 0.188599 | 0 | 0 | 0 | 0 | 0 |
| -1 | 0 | 0 | 0 | 0 | 0 | 0 | 0.223405 | 0 | 0 | 0 | 0 | 0 |
| 0 | 0 | 0 | 0 | 0 | 0 | 0 | 0.32891 | 0 | 0 | 0 | 0 | 0 |

Table 21: Double-stranded high damage nucleotide misincorporation rate matrix for the 5' end.

| | A | | | C | | | G | | | T | | |
|---|---|---|---|---|---|---|---|---|---|---|---|---|
| | →C | →G | →T | →A | →G | →T | →A | →C | →T | →A | →C | →G |
| 0 | 0 | 0 | 0 | 0 | 0 | 0.329405 | 0 | 0 | 0 | 0 | 0 | 0 |
| 1 | 0 | 0 | 0 | 0 | 0 | 0.221745 | 0 | 0 | 0 | 0 | 0 | 0 |
| 2 | 0 | 0 | 0 | 0 | 0 | 0.187678 | 0 | 0 | 0 | 0 | 0 | 0 |
| 3 | 0 | 0 | 0 | 0 | 0 | 0.161196 | 0 | 0 | 0 | 0 | 0 | 0 |
| 4 | 0 | 0 | 0 | 0 | 0 | 0.144011 | 0 | 0 | 0 | 0 | 0 | 0 |

Table 22: Single-stranded low damage nucleotide misincorporation rate matrix for the 3' end.

| | A | | | C | | | G | | | T | | |
|---|---|---|---|---|---|---|---|---|---|---|---|---|
| | →C | →G | →T | →A | →G | →T | →A | →C | →T | →A | →C | →G |
| -4 | 0 | 0 | 0 | 0 | 0 | 0.00242025 | 0 | 0 | 0 | 0 | 0 | 0 |
| -3 | 0 | 0 | 0 | 0 | 0 | 0.0035038 | 0 | 0 | 0 | 0 | 0 | 0 |
| -2 | 0 | 0 | 0 | 0 | 0 | 0.00400941 | 0 | 0 | 0 | 0 | 0 | 0 |
| -1 | 0 | 0 | 0 | 0 | 0 | 0.0277661 | 0 | 0 | 0 | 0 | 0 | 0 |
| 0 | 0 | 0 | 0 | 0 | 0 | 0.074432 | 0 | 0 | 0 | 0 | 0 | 0 |

Table 23: Single-stranded low damage nucleotide misincorporation rate matrix for the 5' end.

| | A | | | C | | | G | | | T | | |
|---|---|---|---|---|---|---|---|---|---|---|---|---|
| | →C | →G | →T | →A | →G | →T | →A | →C | →T | →A | →C | →G |
| 0 | 0 | 0 | 0 | 0 | 0 | 0.0329203 | 0 | 0 | 0 | 0 | 0 | 0 |
| 1 | 0 | 0 | 0 | 0 | 0 | 0.00318909 | 0 | 0 | 0 | 0 | 0 | 0 |
| 2 | 0 | 0 | 0 | 0 | 0 | 0.00279628 | 0 | 0 | 0 | 0 | 0 | 0 |
| 3 | 0 | 0 | 0 | 0 | 0 | 0.00222379 | 0 | 0 | 0 | 0 | 0 | 0 |
| 4 | 0 | 0 | 0 | 0 | 0 | 0.00274437 | 0 | 0 | 0 | 0 | 0 | 0 |

Table 24: ATP2 damage nucleotide misincorporation rate matrix for the 3' end.

| | A | | | C | | | G | | | T | | |
|---|---|---|---|---|---|---|---|---|---|---|---|---|
| | →C | →G | →T | →A | →G | →T | →A | →C | →T | →A | →C | →G |
| -4 | 0 | 0 | 0 | 0 | 0 | 0 | 0.32891 | 0 | 0 | 0 | 0 | 0 |
| -3 | 0 | 0 | 0 | 0 | 0 | 0 | 0.223405 | 0 | 0 | 0 | 0 | 0 |
| -2 | 0 | 0 | 0 | 0 | 0 | 0 | 0.188599 | 0 | 0 | 0 | 0 | 0 |
| -1 | 0 | 0 | 0 | 0 | 0 | 0 | 0.164419 | 0 | 0 | 0 | 0 | 0 |
| 0 | 0 | 0 | 0 | 0 | 0 | 0 | 0.146352 | 0 | 0 | 0 | 0 | 0 |

Table 25: ATP2 damage nucleotide misincorporation rate matrix for the 5' end.

| | A | | | C | | | G | | | T | | |
|---|---|---|---|---|---|---|---|---|---|---|---|---|
| | →C | →G | →T | →A | →G | →T | →A | →C | →T | →A | →C | →G |
| 0 | 0 | 0 | 0 | 0 | 0 | 0.329405 | 0 | 0 | 0 | 0 | 0 | 0 |
| 1 | 0 | 0 | 0 | 0 | 0 | 0.221745 | 0 | 0 | 0 | 0 | 0 | 0 |
| 2 | 0 | 0 | 0 | 0 | 0 | 0.187678 | 0 | 0 | 0 | 0 | 0 | 0 |
| 3 | 0 | 0 | 0 | 0 | 0 | 0.161196 | 0 | 0 | 0 | 0 | 0 | 0 |
| 4 | 0 | 0 | 0 | 0 | 0 | 0.144011 | 0 | 0 | 0 | 0 | 0 | 0 |

##### J.2 Downsampled Ancient BAM Files for `HaploCart` Experiment

The forty BAM files used in the `HaploCart` experiment are publicly available from the European Nucleotide Archive (ENA)[15]. Accession numbers are provided in [29], S1 file Section 2.7.

##### J.3 Simulated Metagenomic Environment for `euka` Experiment

We generated a simplified metagenomic environment by modifying the published `Snakefile` emulating a Late Pleistocene cave environment from Mexico (`https://github.com/nicolaavogel/eukaPaperData/tree/master/CreateSimulatedEnv/Snakefile_cave_env`). A detailed discussion of the curated and processed data for this environment is provided in [33], Supplementary Material Section 8. For the purposes of our `SAFARI` benchmarking experiment we somewhat simplified this environment. The modified `Snakefile` can be found at `https://github.com/JoshuaDanielRubin/SAFARI_paper_data/blob/main/cave_env_Snakefile`.

In our environment we included three eukaryotic taxa, namely *Ursidae* (bears), *Sciuridae* (squirrels), and *Falconiformes* (falcons and related birds). 5000 fragments of each taxon were present in the environment. Also present were 40000 viral fragments, 80000 archaeal fragments, 80000 bacterial fragments, 10000 plantal fragments, and 60000 fungal fragments. In addition 25000 nuclear fragments from each of the present eukaryotic taxa were incorporated. aDNA damage was simulated using the `deamSim` subcommand of `gargammel` using the high damage rate matrices provided in Section J.1[28].
